# Social hosts evade predation but have deadlier parasites

**DOI:** 10.1101/2021.09.10.459661

**Authors:** Jason C. Walsman, Mary J. Janecka, David R. Clark, Rachael D. Kramp, Faith Rovenolt, Regina Patrick, Ryan S. Mohammed, Mateusz Konczal, Clayton E. Cressler, Jessica F. Stephenson

## Abstract

Parasites exploit hosts to replicate and transmit, but overexploitation kills host and parasite (*1*): predators may shift this cost-benefit balance by consuming hosts (*2–4*) or changing host behavior, but the strength of these effects remains unclear. Modeling both, we find a primary, strong effect: hosts group to defend against predators (*5*), increasing parasite transmission, thus multiple infections, and therefore favoring more exploitative, virulent, parasites (*6*). Indeed, among 18 Trinidadian *Gyrodactyus* spp. parasite lines, those collected from high predation guppy populations were more virulent in common garden than those from low predation populations. Our model accurately predicted this result when parametrized with our experimentally demonstrated virulence-transmission trade-off, implicating the behavioral effects of predation. Broadly, our results indicate that reduced social contact selects against parasite virulence.

**One-Sentence Summary:** Our theory and data show predators cause increased host social grouping; the resulting transmission favors parasite virulence.

## Main text

Unprecedented infectious disease emergence among human and wildlife populations demands that we improve our understanding of the evolutionary ecology of parasites and pathogens (hereafter ‘parasites’). A key tenet of the evolutionary theory of infectious disease is that parasites face a trade-off between virulence and transmission (*8, 9*). Parasites exploit their hosts to replicate and transmit but increasing this exploitation harms parasite fitness by killing hosts (‘virulence’), which reduces the time window for successful transmission, or the ‘infectious period’ (*1*). With appropriate curvature, this trade-off should result in stabilizing selection for intermediate virulence (*10–13*). While theory predicts various mechanisms can shift the costs and benefits of exploitation, altering virulence evolution (*2–4, 6*) and host disease outcomes (*9, 12, 14, 15*), empirical evaluation of their relative importance remains scant (*9, 16, 17*).

Predation, a ubiquitous ecological interaction, can theoretically select for higher or lower virulence (*2, 4, 9*) through multiple pathways (we highlight four in Fig. 1). Predation directly increases host mortality, selecting for faster exploitation and transmission before the host dies, and hence increased virulence (pathway 1 in Fig. 1; *2, 3, 17*). Conversely, predation selects for lower virulence by reducing the density of infected hosts, reducing the per-host rate at which new infections arise (‘force of infection’); lower force of infection reduces infections with multiple parasite genotypes (‘multiple infections’) and thus the within-host competition between parasite genotypes that favors higher virulence (pathway 2 in Fig. 1; *4, 16*). In addition to these consumptive effects, predators can also affect host traits (*18*), non-consumptively shifting how hosts interact with their parasites (*19–21*), and probably virulence evolution. One example is host grouping, a common defense (*5*) which effectively decreases predator-induced mortality (*22*), potentially selecting for lower virulence (pathway 3 in Fig. 1). However, grouping rate also increases host-host contact rate, increasing the force of infection, so host grouping could increase multiple infections and therefore select for higher virulence (pathway 4; *6, 16, 23*). Both consumptive and non-consumptive effects of predation act simultaneously in natural communities (*20*), but their relative importance for selection on parasite virulence lacks empirical and theoretical clarification.

**Figure 1.**
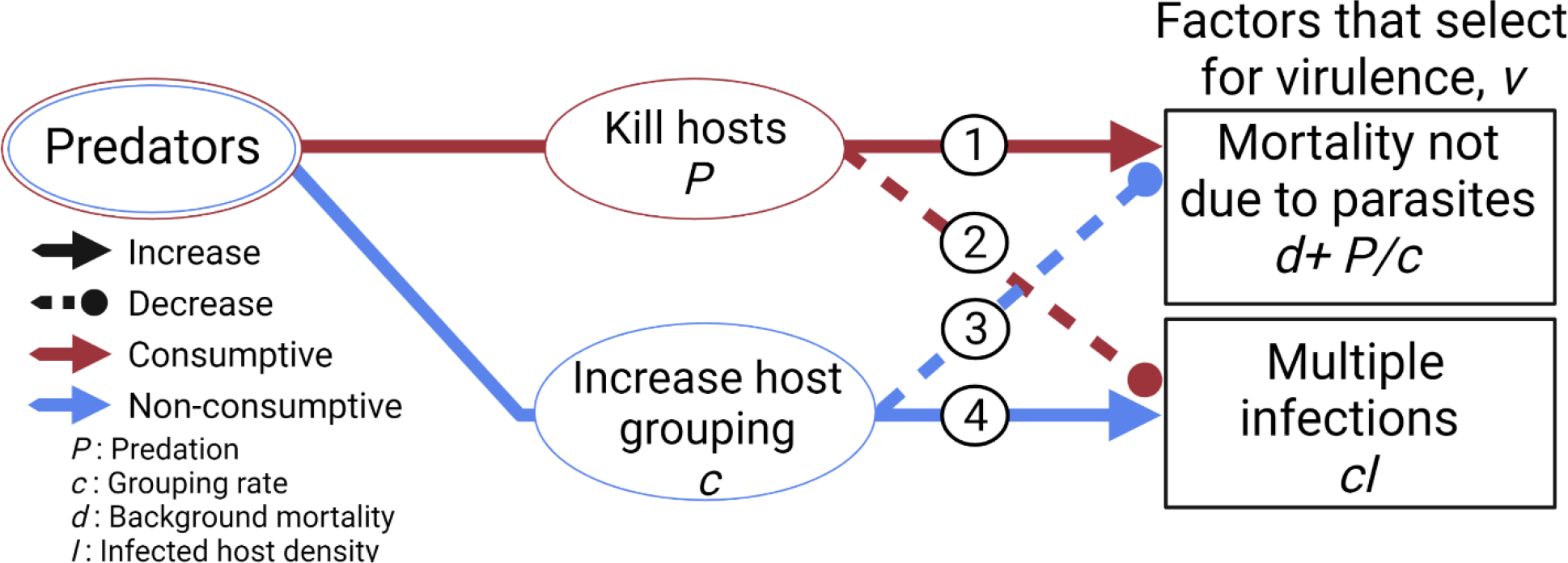
Predators alter selection on virulence through consumptive and non-consumptive pathways. The consumptive (red) and non-consumptive (blue) effects of predation simultaneously act to increase (solid, triangular arrows) and decrease (dashed, rounded arrows) factors that select for higher virulence. Mortality not due to parasites is *d+P/c* while the force of infection for multiple infections is proportional to *cI*. Host grouping rate, *c*, controls host-host contact rate for transmission.

### Scope of study

Here, we elucidate how predators affect virulence using theory and data from Trinidadian guppies, *Poecilia reticulata* and their *Gyrodactylus* spp. parasites (Fig. 2). Persistent, natural variation in predation risk drives population-level variation in guppy grouping rate (‘shoaling rate’) (*24–26*), probably influencing transmission of *Gyrodactylus* spp. ectoparasites (*27–30*) : high-predation populations shoal more and suffer higher *Gyrodactylus* spp. infection prevalence (*28, 31, 32*). First, we test for the commonly-assumed virulence-transmission trade-off and multiple infections (Table S2). We then use laboratory and field measurements to parametrize a model with selection on host and parasite phenotypes interacting with host and parasite densities (‘eco-coevolutionary model’). Finally, we test model predictions against wild-collected, *Gyrodactylus* spp. parasites, assaying parasite virulence in common garden conditions. The common garden isolated differences in parasite exploitation of hosts while keeping host defenses constant.

**Figure 2.**
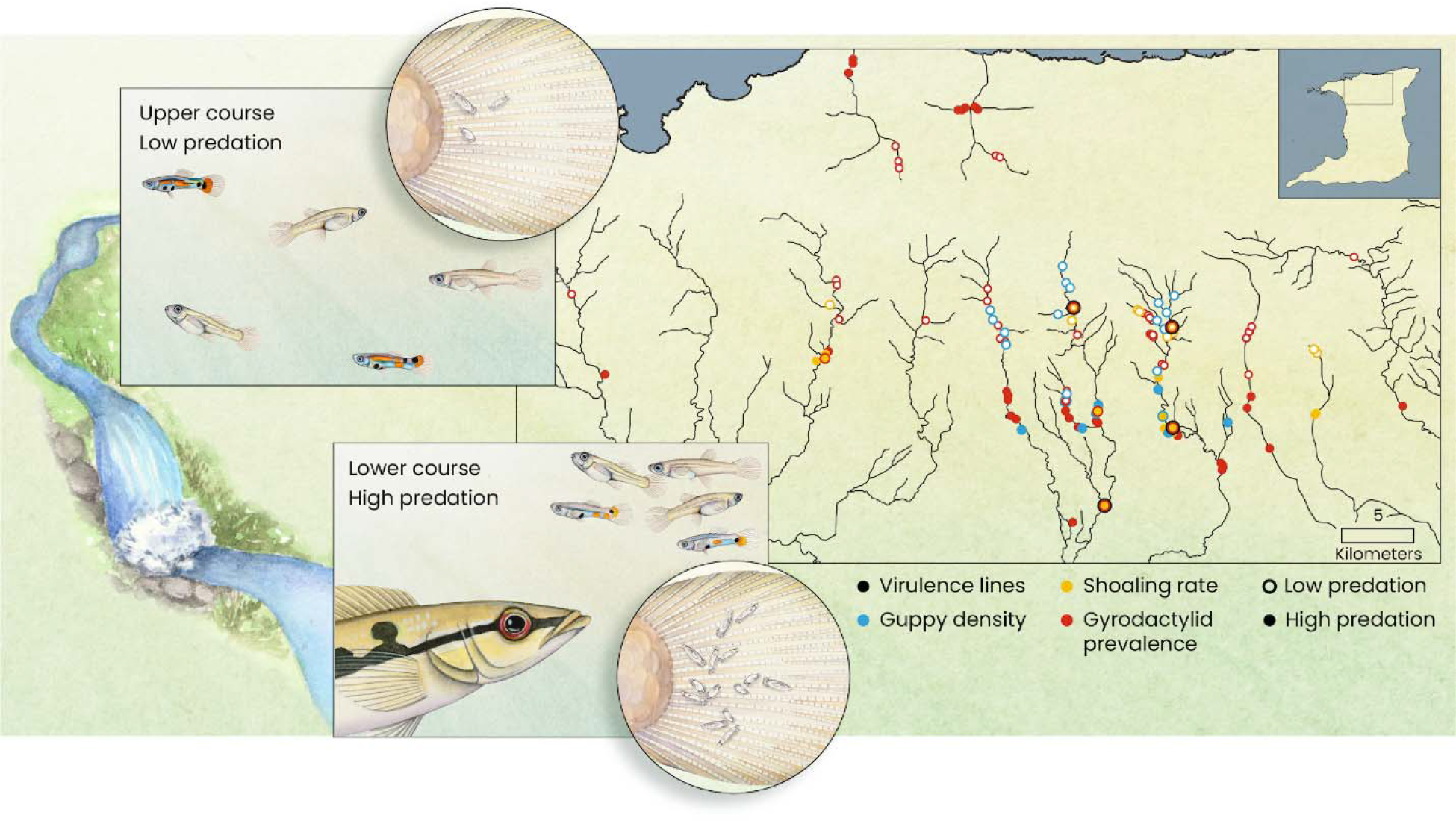
Natural guppy populations differ in predation, driving evolutionary divergence in shoaling rate. Waterfalls divide upper and lower course guppy populations, preventing upstream migration of large piscivores. Natural populations have therefore evolved under different predation regimes, replicated across rivers. Shoaling rate differences apparently drive population-level differences in transmission rate of (*28–30*), and thus selection on the virulence of, their highly prevalent, directly transmitted monogenean ectoparasites *Gyrodactylus* spp.. The map shows locations and data types that parameterized and validated model quantities [see Materials and Methods]. To focus on the various effects of predation, we used data from a river if that data type was available in low- and high-predation populations for that river.

## Results

### Gyrodactylus spp. transmission trades off with virulence, mediated by parasite intensity

We found a virulence-transmission trade-off among *Gyrodactylus* spp. lines by measuring disease traits on individual hosts. First, we found that transmission rate from a donor to sentinel host increased asymptotically as intensity increased [Fig. 3A; β = 6.08x10 intensity^0.138^; Generalized linear model (GLM) *N* = 101, *P* = 3.48 x 10^-5^, *r* = 0.35; see Materials and Methods for more details]. This pattern did not differ across two experiments using different, domestic parasite lines (isolated from commercial guppies instead of wild-caught; line effect: GLM, *N* = 101, *P* = 0.596, φ = 0.12). Second, we measured worm parasites per fish (intensity) and the death rate of infected hosts for 22 laboratory-maintained lines (including three of four from Fig. 3A). These lines differed in the mean intensity they attained (ANOVA, *N* = 1171, *P* = 9.36 x 10^-25^, η^2^ = 0.13), likely due to faster exploitation of and reproduction on individual hosts; higher intensity lines imposed a higher mean death rate on infected hosts (Fig. 3B; GLM, *N* = 22, *P* = 0.008, *r* = 0.49). Across intensities, domestic lines imposed less death (GLM, *N* = 22, *P* = 1.24 x 10^-4^, *r* = 0.63; domestic = 0.016 *vs*. wild = 0.052 at overall mean intensity). We used back-transformed, partial residual death rate to control for non-focal predictors (line origin: Fig. 3B shows the relationship as if all lines were wild) so that we could determine the relationship relevant to wild parasites (all results are very similar if instead we set all lines as domestic, see Supplementary Code).

**Figure 3.**
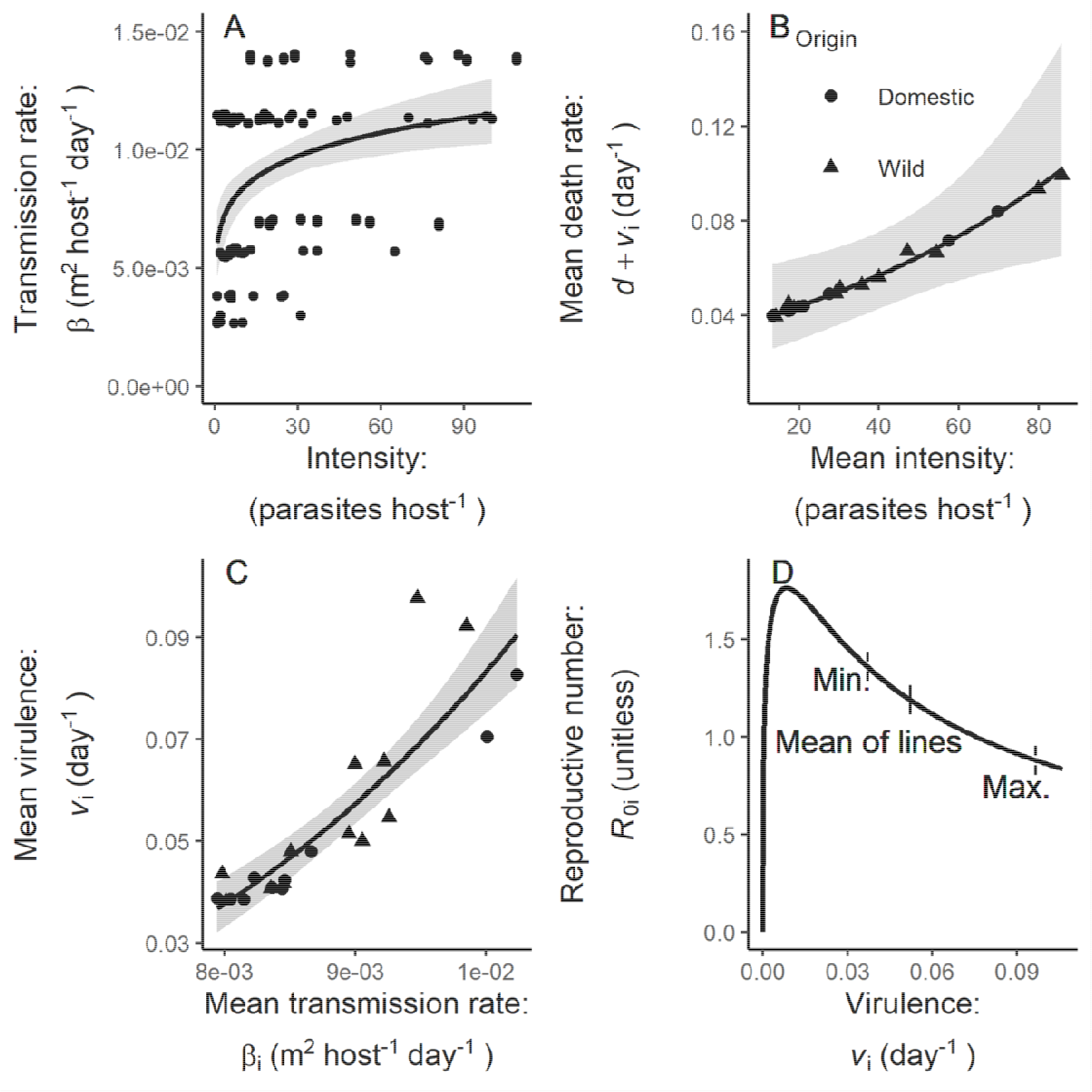
Infection intensity links transmission rate and virulence with a stabilizing trade-off. (**A**) Across four parasite lines, transmission rate increased with infection intensity. Points are transmission events jittered vertically. (**B**) Across 22 [‘domestic’ (circles) or ‘wild’ (triangles)] parasite lines, higher intensity (a measure of parasite growth rate) induced higher host death rate (d+v_i_, a measure of virulence; back-transformed partial residuals). (**C**) (A) allows a link between transmission rate and virulence across lines. Points in (B, C) are mean line traits. In (A-C), bands are 95% C.I. **(D),** The curvature in (C) maximizes R_0i_ at intermediate transmission rate/virulence.

Third, we predicted the mean transmission rate for each line, given its intensity measurements and the transmission-intensity relationship in Fig. 3A. We found that the mean transmission rate for each line traded-off with the mean virulence (we get virulence, *v*, from infected host death rate, *d+v*, given *d* = 1.30 x 10^-3^ day^-1^): parasite lines that attained higher intensity benefitted from higher transmission rate but higher virulence shortened their infections, on average (Fig. 3C). Since *Gyrodactylus* spp. are capable of sexual and asexual reproduction, we tested whether any of our lines were clones (i.e., identical multi-locus genotypes, MLGs).

Genotyping a subset (13/22, based on sample availability) found no identical MLGs (all pairs within populations differed at ≥ 16 loci, representing at least half of all variable loci for each source population, see Table S2).

The curvature of this trade-off indicated stabilizing selection on virulence (GLM fit: *v*_i_ = 1.38 x 10^6 3.61^; *v* and are average traits for a line). Bootstrapping parasite lines included in the analysis showed this curvature was significant (exponent > 1 in all 10^4^ bootstrapped samples) and AIC indicates a better fit to the data than for a linear model (ΔAIC = 13.3). Theory predicts that the trade-off curvature leads to optimal parasite fitness at intermediate transmissibility (β_i=_*cT*_i_, *c* is host shoaling rate, *T*_i_ is a line’s transmissibility) and virulence [Fig. 3D; *R*_0i_ = *cT*_i_*S*/(*d*+*v*_i_+*y*) for pure infections where we use reasonable values of *S* = 10 susceptible host m^-2^-1 and γ = 0.020 day , see Table 1]. Unsurprisingly, parasite lines fell to the right of this optimum: higher background mortality in wild than laboratory guppies shifts the optimum higher; the mixed-stock guppies in our common garden assay may differ substantially from local, wild hosts; and multiple infections (common in the wild, Table S1) select for higher virulence than would optimize *R*_0i_ of pure infections (*4*).

**Table 1.**
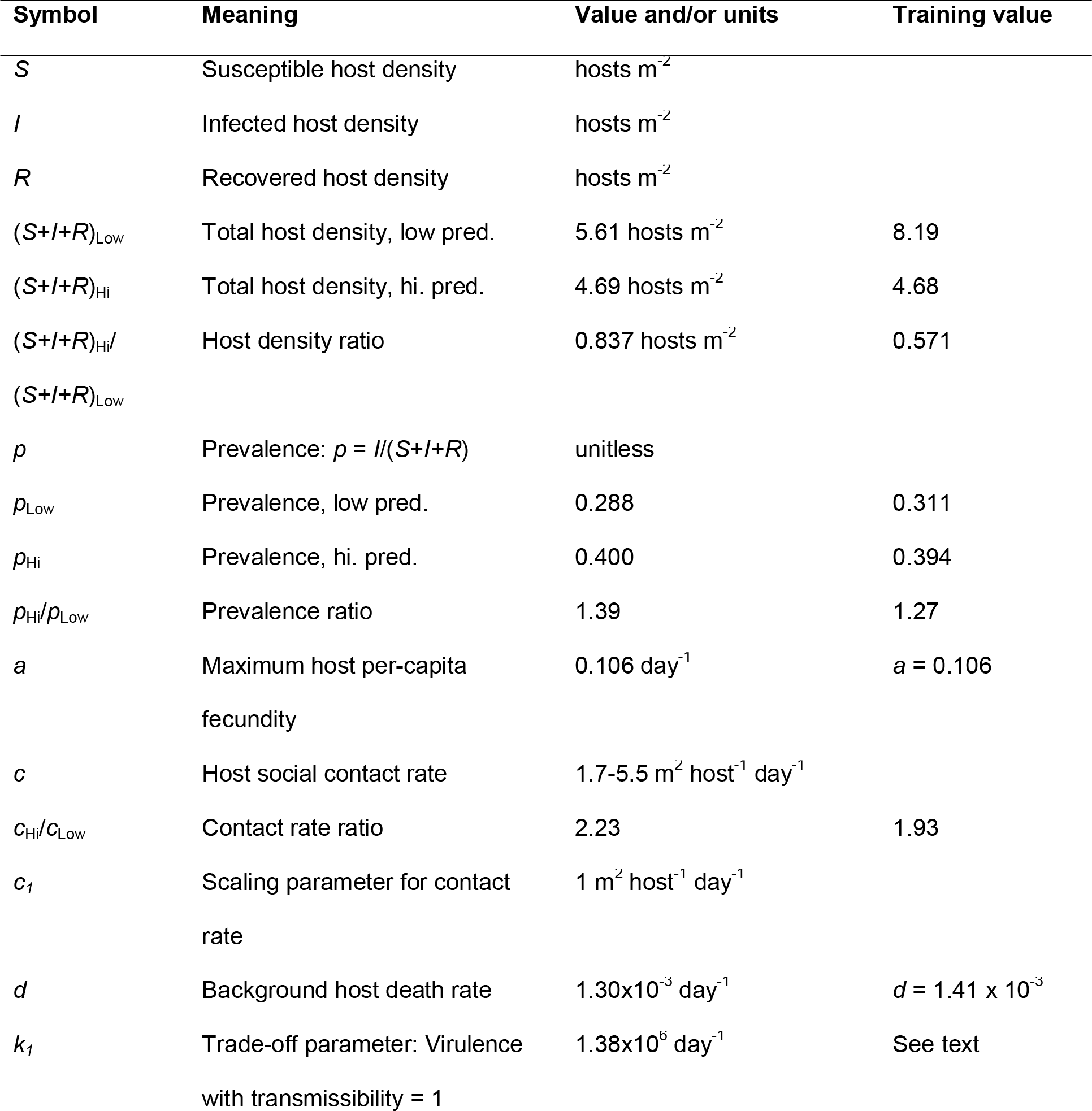
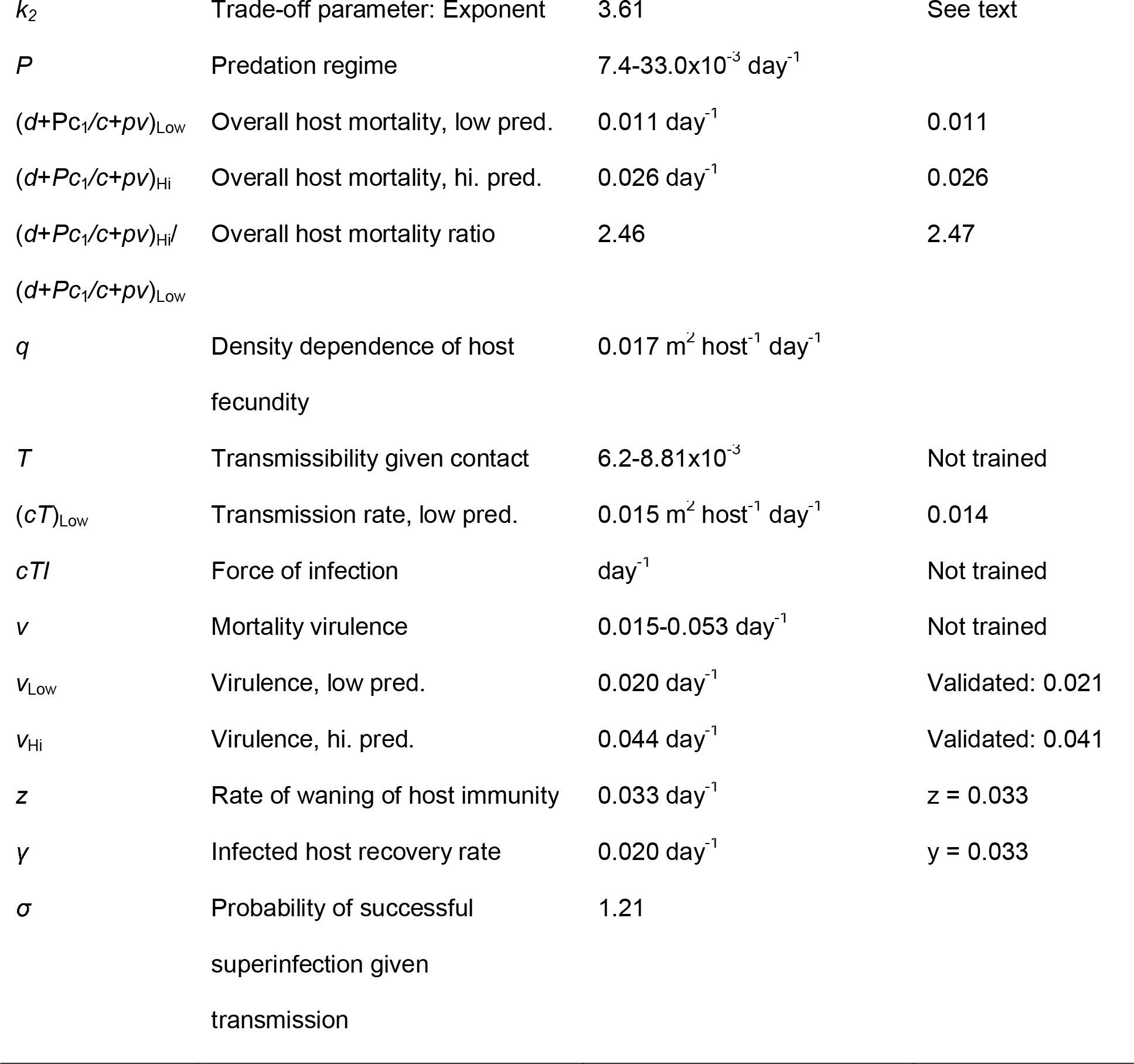
Meaning, value, and units for state variables, parameters, or outputs. Training values may be for a parameter, an output, or outputs related to a parameter or output. Training values were used in the model fitting and their sources are provided. Low/hi. pred. indicates predation regime. Values result from the model training. Note that the model was not trained with evolved virulence data but instead validated against it. See Materials and Methods for sources of training values.

### Eco-coevolutionary model resolves complexity to show that social hosts have deadlier parasites

Our model of predators that shift coevolution of host shoaling rate and parasite virulence provides two general insights. First, it demonstrates that the net, selective effect of predation, shoaling rate, or other ecological factors on virulence evolution depends only on their effect on the rate at which infections are lost to mortality or multiple infections (inverse of infectious period; see Supplementary Text for proof). Second, the model shows that shoaling rate evolves to balance predator-induced mortality against parasite-induced mortality. When predation increases, hosts evolve higher shoaling rate, increasing transmission rate. When parasites are more abundant and virulent, hosts evolve lower shoaling rate to prevent infection. For some parameter values, higher virulence can select for higher shoaling rate if parasites become so virulent that the density of infected hosts and force of infection decline strongly (Fig. S1).

When parametrized with empirical data from the guppy-*Gyrodactylus* spp. system, the model complexity resolves into one dominant pathway: predation increases co-evolutionarily stable contact rate, multiple infections, and thus virulence (Fig. 4; strength of pathways from Fig. 1: pathway 1 = 0.131, pathway 2 = -0.166, pathway 3 = -0.063, pathway 4 = 1.18; see Supplementary Text for derivation). When predators only have consumptive effects on predation (i.e., hosts do not evolve; pathways 1 and 2), predation decreases prevalence and virulence (grey curves in Fig. 5A-E; presence/absence of parasite evolution has little effect on grey curves).

**Figure 4.**
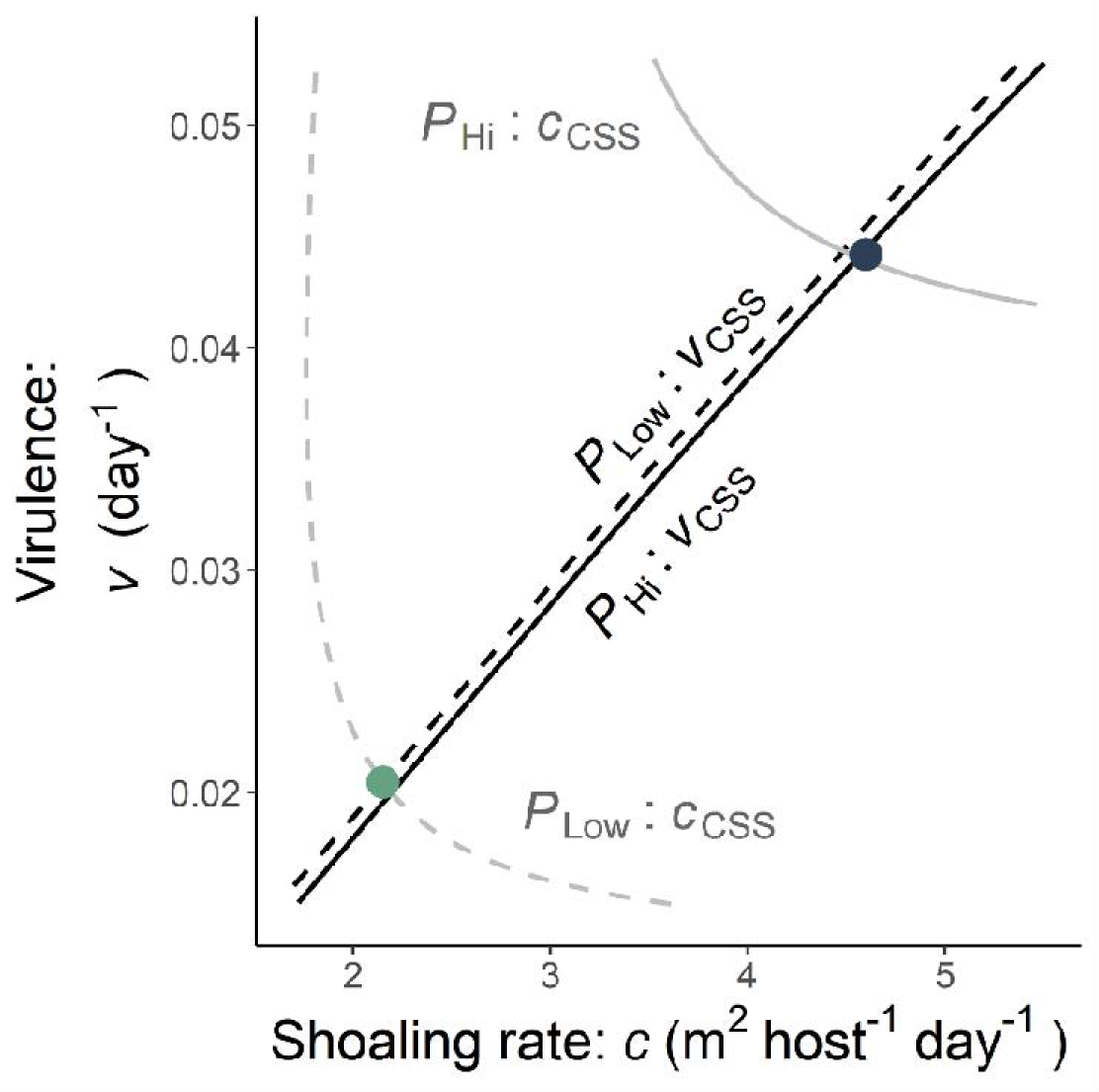
Predation drives increased shoaling rate and virulence in the eco-coevolutionary model. Curves give the continuously stable strategy (CSS) for parasite (*v*_CSS_ black) or host (*c*_CSS_ grey) evolution at predation levels fitted to correspond to natural populations (*P*_Low_; dashed or *P*_Hi_; solid). Parasites evolve higher virulence with higher shoaling rate. The consumptive effect of predation have little, net effect on evolved virulence. Hosts evolve lower shoaling rate with increasing virulence (grey curves move toward lower *c* as *v* increases). Predation substantially increases *c*_CSS_. Host and parasite curves intersect at coevolutionarily stable points (green and blue points). As predation increases (from green point to blue point), coevolutionarily stable shoaling rate (*c*_coCSS_) and virulence (*v*_coCSS_) increase.

**Figure 5.**
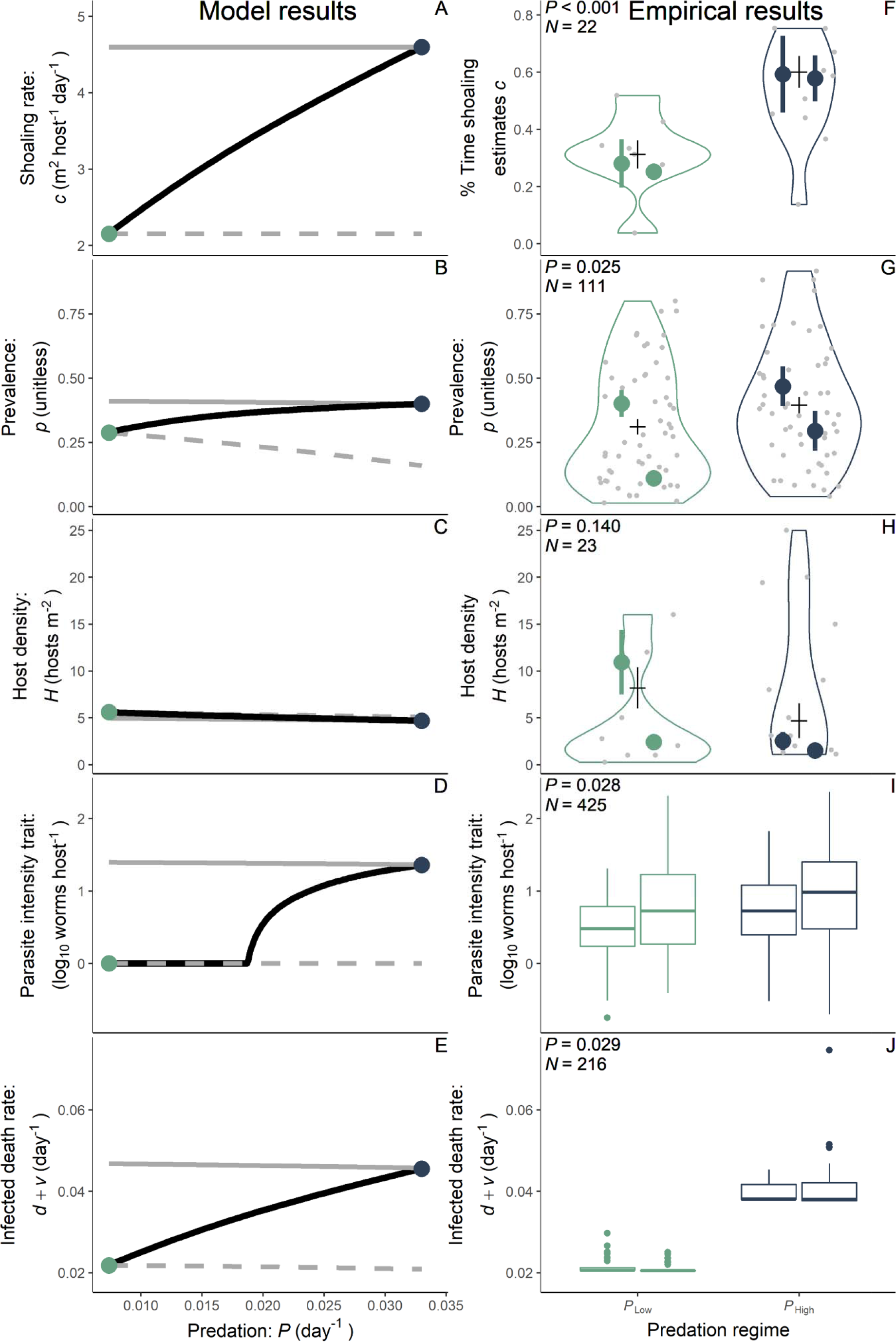
Predation increases shoaling rate and thus selects for higher virulence. (**A-E**) Model results. Black curves: eco-coevolution (connecting Fig. 4 colored points). Grey curves: no host evolution. (**F-H**) Empirical training data. Horizontal segments: predation regime means. Grey points: one river/regime/year mean. Colored points: one focal river/predation regime mean (Aripo left of Guanapo). Vertical bars: standard errors. Training data: violins. Validating data (**I, J**): boxplots (back-transformed partial residuals; center line, median; box limits, first and third quartiles; whiskers, 1.5x interquartile range; points, outliers). *P*-values for the effect of course and sample sizes (*N*) provided. (A, F) Shoaling increased with predation. (B, G) Prevalence increased with predation while (C, H) host density decreased non-significantly. (D, I) Parasite intensity [note, we do not model intensity lower than log(1 worm)=0] and (E, J) virulence increased with predation.

However, as hosts evolve higher shoaling rate in response to predation (Fig. 5A), predation also increases prevalence (black curve in Fig. 5B) in all tested parameter sets (see sensitivity analysis methods). Increasing predation increases overall host mortality and reduces host density, particularly when hosts evolve (Fig. 5C). Further, most of the increase in mortality (76%) with predation is due to increased parasite-induced mortality (prevalence × virulence), rather than increased predator-induced mortality (Fig. S2; robust to parameters: Fig. S3). Higher shoaling rate increases the force of infection (and thus multiple infections), selecting for higher parasite intensity and virulence (Figs. 5D, E; note intensity was not modelled directly but back-calculated from transmissibility according to Fig. 3B); in all parameter sets, higher predation increased virulence. Importantly, predators increase virulence through their non-consumptive effects on shoaling rate (across parameter sets, see Fig. S3).

### Empirical results quantitatively match model predictions that social hosts have deadlier parasites

The model’s coevolutionary outcomes successfully fit training data from our collections and the literature (Fig. 5A-C, F-H; table 1). High predation fish spent more time shoaling in our assay (Generalized Linear Mixed Model [GLMM], *N* = 68, *P* = 0.006, *r* = 0.32) and across population estimates extracted from the literature (GLMM, *N* = 22, *P* = 8.31 x 10^-4^, *r* = 0.58; Fig. 5F). This helps explain why high predation populations suffered higher parasite prevalence across multiple rivers, sites, and years (GLMM, *N* = 107, *P* = 0.025, *r* = 0.21; Fig. 5G). Higher levels of predation and parasitism may depress host density (though not significantly so: GLMM, *N* = 23, *P* = 0.140, *r* = -0.29; Fig. 5H). Overall, the model fit the training data reasonably well, with an average relative error of 10.5% per fitted quantity (shown in Fig. 5A-C, F-H and Table 1).

Alternative models do not fit the data as well, bolstering confidence in our focal eco- coevolutionary model. A model without host evolution (only consumptive effects of predators) fits the data poorly (relative error of 15.2% per fitted quantity, see Fig. S4) despite having an additional free parameter. A different alternative model fits an exponent governing how effectively shoaling protects from predation (exponent = 1 in our focal model) but the miniscule shift in model fit (relative error improves from 10.4% to 10.1%, see Fig. S4) does not justify the additional free parameter. Accounting for model complexity, our focal model provided the best fit to the training data, and of the three models it thus likely best captures the consumptive and non-consumptive effects of predation.

We tested the focal model’s trained predictions by quantifying the traits of 18 parasite lines isolated from wild populations and maintained in the lab under common garden conditions for 65 days. Lines from high predation populations attained higher intensity on infected, mixed-stock hosts (GLMM, *N* = 425, *P* = 0.028, *r* = 0.11; Fig. 5I) and induced higher death rate (GLMM, *N* = 216, *P* = 0.029, *r* = 0.15; Fig. 5J). The two parasite species, *G. turnbulli* (11 known lines) and *G. bullatarudis* (3 known lines) did not differ significantly in intensity (GLMM, *N* = 345, *P* = 0.098, *r* = 0.09) or virulence [GLM, *N* = 177, *P* = 0.345, *r* = 0.07; unlike (*33*)].

Restricting the analysis for Fig. 5J to *G. turnbulli* found marginally higher intensity (GLM, *N* = 208, *P* = 0.076, *r* = 0.12) and significantly higher virulence in high predation populations (GLM, *N* = 102, *P* = 0.009, *r* = 0.25). The overall quantitative match between the model-predicted mortality and our empirical results supports our model’s inferences (*d+v* in the model *vs.* mean of *d+v* back-transformed partial residuals: low predation 0.020 *vs*. 0.021; high predation: 0.044 *vs*. 0.041). We compared theoretical predictions to partial residuals to control for non-focal predictors in the statistical model, especially duration of parasite maintenance in the laboratory, to obtain the most biologically relevant estimate of virulence. Our theoretical model makes this prediction based on higher shoaling rate increasing multiple infections, which seem more frequent in high predation populations on a similar scale to shoaling rate (shoaling rate ∼2 times higher and coinfection ∼3 times higher in high predation; Table S1).

The model-data agreement on evolved virulence aids model-data agreement on how mortality changes across predation regime. In our model, 76% of the increased mortality across predation regime is increased parasite-induced mortality while increased predator-induced mortality accounts for 24%. Prevalence in natural populations and virulence data on laboratory fish (*p*_Hi_*v*_Hi_*-p*_Low_*v*_Low_) indicated that parasitism would explain 64% of the mortality difference from Reznick *et al* (*34*) while predation may account for the remaining 36%.

## Discussion

Our theoretical-empirical approach clarifies that predation drives the evolution of parasite virulence by increasing host shoaling rate. As a result, shoaling in response to increasing predation pressure leads to parasite-induced mortality rising more than that induced by predators. In contrast to these strong non-consumptive effects, we found that the consumptive effects of predation are small, balance one another out, and barely alter virulence evolution. To our knowledge, our study is the first to model the non-consumptive effects of predation on virulence evolution. Further, we used data to train and test a model that infers and compares the strength of consumptive and non-consumptive effects in natural populations. We discuss key emergent patterns.

Theory shows that shoaling rate drives virulence evolution depends on multiple infections. Without multiple infections, shoaling decreases predator-induced mortality and thus selects for decreased virulence (*35*). With multiple infections, we and others (*6*) find that increased grouping rate selects for higher virulence. Empirically, we do not know of previous tests of the effect of host-host contact rate on virulence evolution, but our result is analogous to more host dispersal (*15, 36, 37*) and parasite founder diversity (*14*) selecting for higher host exploitation through high local parasite diversity (*38*).

We assayed parasite traits using infections on mixed-stock, wildtype guppies: this approach allowed us to draw robust conclusions about parasite evolution but leaves untested how host defenses, other than shoaling rate, may affect disease dynamics in natural communities.

Thus far, guppy defenses against parasites have not been robustly characterized across predation regime. Illustrative data suggest that, in our focal Guanapo river (experimental test; *39*), and perhaps more generally (field surveys; *28, 31*), guppies from low predation regimes are better defended against *Gyrodactylus* spp.. Low predation guppies may be better able to evolve in response to *Gyrodactylus* spp.: they need not balance this investment with defense against predators (*22*), and sexual selection, which acts on gyrodactylid resistance (*40*), is more effective in the absence of predators (*41*). However, in our focal Aripo river, low predation guppies appear less resistant than high predation guppies (experimental test; *42*). Despite this apparent difference between our focal rivers, our measured virulence matched our model predictions for both, suggesting the patterns we observe may be robust to population-level differences in host defenses. Nevertheless, how host behavioral and immunological defenses may coevolve with parasite virulence in natural communities remains an outstanding and complex theoretical as well as empirical challenge.

The importance and implications of predation and social parasite transmission, evident from our model and data, may hold across directly transmitted parasites of group-living hosts (*43*). Predators drive defensive group-living in animals across taxa (*5*), which can increase parasitism, creating ecological and evolutionary feedbacks between host sociality and parasites (*43*). Parasites evolve along virulence-transmission trade-offs in systems ranging from viruses of humans (*10, 13*), bacterial pathogens of birds (*11*), protozoan pathogens of insects (*12*), and our monogenean fish ectoparasite. Multiple infections are common for many parasites, allowing the simple mechanism of competition for within-host resources to select for higher virulence (*16*). Thus, diverse systems likely meet the essential assumptions of our model: predators may frequently shift the antagonistic interplay of host sociality and parasite virulence, driving hosts into the arms of more virulent parasites. Conversely, these results also indicate that social distancing may select for lower virulence when parasites exhibit multiple infections and a virulence-transmission trade-off (e.g., influenza A virus; *10, 44*). Host behavior that reduces contact may thus effectively control both the spread and virulence evolution of pathogens and parasites.

## Acknowledgments

We thank Mahase Ramlal, David Reznick and Elizabeth Rudzki for assistance with fieldwork. Emmalina Calcaterra, Lindsay Colgan, Jukka Jokela, Maura Sackett, and Nadine Tardent provided technical assistance with parasite genotyping. Julie Johnson made artwork for Fig. 1. Jukka Jokela, Cock van Oosterhout, Martin Turcotte, and Kyle A. Young commented on an earlier version of this manuscript.

## Author contributions

Conceptualization: JFS, JCW Theoretical modeling: JCW, CEC, JFS Data collection from literature: FR Sensitivity analysis: JCW, FR

Field collections: MJJ, DRC, RP, RSM

Laboratory trait measurements: RP, DRC, MJJ, RDK, JFS Parasite molecular work: MJJ, RDK, MK

Parasite genetic analysis: MJJ, MK Trait data analysis: JCW, DRC, JFS

Density and prevalence data analysis: JCW Funding acquisition: JCW, MJJ, FR, JFS Writing – original draft: JCW, JFS

Writing – review & editing: JCW, MJJ, DRC, RDK, FR, RP, RSM, MK, CEC, JFS

## Data and materials availability

All data and code are available in the main text or the supplementary materials.

## Supplementary Materials

Materials and Methods Supplementary Text Figs. S1 to S4

Tables S1 to S3 References (*45–72*) Supplementary Code Data files S1-S7

## Materials and Methods

### 1. Molecular methods

We used two methods of molecular species identification for individual parasites: we sequenced the mitochondrial COII gene for a subset of domestic lines established before March 2020, and we used a newly developed restriction enzyme assay for wild and domestic lines established after March 2020 (see Supplementary Text for further details). To prevent disrupting data collection, sample collection was restricted to already-dead hosts, stored in 70 % EtOH. This conservative precaution meant we were unable to collect useable samples from some of our parasite lines.

We developed panels of single nucleotide polymorphisms (SNPs) for *G. turnbulli* and *G. bullatarudis* to examine the incidence of multi-genotype coinfections from wild collections, and to verify that the lines we established in the lab were genetically distinct. Genotypes were called using Fluidigm SNP genotyping analysis software. A subset of individuals was re-genotyped to identify and remove error-prone loci and estimate error rates (the proportion of mismatches to matches for re-genotyped loci). Loci which amplified consistently and with score calls greater than 90% were selected for use in the analysis. This resulted in a total of 140 variable loci for *G. turnbulli* (error rate 1.5%) and 83 loci for *G. bullatarudis* (error rate 2 %) across all populations. However, due to significant local variation in informative loci, the total number of loci used for each source population varied (see Table S2). The resulting multilocus genotypes (MLGs) were assessed using GENALEX 6.502 to estimate the fraction of multi-genotype infections at six sites across Trinidad including our four focal populations (low- and high-predation in the Aripo and Guanapo rivers), and to assess the genotypes of the established lines.

As a conservative estimate, two individuals were considered to have different genotypes if they varied in at least 50% of the total variable loci from a given source population (see Supplementary Text for further method details). This applied both to determining that our lines were different genotypes and finding the frequency of multiple infections. In finding the frequency of multiple infections, we genotyped worms from fish with at least two worms and accounted for the proportion of fish with only one worm, estimated for each site from our 2020 survey and Stephenson *et al*. (*28*).

### 2. Parasite trait data

We established parasite lines by transferring a single worm to an uninfected host from our mixed, laboratory-bred stocks descended from wild populations. Each founding worm was obtained from a single guppy either from a commercial supplier (“domestic” lines) or wild-caught in Trinidad (“wild” lines from Caura, Aripo, Guanapo, and Lopinot rivers in Caroni drainage). Wild-caught adult guppies (∼50/population) were shipped from Trinidad to the University of Pittsburgh in March 2020 (see Animal use ethics statement). We established 43 parasite lines (18 domestic and 25 wild, 10 from low predation populations and 15 from high predation). Different lines were included in different analyses, as explained below. Lines were maintained under common garden conditions for 65 days (some domestic lines were maintained in the lab much longer) on groups of 3-6 guppies. We added uninfected fish to each line as required to replace those that either died or were found parasite-free during twice-weekly screening of all fish in each line under anaesthetic (tricaine methanesulfonate; 4g/L) using a dissecting microscope. During these screens, we recorded the number of parasites infecting each fish. Each line was housed in a single 1.8 L tank on a recirculating system (Aquaneering Inc.; 12L:12D; 24°C). Recirculated water passes through fine foam, sand, and ultraviolet filters before re-entering other tanks: individual parasites cannot transmit between lines, as supported by our genotyping.

We estimated transmission rate by re-analysing data from Stephenson *et al.* (*45*) and conducting a similar new experiment. Donor fish received an infection of one parasite line or multiple infection and were individually housed in 1.8 L tanks [donor fish received line A, B, C, or A and B]; no lines in the new experiment were used by Stephenson *et al.* (*45*). We added a parasite-naïve recipient fish on day 8 of the donor’s infection, and screened both fish for infection every 24 hrs.

We also assayed intensity and death rate. For intensity (a proxy for within-host growth rate), we counted the number of parasites on each host at each date (1 observation). We estimated the per-capita host death rate from the number of fish found dead divided by the number of infected fish in the tank at the previous observation time point and the days between observations (1 observation; 79% of observations were 3±1 days after the previous one).

We calculated transmission rate from the number of days until successful infection of the recipient and intensity as the number of worms on the donor fish on the day of transmission, following Stephenson *et al.* (*45*) (total *n* = 101 transmission events). Worm numbers changed slowly on the scale of the number of days until successful infection. In these assays, dS/dt = - *cT_i_IS* (*c* is contact rate, *T*_i_ is the transmissibility of parasite line i, *I* is infected host density, and *S* is susceptible host density) so susceptible hosts follow an exponential decay pattern; thus, mean time until infection is 1/(*cT_i_I*). If days until transmission are *D*, then the transmission rate estimated by a given transmission event is *cT_i_* = *1/(DI)*. Based on the area of enclosures, infected host densities were *I* = 88.5 hosts m^-2^ for Stephenson *et al.* (*45*) and *I* = 72.0 hosts m^-2^ for the follow-up experiment. We fit these estimates of transmission rate to log_10_ intensity with parasite line (line A, B, C, or coinfection of lines A and B) as a fixed effect using a generalized linear model with a Gamma error family and log link function. Parasite line was non-significant, so we refit the model without parasite line (P-value for intensity reported from this model; all P-values represent two-sided, Type II Wald chi-square or F tests for this and other methods). Effect sizes are ^2^ η for ANOVA with only two levels of predictors, φ for ANOVA with more than two levels of predictors, or partial correlation coefficients, *r*, for other analyses. For all analyses, we provide the code, output, and validation of the model fit in the Supplementary Code.

When determining average traits for a line, we only included lines maintained in the laboratory for more than 30 days (*n* = 22; 10 domestic, 12 wild lines, including 1-4 lines from each of the four focal populations). We used an ANOVA to determine whether lines differed significantly in log intensity (1171 observations of intensity for the 22 parasite lines, at least 25 for each line). We modelled mean death rate for each line (344 observations and at least 13 for each line) as a function of mean intensity and line origin (wild *vs.* domestic) with a beta error family and logit link function. We estimated virulence (*v*) by subtracting background death rate (see Table 1) from partial residuals of infected death rate (to get *v* from *d*+*v*). We used partial residuals of infected death rate back-transformed onto the response scale to control for non-focal predictors (here, line origin) and find the relationship for wild parasites. We estimated mean transmission rate for a line by mapping the individual intensity measurements for each line onto the relationship observed between intensity and transmission in the experiments described above (Fig. 3A); we then calculated the mean transmission rate for each line. With estimates of transmission rate and virulence for each line, we examined the relationship between them using a generalized linear model of virulence with Gaussian error distribution and log link function as a function of natural log transmission rate. The statistical model was free to fit a positive (exponent > 1), neutral (exponent = 1), or negative curvature (exponent < 1) to these data. We bootstrapped this fit by sampling lines to include in the analysis from the 22 parasite lines with replacement to determine how often the exponent was more than one.

To examine the patterns in Fig. 5I and 5J, we used our estimates of parasite intensity and host death, (calculated as above) as response variables in GLMMs. These traits were assayed from parasites collected from low and high predation populations of our focal rivers (low/high predation; river distances: low Guanapo-high Guanapo = 11.1 km, high Guanapo-high Aripo =

23.9 km, high Aripo-low Aripo = 6.6 km; 3 lines from high-predation Aripo and 5 from each of the other focal populations; 18 lines total). Each observation was of one line at one date but intensity on and death of multiple hosts for each line. This analysis included some lines maintained in the laboratory for less than 30 days but these lines necessarily had fewer observations and thus did not influence the analysis as much as lines with more observations. For intensity (*N* = 425 observations), we used predation regime, river, and number of fish in the common garden at time of measurement (since *Gyrodactylus* spp. can move relatively freely among fish) as fixed effects. Parasite line and lineday (days the line had been in the lab) nested within parasite line were included as random effects. We used log(intensity) as the response variable. For death rate (*N* = 216), we used predation regime and river as the fixed effects with parasite line and lineday nested within line as random effects, a Tweedie error family, and a log link function. The same analyses were used when determining the effect of predation regime on intensity (*N* = 208) and host death rate (*N* = 102) for known *G. turnbulli* lines. We compare the theoretical model to mean of back-transformed partial residuals of infected host death rate to control for non-focal predictors, such as lineday, to find the most representative estimate of death rate in each predation regime.

To determine trait differences by species, we considered all wild lines that were identified to species (including 7 from rivers considered non-focal due to lack of high predation *vs.* low predation comparisons). Three of these lines were *G. bullatarudis*and 11 were *G. turnbulli*.

Because all three *G. bullatarudis* lines were from high predation populations, we only compared species differences within that regime. Species identity was used as a predictor instead of predation regime but statistical models were otherwise identical to those for the effect of predation regime for intensity (*N* = 345) and infected host death rate (*N* = 177).

### 3. Theoretical modelling

We wrote a relatively simple, ordinary differential equation model that retains the key biology of the predator-prey/host-parasite system. In the model, predators are not limited by prey/host density (e.g., because predators are generalists), allowing us to represent predation as a parameter (*P*) for simplicity instead of a state variable. Susceptible hosts (*S*) become infected (*I*) and can recover to an immune state (*R*) with waning immunity (see Table 1 for symbols, values, and units):

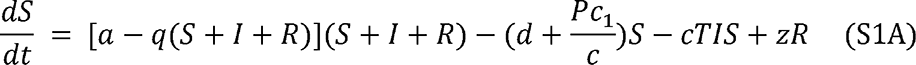

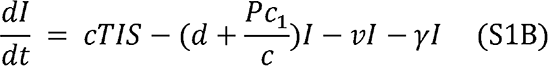

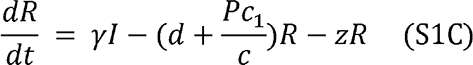

Susceptible hosts are born from all hosts with density dependent, per-capita birth rate *a*- *q*(*S+I+R*) (i.e., hosts grow logistically, eq. S1A). Hosts die at background rate *d* and from predation at rate *Pc*_1_/*c* that decreases with host shoaling rate (*c*; that *P* and *c*_1_ together determine the strength of predation). Along with parasite transmissibility given contact (*T*), the rate of density-dependent transmission from *I* to *S* depends on the host-host contact rate, dictated by shoaling rate. Recovered hosts also move back into the susceptible class as immunity wanes at rate *z* (average duration of immunity is 1/*z*).

Infected hosts (eq. S1B) suffer background mortality and predation (predation is not selective in this model) while suffering additional mortality due to virulence, *v*. If predation was selective, especially to remove the sickest hosts, the consumptive effects of predation would be even more likely to select against virulence (*3*), strengthening our conclusion that shoaling rate drives virulence. Infected hosts recover with rate γ (eq. S1C). Recovered hosts suffer background mortality and predation while losing immunity over time.

We modelled parasite and host evolution via Adaptive Dynamics (*46*). Biologically, parasite evolution is rendered more complex by the presence of multiple lines as well as two phenotypically similar species; host evolution is also complex because shoaling rate has important genetic and plastic components (*27*). For both host and parasite, our model is agnostic regarding the basis of adaptation but simply examines competition between phenotypes based on fitness of a rare phenotype (i.e., invasion analysis). Each phenotype corresponds to an asexual genotype in the model. Fitness when invading depends on the traits of the mutant (m) and resident genotypes (r) for parasites (eq. S2A):

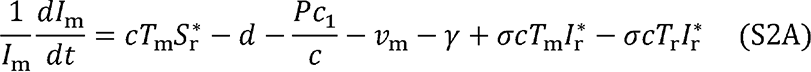

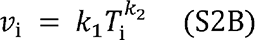

The resident sets host densities at *S*_r_ and *I*_r_ which depend on its traits, *T*_r_ and *v*_r_ (linked by the trade-off in eq. S2B). Mutant fitness depends on transmission to susceptible hosts, background death, predation, virulence, recovery, and gains and losses due to superinfection (eq. S2A). In superinfection, parasites of one genotype take over a host already infected by another genotype. We assume that each genotype has the same probability of successful superinfection given transmission, σ. This assumption is likely conservative as the probability of successful σ superinfection is often expected to increase with virulence (*4*), which could amplify the impact of multiple infections to select for higher virulence. From parasite fitness when invading, we find the continuously stable strategy, ‘CSS’, (*46*) for parasite virulence; a CSS represents a trait value that can be reached by gradual trait changes but that cannot be invaded by rare phenotypes with slightly different traits.

For host fitness, we used the Next Generation Matrix technique (*47*) that accounts for fitness within each class and the rates of movement between classes (see Supplementary Code). In the model, host genotypes differ in their contact rates, *c_i_*. An invading host genotype (contact rate *c*_m_) contacts the resident genotype (*c*_r_) at rate sqrt(*c*_m_*c*_r_) (for the purposes of transmission and predation, eq. S3), following Bonds et al. (*35*). An invading genotype suffers crowding (-*q* term, eq. S3A) from all hosts. Only intersections of a host CSS curve and parasite CSS curve (Fig. 4) can be a coevolutionary stable point (coCSS). We confirmed that a potential coCSS is indeed a coCSS with the strong convergence stability criterion (*48*).

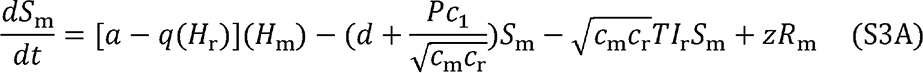

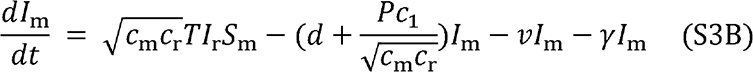

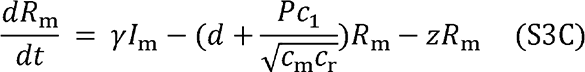

Note: *H*=*S+I+R*. In the second alternate model, the rate of predation on the resident is *Pc*_1_/*c*_r_*^x^* and predation on the mutant is *Pc*_1_/(*c*_r_*c*_m_)*^x^*^/2^ where 0 < *x*. *x >* 1 indicates that protection from predation more than doubles when shoaling rate doubles. *x* < 1 would mean that protection from predation less than doubles when shoaling rate doubles.

We have empirical estimates of some, but not all, of the parameters of the eco-coevolutionary model. As such, we used a simple evolutionary algorithm (described below) to estimate all model parameters, using empirical parameter estimates and field estimates of model outputs (e.g., prevalence) to fit the model to the data. Importantly, we did not use empirical data on evolved virulence in the fitting process, i.e., model virulence is free to evolve along the virulence-transmissibility trade-off however best fits the other data; instead, we used virulence data to validate the model.

Many empirical estimates came from the literature (see below). We estimated maximum birth rate (*a*) from lifetime fecundity of guppies with high food (*49*) in the laboratory. We estimated background mortality in the absence of parasites and predators (*d*) as the inverse of life expectancy in the laboratory (*49*). We estimated recovery rate (γ) as the inverse of the time γ required for fish to clear infection in the laboratory (*50*). We estimated the rate at which immunity wanes (*z*) from the observation that fish were roughly half resistant (averaging across totally resistant, partially resistant, and not resistant) 21 days after initial infection (*51*). Our trait measurements provided estimates for the trade-off parameters (Fig. 3C) given an estimate of *d*.

For parameters where we lacked estimates [importantly *Pc*_1_ (which acts as one parameter) in low- and high-predation populations and the superinfection parameter, σ , we also fit the model’s eco-coevolutionary outputs at low and high predation to estimates of such outcomes in the field. We converted recapture rates from a mark-recapture experiment (*52*) into instantaneous mortality rates to determine overall mortality rate in the field (*d*+*Pc*_1_*/c+pv*) for both a high predation population and a low predation population. We found a transmission rate reasonable for low predation populations (*cT*_Low_) by fitting transmission rate to peak prevalence and timing of peak prevalence found for low predation Aripo guppies and parasites in a stream mesocosm (*53*). We also used estimates of shoaling rate, prevalence, and host density (see Analysis of shoaling rate, prevalence, and host density below). We were able to fit some outputs to estimates from low predation populations, high predation populations, and the ratio of the two (representing the impact of changing predation regime). For others, we only have estimates of one of these three. This scheme weights the model training toward quantities that are well-characterized. See Table 1 for estimates used.

We optimized the model’s fit to both these empirical parameter estimates and field-estimated model outputs with a simple evolutionary algorithm (*54*). We began with parameters at their estimated values (for *a*, *d*, *k*_1_*, k*_2_*, ,* and *z*). Initial values of other parameters were chosen arbitrarily [*q* = 1 x 10^-2^, (*Pc_1_*)_Low_ = 1 x 10^-2^, (*Pc_1_*)_Hi_ = 3 x 10^-2^, σ = 0.5]. Each parameter was mutated by a factor of 10~*x* where *x* is a single sample from the normal distribution *N*(0,0.05) to create a parameter set; each parameter set formed one strategy in the evolutionary algorithm.

One un-mutated strategy and 299 mutated strategies were evaluated in one “generation” of the algorithm by their summed relative error from all inputs and outputs with known estimates: summed relative error of strategy x = |*a*_x_-*a*_data_|/*a*_data_ +…+|*cT*_Low_,_x_-*cT*_Low_,_data_|/*cT*_Low_,_data_. There are four constrained parameters, four unconstrained parameters, and 8 independent scored outputs (e.g., *p*_Low_ is independent from *p*_Hi_ but not *p*_Hi_/*p*_Low_). The strategy with the lowest summed relative error was passed to the next generation of the algorithm along with 299 mutated versions of itself. This process continues for 20 generations of the algorithm, as the model had asymptotically converged to a fit, yielding the parameter values used for the model (see Table 1).

We performed a sensitivity analysis to determine how key outcomes responded to parameter values. We performed this analysis for coCSS parasite prevalence, host density, parasite virulence, host contact rate, overall host mortality, transmission rates, and the ratios of these quantities between high and low predation populations. We also examine two other key ‘proportion change metrics’ representing how key factors change between the two, focal predation levels. First was the proportion of total host mortality change with predation regime that is caused by increased parasite-induced mortality: (*p*_Hi_*v*_Hi-_*p*_Low_*v*_Low_)/[(*d+p*_Hi_*v*_Hi_ *+P*_High_*c*_1_*/c*_Hi_) - (*d+p*_Low_*v*_Low_ *+P*_Low_ *c*_1_*/c* _Low_)]. Second was the proportion of change in coevolutionary virulence across predation regime that is via predator’s non-consumptive effects (evolution of higher shoaling rate). We defined this as the strength of pathway 3 (negative) plus the strength of pathway 4 (positive) divided by the summed strength of all pathways (see Fig. 1); we averaged pathway strength at the low predation point and the high predation point together to get one strength for each pathway (see eq. S5 for details). To generate random parameter sets, we used Latin Hypercube sampling with the ‘lhs’ R package (*55*) and then computed the partial rank correlation coefficients of the parameters with respect to each model outcome of interest (coevolutionary outcomes and key proportion change metrics) for 5 x 10^3^ runs using the R package ‘sensitivity’ (*56*). We provide the code for the model and sensitivity analysis in the Supplementary Code.

### 4. Analysis of shoaling rate, prevalence, and host density

We extracted estimates of shoaling rate, prevalence, and host density from the literature and supplemented them with our own, new data on prevalence and shoaling rate. We used mixed models to test if guppy shoaling behaviour, *Gyrodactylus* spp. prevalence, and guppy density differed between predation regimes. In these models we used predation regime as a fixed effect along with river and year as random effects. We only included rivers with paired estimates from high and low predation populations because of variation across rivers. Each population average for a given site+year was one data point in these analyses. For % time shoaling (22 estimates, 4 rivers; *26, 57-59*) and prevalence (107 estimates, 11 rivers; *28, 31, 52, 53, 60-62*), we used a generalized linear mixed model beta regression with a logit link. For host density (23 estimates, 4 rivers; *63, 64-66*), we used a linear mixed model.

### 5. Animal use ethics statement

All collections and fish handling protocols were approved by the University of Pittsburgh’s Institutional Animal Care and Use Committee: protocol 18072155 and 21069471. The permit to collect and export guppies was granted by the Director of Fisheries in the Ministry of Agriculture, Land and Fisheries Division, Aquaculture Unit of the Republic of Trinidad and

Tobago on 2/26/2020 (copy available on request). The United States Import/Export License for live and preserved fish was issued by the U.S. Fish and Wildlife Service, Office of Law Enforcement Permit number A107080 on February 21, 2020 for the shipment of live guppies to the University of Pittsburgh. Samples were declared and cleared via USFWS form 3-177 on 03/11/2020.

### Supplementary Text

### 1. Supplementary equations

Host and parasite each evolve until selection favors neither a decrease nor an increase in the value of the evolving trait; at such “singular points”, the effect of an invader’s trait on the invader’s fitness (“selection gradient”, abbreviated *G* here) is zero. The relative simplicity of *G_T_* (*G* for parasite transmissibility) provides insight into parasite evolution (eq. S4A derived from eq. S2):

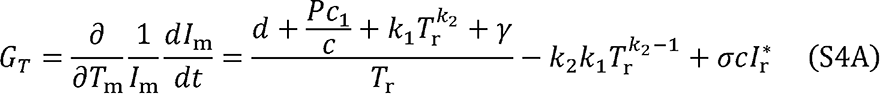

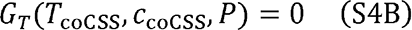

Eq. S4B holds at all singular points; CSS and coCSS points are subsets of singular points. Since *G_T_* is zero, the derivatives sum to zero (eq. S5A), allowing disentangling the consumptive and non-consumptive effects of predation on parasite evolution.

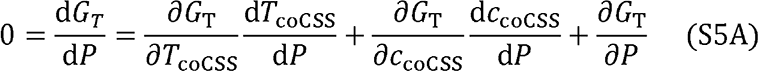

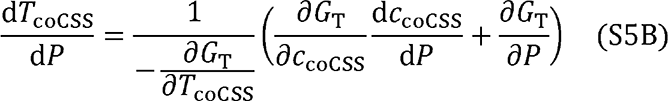

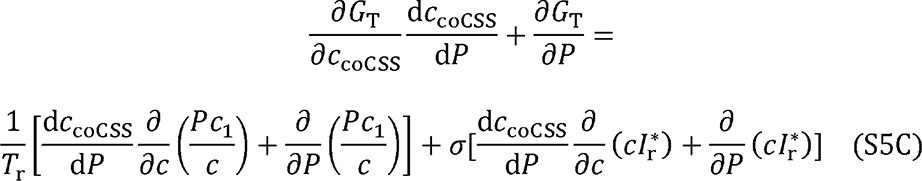

Note that ∂ *T*_coCSS_ < 0 by the evolutionary stability condition of Adaptive Dynamics, so 1/(-∂ *T*_coCSS_) acts as a positive scaling factor in eq. S5B. Eq. S5C shows the consumptive and non-consumptive effects of predation in four terms on the right-hand-side. The first term finds the non-consumptive effect of predation on shoaling rate, thus on non-parasite induced mortality, and thus on parasite evolution (pathway 3 in Fig. 1). The second term finds the consumptive effect of predation on non-parasite induced mortality and thus on parasite evolution (pathway 1). The third term finds the non-consumptive effect of predation on shoaling rate, thus on multiple infections, and thus on parasite evolution (pathway 4). The fourth term finds the consumptive effect of predation on multiple infections and thus on parasite evolution (pathway 2). It is trivial to convert pathway strengths for d*T*_coCSS_/d*P* to d*v*_coCSS_/d*P*, given the trade-off. Thus, this model allows partitioning of the strengths by which predation consumptively and non-consumptively affects parasite evolution.

This same reasoning also shows how predation and host shoaling rate influence virulence evolution by influencing infectious period. From eq. S4A, we derive eqs. S6A and S6B.

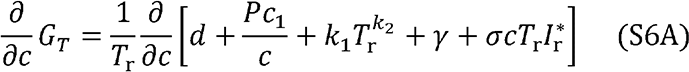

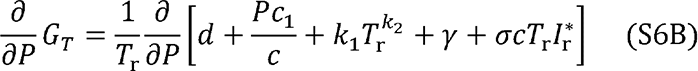

Infected host density of the resident genotype (*I*_r_*\**) is not expanded because its expression is long.

Infections for an invading genotype are lost at rate *d+Pc*_1_*/c+v+ + cT*_r_*I*_r_*\**; thus, the effect of γ σ predation and shoaling rate on *G_T_* is proportional to their effect on the rate at which infections are lost. From eq. S5B (or a similar expression, e.g., if *c* is simply a fixed parameter that is influencing parasite evolution), change in *T*_coCSS_ is proportional to change in *G_T_*. Thus, the effect of predation or shoaling rate on the rate of infection loss captures their effect on transmissibility and virulence evolution. This derivation holds for all parameters not part of the parasite trait tradeoff (i.e., not *k*_1_ or *k*_2_).

### 2. Identifying parasites to species

For the first, sequencing method for species ID, we extracted parasite DNA from three individuals per line with the Qiagen DNeasy Blood and Tissue (Qiagen, Inc., Valencia, CA). We amplified a 262 bp fragment of the mitochondrial COII gene following Xavier et al. (*67*) with the addition of a final extension step of 72 C for 7 min. Amplified DNA from 5 five domestic lines were sent to the DNA Analysis Facility on Science Hill at Yale University (New Haven CT, USA) for Sanger sequencing. Consensus sequences of reads in both the forward and reverse directions were assembled and edited manually in Sequencher 3.0 (Gene Codes Corporation, Ann Arbor, Michigan, U.S.). Sequences were aligned to reference sequences obtained from GenBank (*G. turnbulli* GenBank accession KP164811, *G. bullatarudis* GenBank accession KP168403) in MEGA-X 10.0.5 (*68*).

The second method was a PCR-restriction fragment length polymorphism (RFLP) assay we developed to differentiate between the three *Gyrodactylus* species that commonly infect guppies in Trinidad. This assay uses SspI-HF® (NEB) enzyme to cut the amplified portion of the mitochondrial COII gene differently depending on the species: *G. turnbulli* remains uncut, *G. bullatarudis* has 2 bands approximately 190 and 80bp, and *G. poeciliae* has 3 bands of approximately 140, 70 and 40bp. The restriction reaction followed manufacturer recommendations with 10ul of PCR product incubated for 12 hours at 37 °C. Bands were then visualized using a 4% agarose gel. Control *G. turnbulli* and *G. bullatarudis* PCR products, identified by Sanger sequencing, were used during the development of the assay and for every digest to ensure quality enzyme activity.

### 3. Parasite single nucleotide polymorphism (SNP) genotyping and analysis

We distinguished parasite genotypes using SNPs. To identify potential SNP loci, we used the resequencing data in Konczal et al. (*69, 70*) from: six *G. turnbulli* from each of the Aripo and Lopinot Rivers, and two from the Caura River; and two *G. bullatarudis* samples from the Aripo River, seven from the Lopinot, three from the Caura, and eight from the Santa Cruz River. Raw reads for *G. turnbulli* were mapped to a *G. turnbulli* reference genome derived from a domestic collection (J. Stephenson, unpublished data) and *G. bullatarudis* to the reference genome (GenBank accession no: GCA_012064415.1) using bwa mem with the default parameters.

Duplicates were marked using Picardtools. SNP calling was performed for the 100 longest scaffolds using samtools mpileup. SNPs within 5 bp of an indel, within 250 bp of another SNP, with quality scores below 100, or with mean mapping quality lower than 30 were excluded from subsequent analysis. Biallelic SNPs that were heterozygous in both Lopinot and Aripo (but with observed heterozygosity lower than 75%) were selected for subsequent primer design.

Primer design for the potential SNP loci was conducted by Fluidigm following their best practice recommendations. Two custom 96.96 dynamic arrays for each species were screened for reliable amplification and consistent SNP calling for 96 individual parasites from both wild and domestic parasites. PCR was performed on a Fluidigm 96.96 Dynamic Array (SNP chip) with the following PCR cycling conditions: 50 °C for 2 min, 70 °C for 30 min, 25 °C for 10 min and 95 °C for 5 min, followed by four touchdown cycles (95 °C for 15 s, from 64 °C to 61 °C for 45 s, 72 °C for 15 s) 34 additional cycles (95 °C for 15 s, 60 °C for 45 s, 72 °C for 15 s), and a final cycle at 20 °C for 10 s.

### 4. Data extraction from the literature

We extracted site-level measurements of guppy shoaling behavior, *Gyrodactylus* spp. prevalence, and guppy density from the literature using GoogleScholar. All searches included the terms “Trinidadian guppy*” and “Poecilia reticulata”. For shoaling estimates (Fig. 5F), we added “anti-predator response/behavior“, “shoal*“, “fusion-fission dynamics“, “social*“, “school*“, “upper“, “lower“, “high predation“, “low predation”. We targeted our focal rivers by adding “Aripo” and “Guanapo”. We recorded river, site, predation regime, and the metric the authors of the study used to estimate guppy social behavior, and any error estimate given. To obtain literature estimates of prevalence (Fig. 5G), we added the terms “Gyrodactylus“, “infection” and “prevalence“, and recorded the river, site, predation regime, and *Gyrodactylus* spp. prevalence among wild guppies, as well as year the estimates were taken, and any error estimates given. For host density (Fig. 5H), we added “density“, and recorded or extracted the estimates of the number of guppies per m^2^. Papers that were obtained for other site-level measurements (shoaling and prevalence) were also examined to see if they incidentally also recorded density. Papers that gave estimates of these metrics under controlled conditions for lab lines of guppies instead of wild guppies were excluded. Where researchers reported multiple estimates, we took the mean value of those measured in one location at one timepoint as a single, independent observation.

### 5. Supplementing data from the literature

In March 2020, we collected guppies from each of our four focal populations using seine nets, transported them to the field station and held them in pools (80 cm diameter) for 24 to 48hrs (mean = 33.6hrs) before assaying their shoaling tendency. We used a standard assay to assess the shoaling tendency of individual fish (*71*) and recorded each trial using GoPro cameras (HERO4). We allowed fish to acclimatize for 10 minutes before removing a partition separating the focal fish from a shoal tank (containing three non-focal individuals from the same population), waited 200s after this removal, and then recorded the proximity of the focal fish to the shoal for a further 10 minutes using Boris (v7.9.8; *72*). All assays and behavioral recordings were conducted by a single observer blind to infection status and predation regime. We removed some fish from the analysis because of camera error, or because the fish did not move for the trial duration. Our final sample sizes for each population were: low-predation Guanapo: 4 females, 5 males; high-predation: 3 females, 3 males; low-predation Aripo: 14 females, 13 males; high-predation: 12 females, 14 males. We recorded fish weight, length, sex, and presence of *Gyrodactylus* spp. infection after its behavior assay, and tested for their effects, along with fish predation regime and river of origin, on % time shoaling using a generalized linear mixed model (Beta error, logit link function). Date of trial, lighting level, and behavior enclosure were included as random effects.

We collected prevalence estimates in March 2020 by catching between 50 and 30 fish per site with a 1 m x 1 m seine net, and screening those from the Caroni drainage for parasites as above at a nearby field station.

---

**Figure S1.**
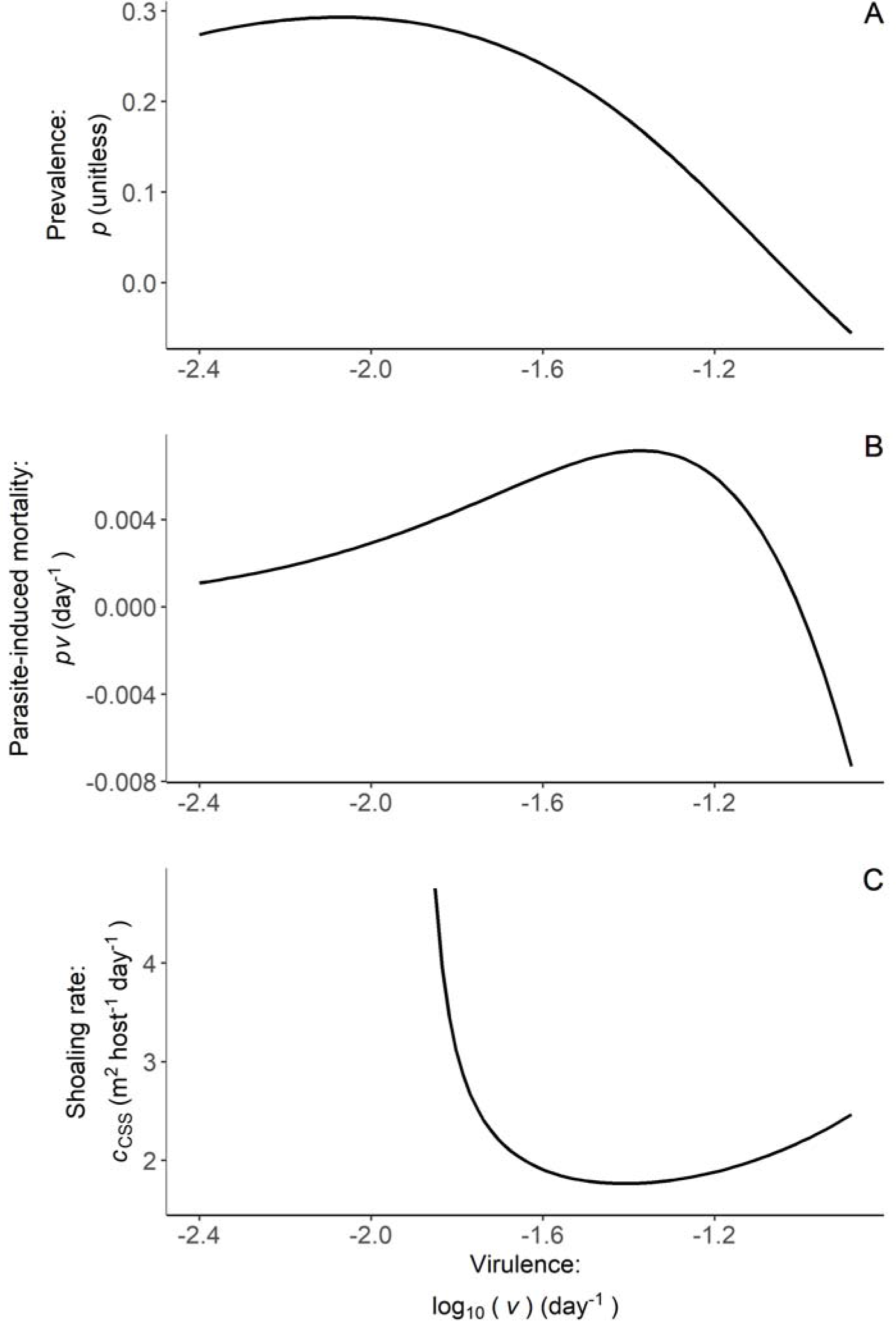
Virulence can select for increased contact rate. Hosts can evolve increasing contact rate in response to increased virulence, especially at very high virulence (beyond range used in main text). (**A**) Increasing virulence (and transmissibility along the trade-off) can decrease prevalence. (**B**) Overall parasite-induced mortality can decrease if prevalence declines sharply enough. This decrease occurs because, while parasites are very virulent, very few hosts are infected and suffering that virulence. (**C**) At high virulence, increasing virulence can select for higher host contact rates. *c* = 2 used for (A) and (B); *P* = 0.074 used for (C). All other parameters at default (Table 1).

**Figure S2.**
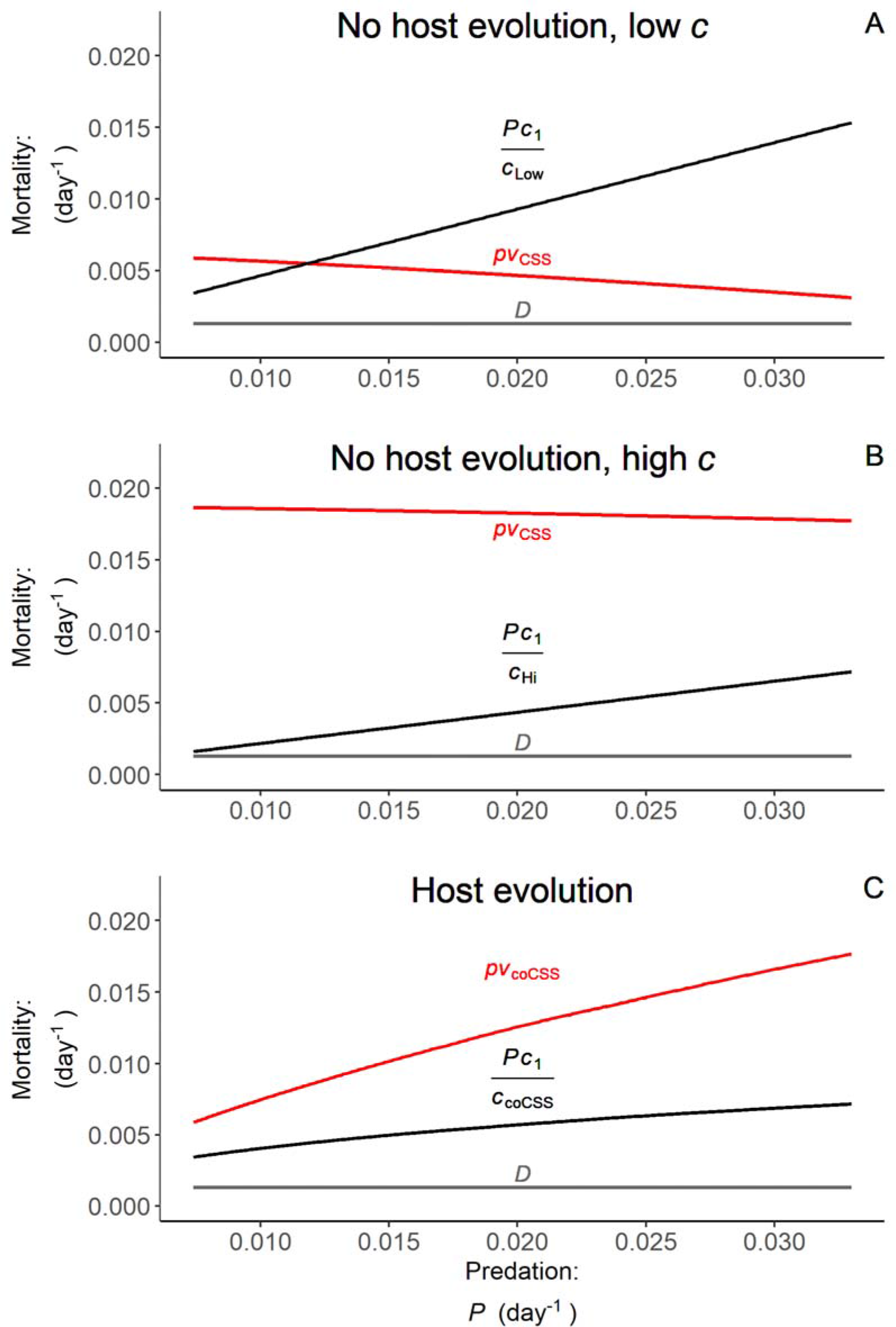
Host evolution in response to increasing predation causes parasite-induced mortality (red curves) to increase more than predator-induced mortality (black curves). (A) Without host evolution (contact rate, c, set to the green point in Fig. 4 while parasites evolve to some CSS), parasite-induced mortality declines with predation while predator-induced mortality increases. Death from background sources (*d*, grey line) does not change. (**B**) This trend is similar for a higher *c* (set to high, blue point in Fig. 4). (**C**) When hosts evolve increasing *c* with increasing predation (coCSS curve connecting green and blue points in Fig. 4), parasite-induced mortality increases more than predator-induced mortality. This pattern is due to host evolution and is qualitatively unchanged if hosts evolve but parasites do not.

**Figure S3.**
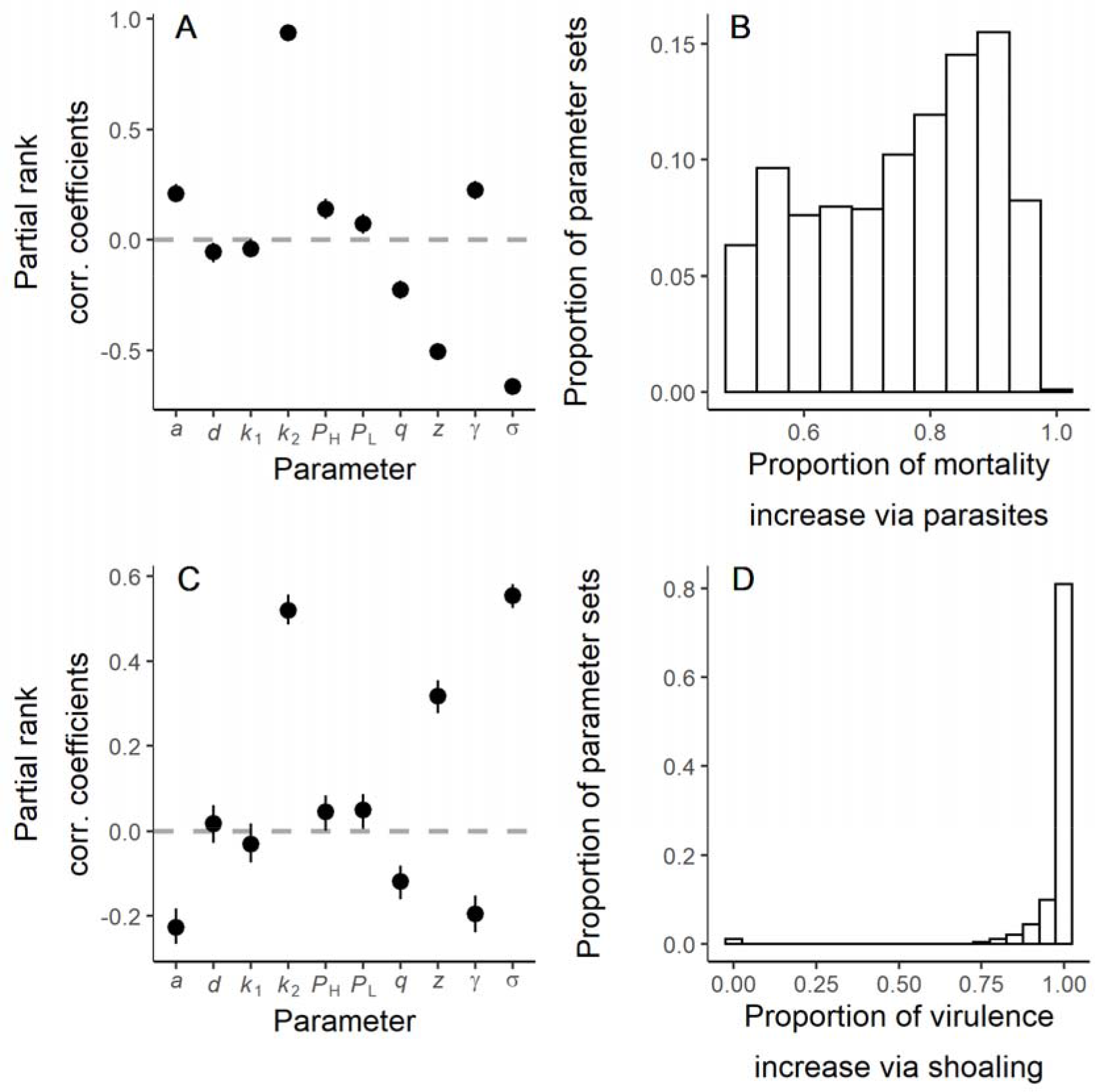
Sensitivity analysis shows the impact of parameters on key outputs and the range for those outputs. Key model outputs: the proportion of the mortality increase from increasing predation that is via increased parasite-induced mortality (**A-B**) and the proportion of the virulence increase from increasing predation that is via non-consumptive effects (**C-D**). Vertical lines are 95% confidence intervals in **(A & C)**. (A) Notable among other parameters, the trade-off exponent, *k*_2_, strongly increases the proportion of mortality increase due to parasites. Note here that *P*_H_ = *P*_Hi_ and *P*_L_ = *P*_Low_. (B) The proportion of mortality increase via parasites is typically well over half (0.5). (C) Both *k*_2_ and the superinfection parameter, , strongly increase σ the proportion of virulence increase via non-consumptive effects. (D) The proportion of virulence increase via shoaling (i.e., non-consumptive effects of predators) is typically well over half. This proportion was less than 0.7 in only 1.10% of parameter sets because the net, non-consumptive effects of predators on selection for virulence were negative in those parameter sets.

**Figure S4.**
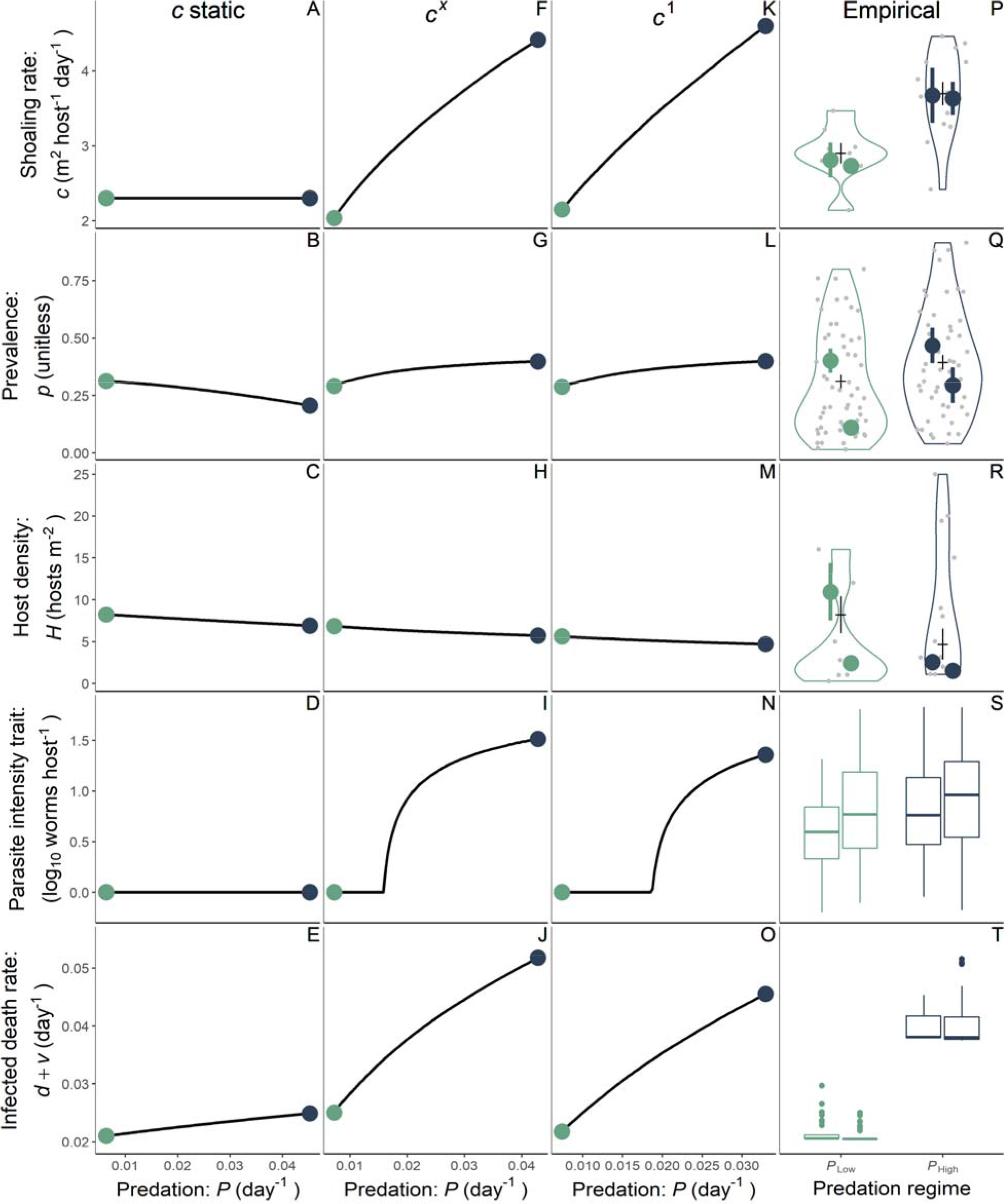
Alternate model fits to and predictions of data. Each of three models were allowed to fit as well as possible to the training data, e.g., by shifting the values of *P*_Low_ and *P*_Hi_. (**A-E**) The model with *c* fit as a static parameter, instead of output from host evolution, fits poorly to the data. Notably, it is a poor fit to prevalence, with prevalence decreasing with predation. (**F-J**) The model with an exponent governing the benefit of shoaling, *c^x^*, fits the training data well but has little difference from the simpler model where the benefit of shoaling is simply *c*, *x* = 1.47 *vs. x* = 1 in simpler model. (**K-O**) Accounting for model complexity, this model is best fits the training data such as (**P-R**) and thus is the focal model presented throughout. The appropriateness of this model is verified by the fact that it is also best at predicting the validating data (**S-T**).

**Table S1.** Coinfection rates in the wild. For each site (river+predation regime), we genotyped multiple worms per a sample of fish hosting more than one worm. Coinfection within species = (no. fish with >1 genotype of species x/no. fish with > 1 worm of species x) × % of infections with >1 worm. Overall coinfection = (no. fish with at least two different worms/no. fish with at least two worms) × % of infections with >1 worm. Infections were diverse overall. Coinfection by identical multilocus genotypes was rare (31 out of the 73 worms possible). G.t. = *G. turnbulli* and G.b. = *G. bullatarudis*.

| Site |  | Sample size |  | Species presence on fish |  |  | Infections >1 worm | Coinfection within species |  | Overall coinfection rate |
| --- | --- | --- | --- | --- | --- | --- | --- | --- | --- | --- |
| River | Predation | Fish | Worms/fish<br>(mean±SE) | %G.t.<br>only | %G.b.<br>only | %Both | % | G.t. | G.b. | % |
| Aripo | High | 7 | 2.43±0.20 | 14.3 | 85.7 | 0 | 70.8 | 0 | 35.4 | 30.3 |
| Aripo | Low | 4 | 3.25±0.75 | 100 | 0 | 0 | 58.2 | 14.6 | NA | 14.6 |
| Guanapo | High | 7 | 2.29±0.18 | 14.3 | 85.7 | 0 | 62.5 | 62.5 | 52.1 | 53.6 |
| Guanapo | Low | 5 | 2 | 80 | 20 | 0 | 61.9 | 15.5 | 0 | 12.4 |
| Santa Cruz | High | 7 | 4.86±1.30 | 0 | 71.4 | 28.6 | 92.6 | 92.6 | 92.6 | 92.6 |
| Santa Cruz | Low | 3 | 3±1 | 66.7 | 33.3 | 0 | 61.7 | 30.9 | 61.7 | 41.1 |

**Table S2.** Threshold loci numbers. For each source population/species combination, a number of variable loci was found. To be considered a unique genotype, an individual had to differ from all other genotypes at half or more of the variable loci for that population. Some source populations only had one individual of a given species and thus have “NA” listed.

| Source population | Species | Number of variable loci<br>within source | Number of loci threshold |
| --- | --- | --- | --- |
| $P_{Hi}$ Aripo | *G. bullatarudis* | 37 | 19 |
| $P_{Hi}$ Aripo | *G. turnbulli* | NA | NA |
| $P_{Low}$ Aripo | *G. bullatarudis* | NA | NA |
| $P_{Low}$ Aripo | *G. turnbulli* | 87 | 44 |
| $P_{Hi}$ Guanapo | *G. bullatarudis* | 22 | 11 |
| $P_{Hi}$ Guanapo | *G. turnbulli* | 46 | 23 |
| $P_{Low}$ Guanapo | *G. bullatarudis* | 17 | 9 |
| $P_{Low}$ Guanapo | *G. turnbulli* | 77 | 39 |
| $P_{Hi}$ Santa Cruz | *G. bullatarudis* | 41 | 21 |
| $P_{Hi}$ Santa Cruz | *G. turnbulli* | 23 | 11 |
| $P_{Low}$ Santa Cruz | *G. bullatarudis* | 19 | 9 |
| $P_{Low}$ Santa Cruz | *G. turnbulli* | 24 | 12 |
| $P_{Hi}$ Lopinot | *G. turnbulli* | 34 | 17 |
| Commercial Source 1 | *G. turnbulli* | 31 | 16 |
| Commercial Source 2 | *G. turnbulli* | NA | NA |
| Commercial Source 3 | *G. turnbulli* | NA | NA |

**Table S3.**
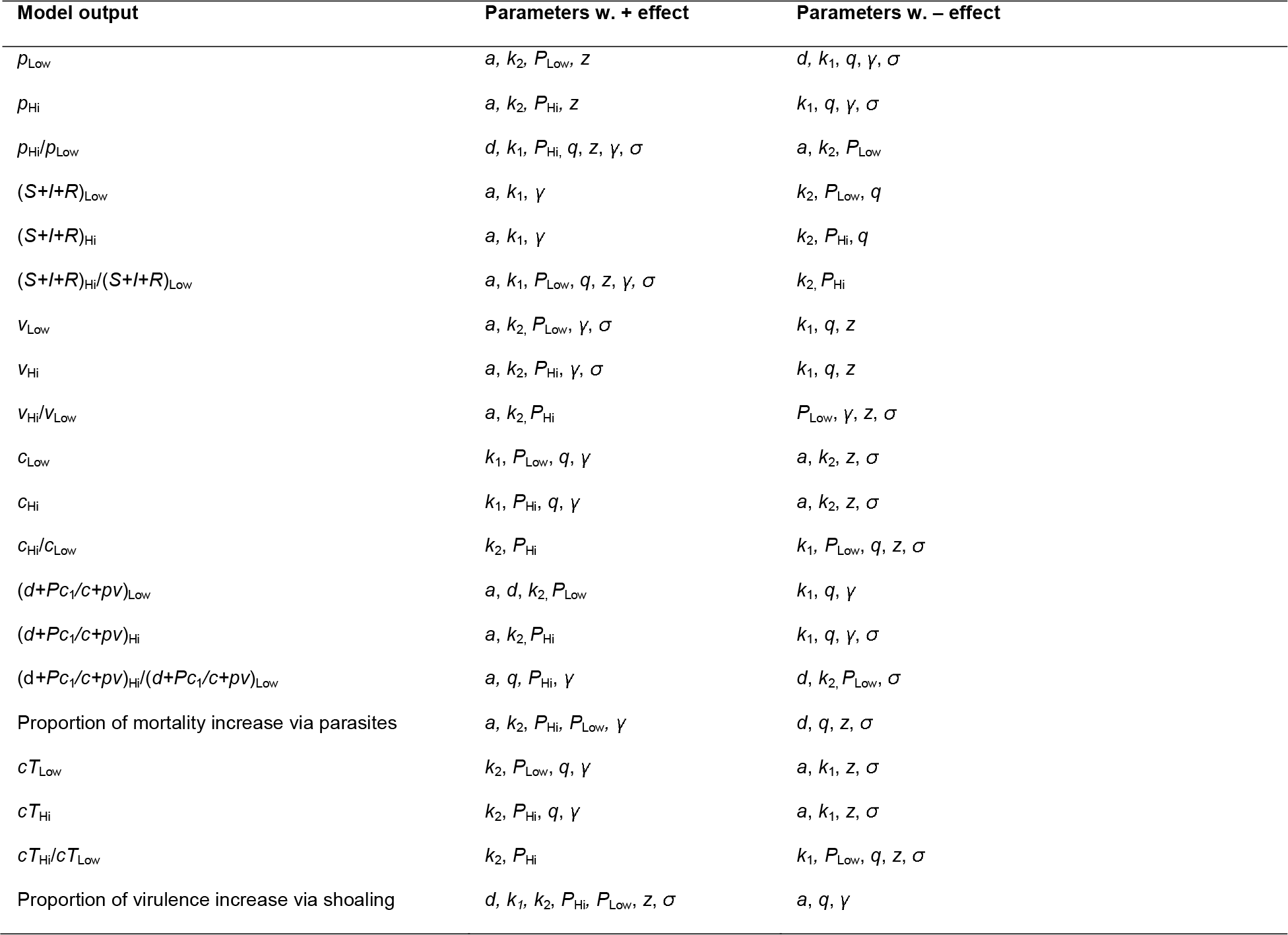
Parameter effects based on 95% CIs. For each model output (first column), we report which parameters increased (second column) or decreased (third column) that model output in the sensitivity analysis. A positive or negative effect is only reported if the 95% CI for that parameter’s effect did not overlap 0. Trait outputs are the coCSS trait values.

