## Supplementary Code for "Social hosts evade predation but have deadlier parasites"

**3 Testing model predictions of parasite intensity and virulence** **20**

3.1 Testing for parasite intensity differences among lines from our four focal populations - Fig. 5I 20

3.2 Testing for host death rate differences among lines from our four focal populations - Fig. 5J . 29

**4 Coevolutionary model - Fig. 4** **35**

### 1 Estimating the virulence-transmission tradeoff

#### 1.1 Load necessary packages

```
# Clear the working environment rm(list=ls())

# Load necessary libraries
require(dplyr)
require(plyr)
require(DHARMa)
require(effsize)
require(effectsize)
require(glmmTMB)
require(ggplot2)
require(car)
require(visreg)
require(lme4)
require(gridExtra)
require(formatR)
require(tidyverse)
require(Rmisc)
require(MuMIn)
require(effects)
require(egg)
require(pracma)
require(SuppDists)
require(ggpubr)
require(Deriv)
require(rootSolve)
require(styler)
require(lhs)
require(sensitivity)
require(knitr)
```

#### 1.2 Examining the relationship between intensity and transmission - Fig. 2A

Here we use data from Stephenson et al 2017 and an additional, unpublished transmission experiment. The dataset is 'SI_transmission.csv', which contains the following variables:

**full_dwormattrans**: the number of parasites the donor had at the point of transmission
**full_rspeedtrans**: the transmission rate, calculated following the details in the main text
**full_line**: the identity of the isogenic line: Gt3 is the line used by Stephenson et al 2017, the others are previously unpublished data

```
dtrans=read.csv("SI_transmission.csv")

#Full model
transpeed<-glmmTMB(full_rspeedtrans~
                        full_dwormattrans+full_line,
                   data=dtrans,family=Gamma(link=log))

Anova(transpeed,type="II")
```

```
## Analysis of Deviance Table (Type II Wald chisquare tests)
##
## Response: full_rspeedtrans
##                      Chisq Df Pr(>Chisq)
## full_dwormattrans 7.2053  1   0.007269 **
## full_line         1.8859  3   0.596416
## ---
## Signif. codes:  0 '***' 0.001 '**' 0.01 '*' 0.05 '.' 0.1 ' ' 1

#Estimating effect size
#chisq_to_phi(1.8859,dim(dtrans)[1])

#Diagnostic plots
sim_residuals_trans <- simulateResiduals(transpeed, 1000)
plot(sim_residuals_trans)
```

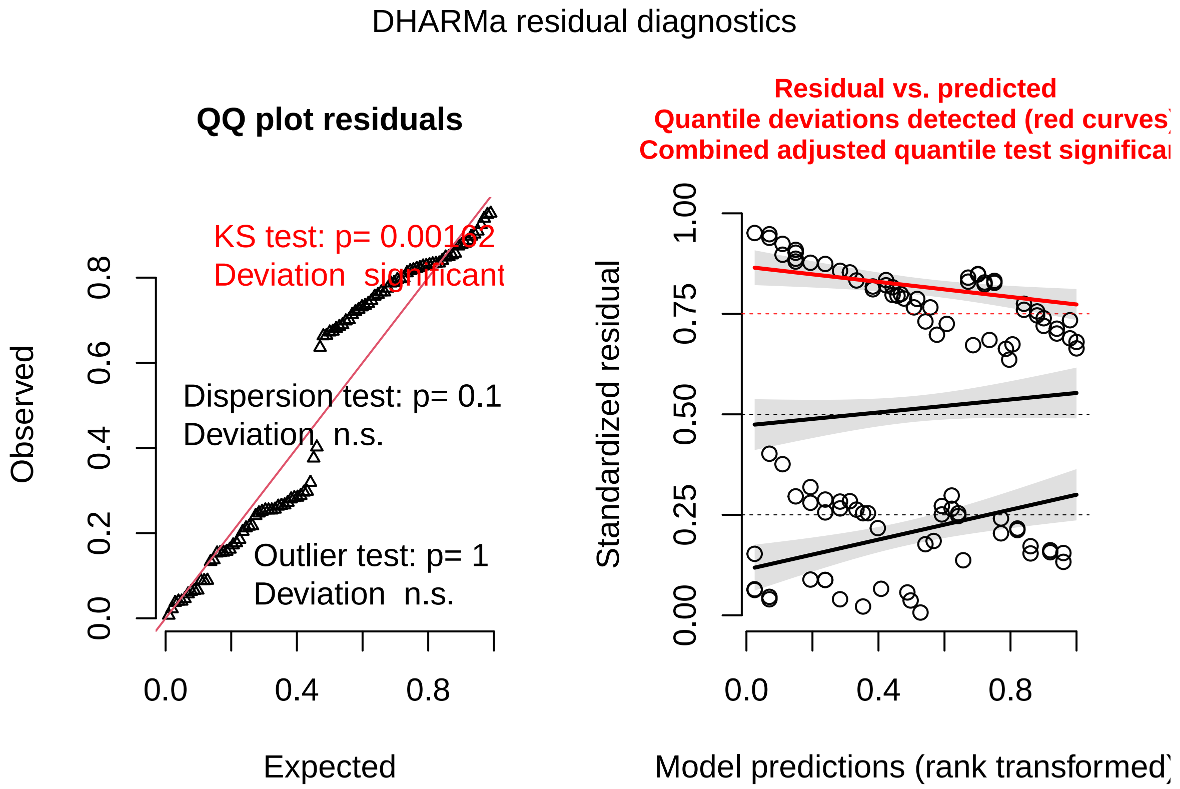

```
testDispersion(sim_residuals_trans)
```

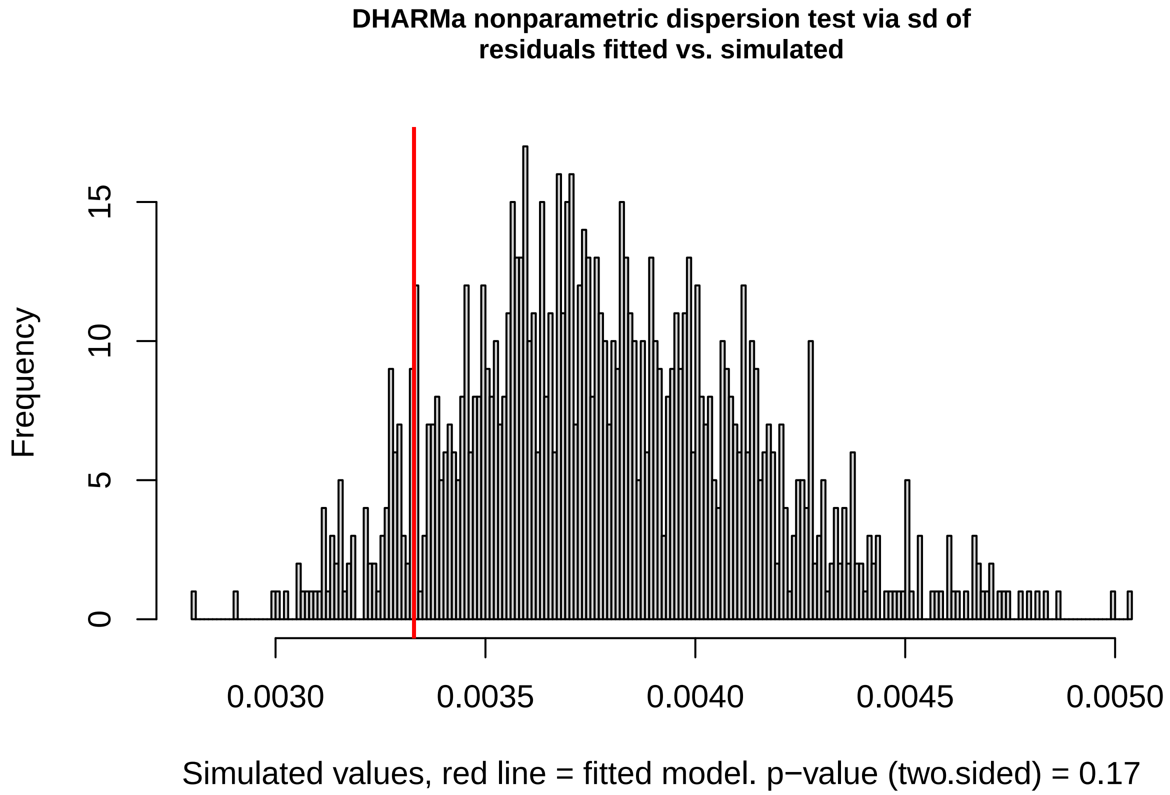

```
##
## DHARMa nonparametric dispersion test via sd of residuals fitted vs.
   simulated
##
## data: simulationOutput
## ratioObsSim = 0.87762, p-value = 0.17
## alternative hypothesis: two.sided

#Eliminating the non-significant parasite line term here
#to improve our ability to predict across lines

transpeed2<-glmmTMB(full_rspeedtrans~
                      full_dwormattrans,
                    data=dtrans,family=Gamma(link=log))

#Estimating effect size
#parameters::model_parameters(transpeed2)
#z_to_r(4.14,dim(dtrans)[1])

#summary(finalspeedmod)
Anova(transpeed2,type="II")

## Analysis of Deviance Table (Type II Wald chisquare tests)
##
## Response: full_rspeedtrans
##                    Chisq Df Pr(>Chisq)
## full_dwormattrans 17.135  1  3.481e-05 ***
```

```
## ---
## Signif. codes:  0 '***' 0.001 '**' 0.01 '*' 0.05 '.' 0.1 ' ' 1

#Diagnostic plots
finalspeedmod_sim<- simulateResiduals(transpeed2, 1000)
plot(finalspeedmod_sim)
```

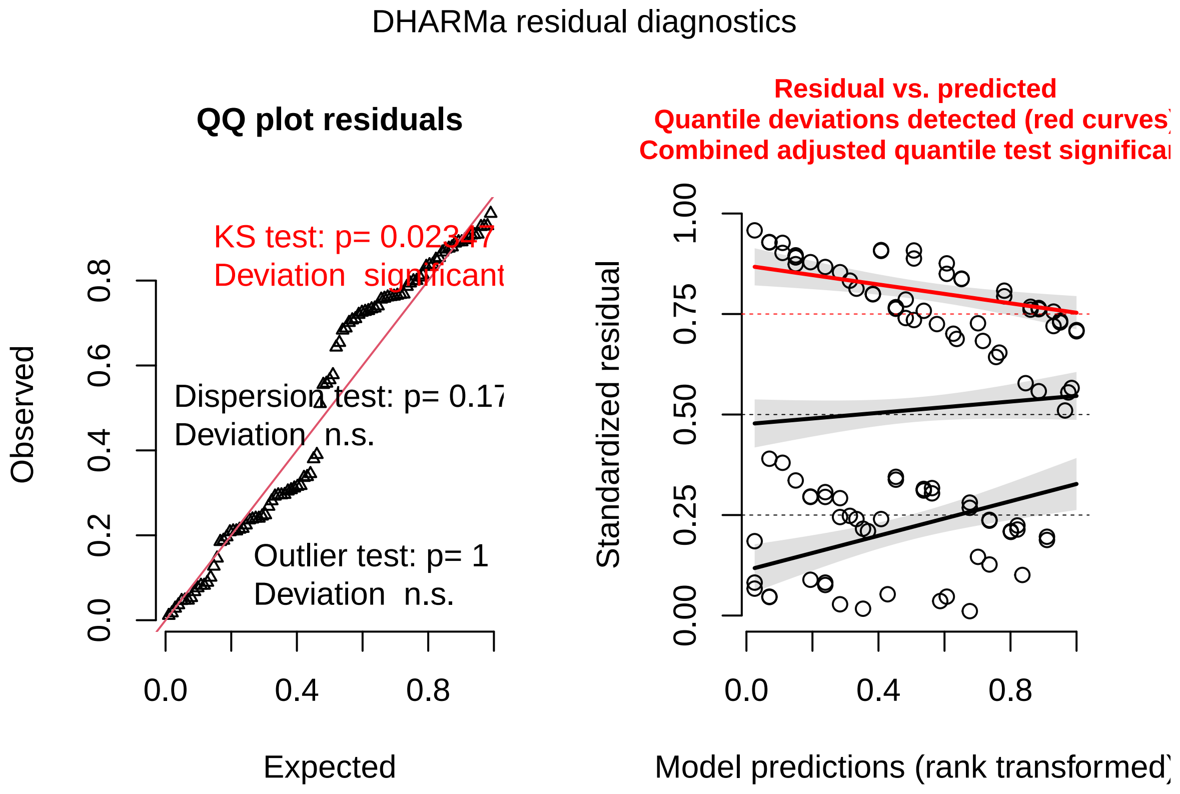

```
testDispersion(finalspeedmod_sim)
```

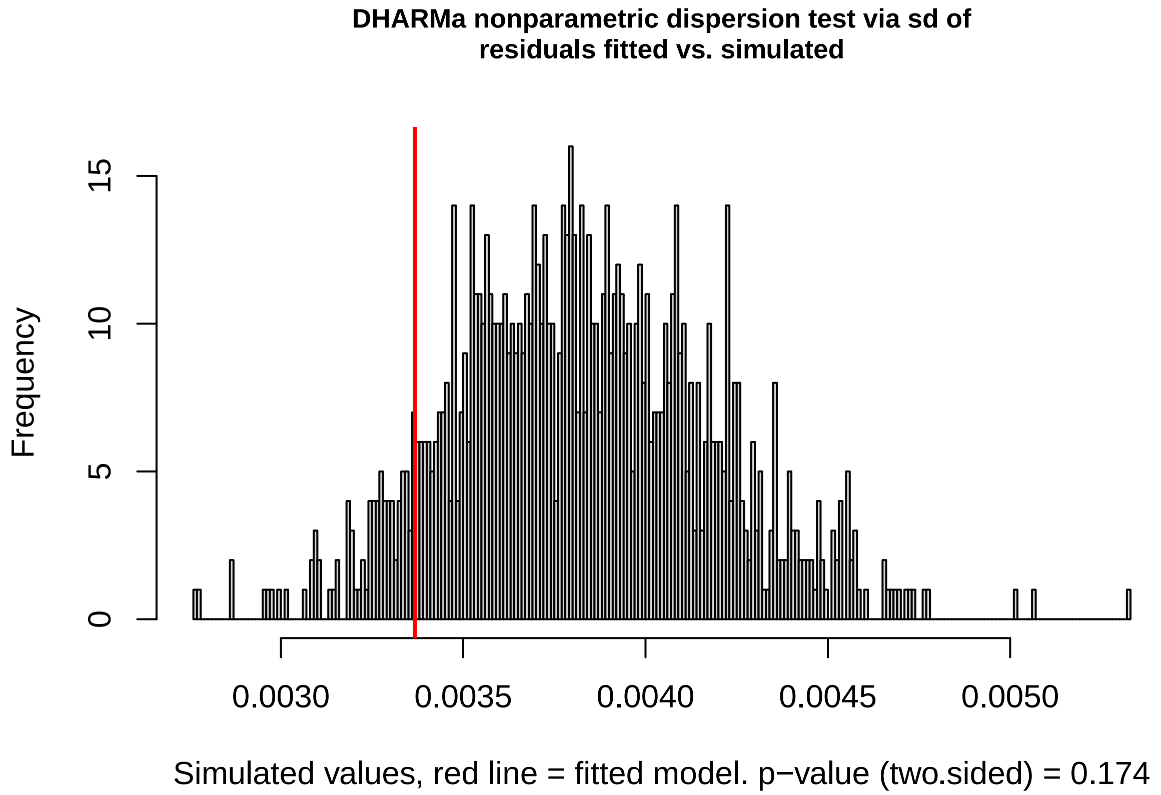

```
##
## DHARMa nonparametric dispersion test via sd of residuals fitted vs.
    simulated
##
## data:  simulationOutput
## ratioObsSim = 0.88041, p-value = 0.174
## alternative hypothesis: two.sided
```

##### 1.3 What is the relationship between intensity and host death in lab maintained lines? Fig. 3B

Here we use data on the average intensity and average host death rate from isogenic lines maintained separately, but on the same stock of fish and under the same conditions in the lab. The data are provided in 'SI_tradeoff.csv', which contains the following variables:

**linename**: the identity of the isogenic line
**meanintensity**: the mean intensity recorded on fish infected with that line
**meandeath**: the mean death rate of fish infected with that line
**meantransmission**: the mean transmission rate of each isogenic line, estimated from their mean intensities and the relationship in Fig. 3A
**origin**: whether the isogenic line originated from a guppy collected in the wild ('Wild'), or obtained from a commercial supplier ('Domestic')

```
# Update here
dto <- read.csv("SI_tradeoff.csv")
```

```
tradeoff_model <- glmmTMB(meandeath ~ meanintensity + origin,
    data = dto, na.action = na.omit, family = beta_family(link = "logit"))

# summary(tradeoff_model)
Anova(tradeoff_model, type = "II")

## Analysis of Deviance Table (Type II Wald chisquare tests)
##
## Response: meandeath
##                  Chisq Df Pr(>Chisq)
## meanintensity  6.9665  1  0.0083050 **
## origin        14.7296  1  0.0001241 ***
## ---
## Signif. codes:  0 '***' 0.001 '**' 0.01 '*' 0.05 '.' 0.1 ' ' 1

# Estimating effect size
# parameters::model_parameters(tradeoff_model)
# z_to_r(2.64,dim(dto)[1]) z_to_r(3.84,dim(dto)[1])

# Diagnostic plots
sim_residuals_tradeoff <- simulateResiduals(tradeoff_model, 1000)
plot(sim_residuals_tradeoff)
```

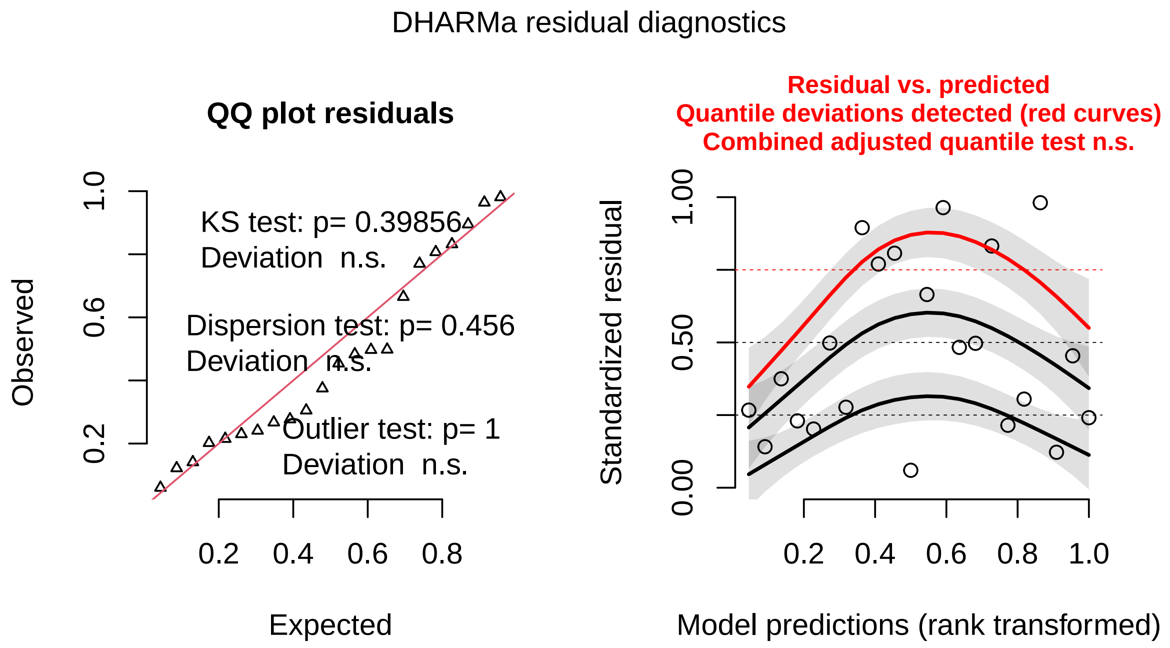

```
testDispersion(sim_residuals_tradeoff)
```

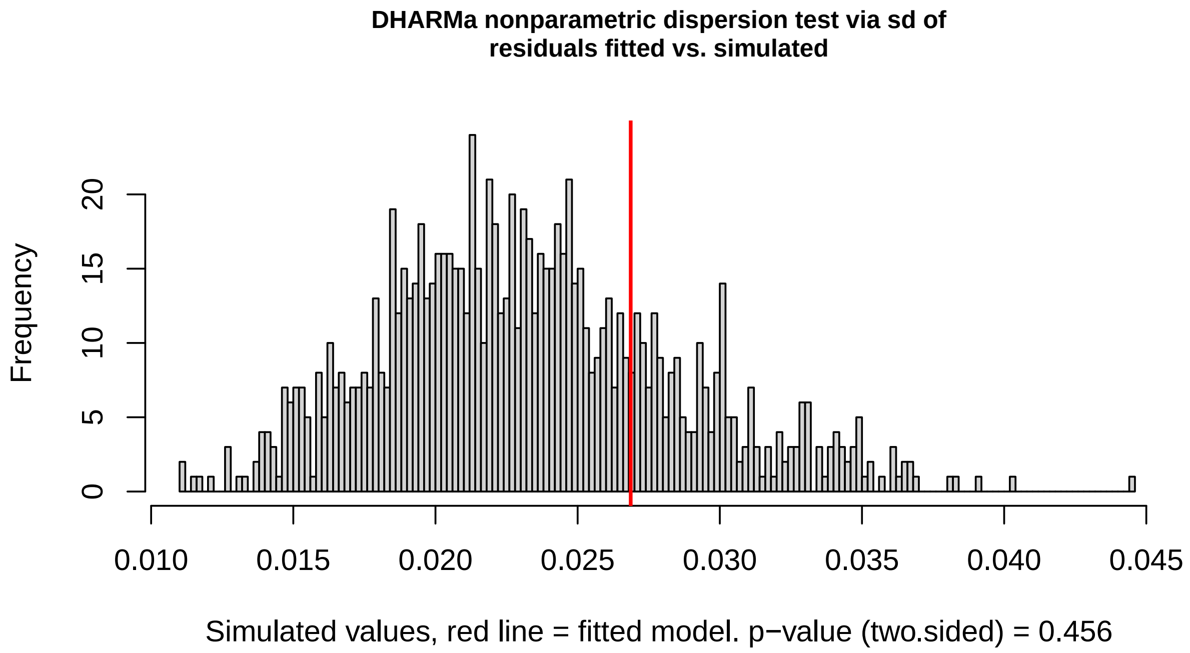

```
##
## DHARMa nonparametric dispersion test via sd of residuals fitted vs.
   simulated
##
## data:  simulationOutput
## ratioObsSim = 1.154, p-value = 0.456
## alternative hypothesis: two.sided
```

##### 1.4 What is the relationship between death rate and transmission rate? Figure 3C

```
# Save the partial residuals from tradeoff_model
myv <- visreg(tradeoff_model, "meanintensity", scale = "response",
    cond = list(origin = "Wild"), plot = F)

# If instead we use partial residuals as if all lines were
# domestic, the tradeoff becomes v_i = 2.80*10^6*beta_i^4.03,
# which still has a stabilizing curvature (exponent > 1) and
# predicted v_CSS at low predation is 0.021 while at high
# predation it is 0.046
myv2 <- visreg(tradeoff_model, "meanintensity", scale = "response",
    cond = list(origin = "Domestic"), plot = F)
myvres <- myv$res
myvres2 <- myv2$res

dto$partialdeaths <- myvres$visregRes
dto$partialdeaths2 <- myvres2$visregRes

# This data relates partial residual mean death rate and mean
```

```
# transmission rate for a line. Given a background death rate
# value, this can be converted into partial residual mean
# virulence and mean transmission rate

# Need to use this to run evolutionary algorithm
curve_data <- cbind(dto$partialdeaths, dto$meantransmission)
curve_data2 <- cbind(dto$partialdeaths2, dto$meantransmission)

# Update here Estimates of background death rate from
# evolutionary algorithm - given below
d_ex <- 0.001297164
y_ex <- 0.02027253

# Subtract the estimate of background death rate to get
# virulence
dto$partialvirulence <- dto$partialdeaths - d_ex
dto$partialvirulence2 <- dto$partialdeaths2 - d_ex
# Make sure that every virulence is non-negative, which is
# usually the case anyway

# Statistics for Figure 3C
summary(Death_curve <- glm(partialvirulence ~ log(meantransmission),
    family = Gamma(link = "log"), data = dto))

##
## Call:
## glm(formula = partialvirulence ~ log(meantransmission), family = Gamma(link
    = "log"),
##     data = dto)
##
## Deviance Residuals:
##      Min        1Q    Median        3Q       Max
## -0.17068  -0.08869  -0.03863   0.04576   0.37131
##
## Coefficients:
##                       Estimate Std. Error t value Pr(>|t|)
## (Intercept)            14.1356     1.8686   7.565 2.73e-07 ***
## log(meantransmission)   3.6079     0.3941   9.155 1.37e-08 ***
## ---
## Signif. codes:  0 '***' 0.001 '**' 0.01 '*' 0.05 '.' 0.1 ' ' 1
##
## (Dispersion parameter for Gamma family taken to be 0.01887556)
##
##     Null deviance: 2.01103  on 21  degrees of freedom
## Residual deviance: 0.34296  on 20  degrees of freedom
## AIC: -154.03
##
## Number of Fisher Scoring iterations: 4

summary(Death_curve2 <- glm(partialvirulence2 ~ log(meantransmission),
    family = Gamma(link = "log"), data = dto))

##
## Call:
```

```
## glm(formula = partialvirulence2 ~ log(meantransmission), family = Gamma(
    link = "log"),
##     data = dto)
##
## Deviance Residuals:
##       Min        1Q    Median        3Q       Max
## -0.19023  -0.09830  -0.04320   0.04964   0.41045
##
## Coefficients:
##                         Estimate Std. Error t value Pr(>|t|)
## (Intercept)              14.7040     2.0772   7.079 7.31e-07 ***
## log(meantransmission)     3.9961     0.4381   9.121 1.45e-08 ***
## ---
## Signif. codes:  0 '***' 0.001 '**' 0.01 '*' 0.05 '.' 0.1 ' ' 1
##
## (Dispersion parameter for Gamma family taken to be 0.02332658)
##
##     Null deviance: 2.47344  on 21  degrees of freedom
## Residual deviance: 0.41965  on 20  degrees of freedom
## AIC: -205.62
##
## Number of Fisher Scoring iterations: 4

# Diagnostic plots
sim_residuals_curve <- simulateResiduals(Death_curve, 1000)
plot(sim_residuals_curve)
```

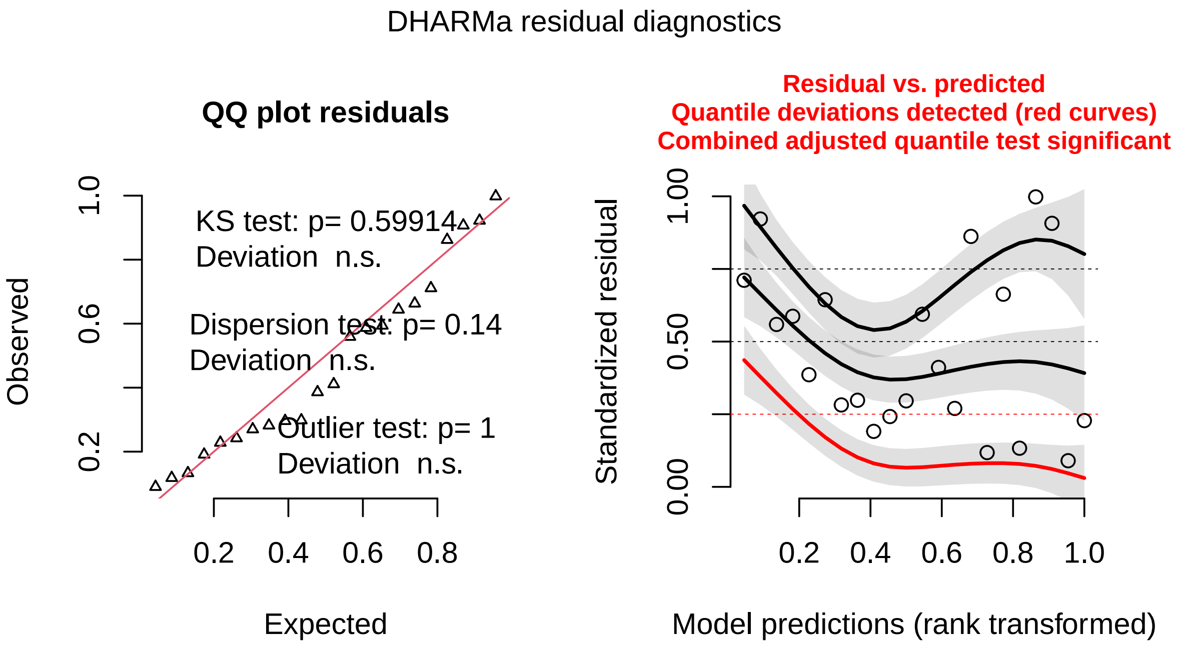

```
testDispersion(sim_residuals_curve)
```

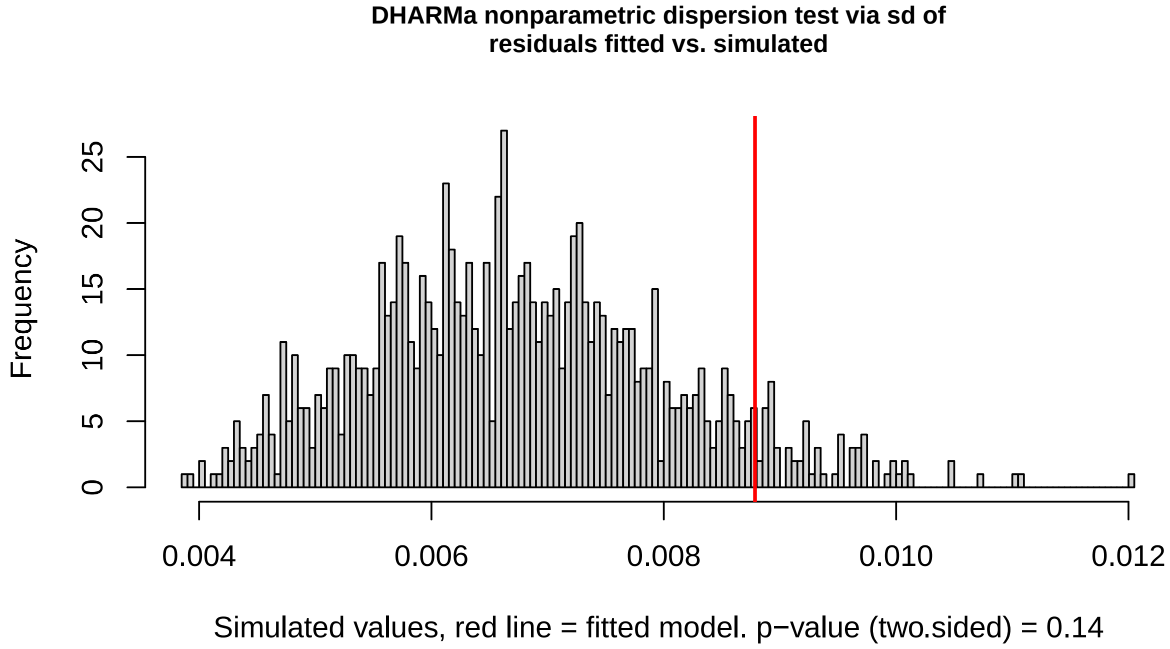

```
##
## DHARMa nonparametric dispersion test via sd of residuals fitted vs.
   simulated
##
## data: simulationOutput
## ratioObsSim = 1.3016, p-value = 0.14
## alternative hypothesis: two.sided
```

### 2 Empirical estimates of model parameters

#### 2.1 Guppy behaviour differs across river courses - Fig. 5F

Here we provide the code used to test for difference in proportion of time spent shoaling between guppies from the upper (low predation) and lower courses (high predation) of five Trinidadian rivers. These data come both from published estimates, and our own lab measurements, and are provided in the files 'SI_Behav.csv' (just our lab measurements) and 'Lit_behavior.csv' (all measurements). First, we present the analysis of our own measurements, which contain the following variables:

**Sex**: sex of the fish
**River**: the river of origin of the fish
**Course**: the course of the river of origin - upper or lower
**Length**: length of the fish (mm)
**Infec**: whether or not the fish was infected
**Prop.Shoal200**: the proportion of time the fish spent shoaling during a behavioral trial
**DOT**: date the data were collected in YYYYMMDD format
**ShoalID**: identity of the shoal used - each was used more than once
**ShoalSide**: side of the enclosure that contained the shoal - to control for side bias
**enclosure**: which enclosure was used for the trial
**Lighting**: whether the trial was conducted in relatively light or dark lighting

```
# Read in data collected in focal rivers, 2020
Behav_final <- read.csv("SI_Behav.csv")

# Our shoaling result
Our_behav <- glmmTMB(Prop.Shoal200 ~ Length + Course + River +
    Infec + Sex + (1 | DOT) + (1 | enclosure) + (1 | Lighting) +
    (1 | ShoalSide), Behav_final, family = beta_family(link = "logit"))
# summary(Our_behav)

# parameters::model_parameters(Our_behav)
# z_to_r(-2.77,dim(Behav_final)[1])

# P-value and test statistic
Anova(mod = Our_behav, type = "II", test.statistic = "Chisq")

## Analysis of Deviance Table (Type II Wald chisquare tests)
##
## Response: Prop.Shoal200
##         Chisq Df Pr(>Chisq)
## Length 0.0034  1   0.953516
## Course 7.6832  1   0.005574 **
## River  0.2129  1   0.644516
## Infec  0.0029  1   0.957183
## Sex    3.5680  1   0.058903 .
## ---
## Signif. codes:  0 '***' 0.001 '**' 0.01 '*' 0.05 '.' 0.1 ' ' 1

# Diagnostic plots
sim_residuals_behav <- simulateResiduals(Our_behav, 1000)
plot(sim_residuals_behav)
```

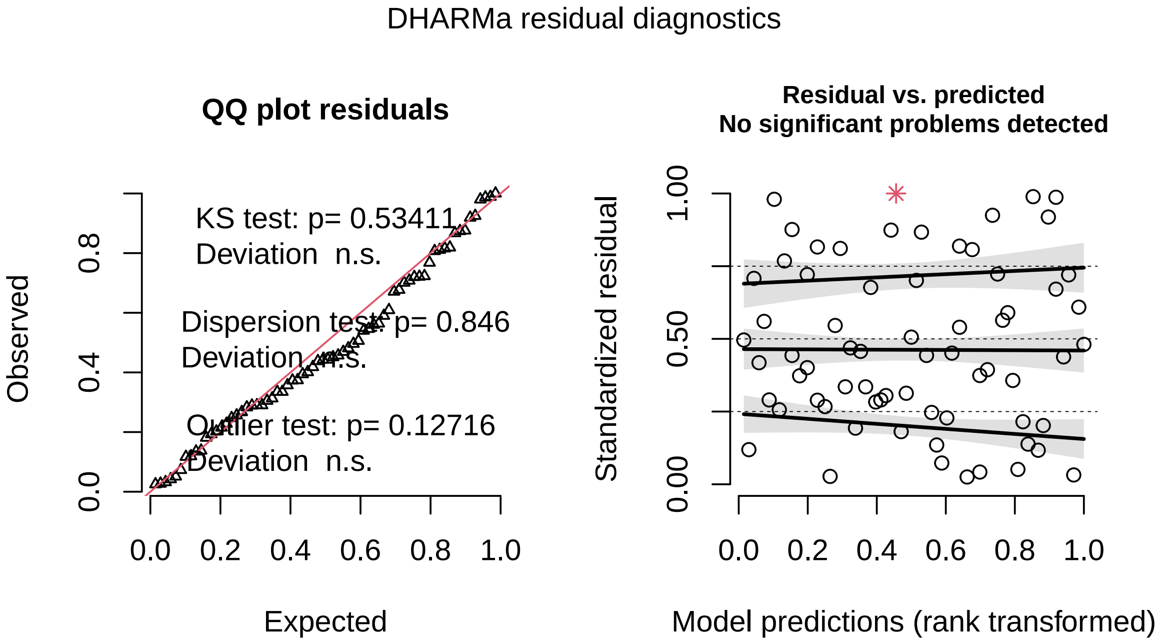

```
testDispersion(sim_residuals_behav)
```

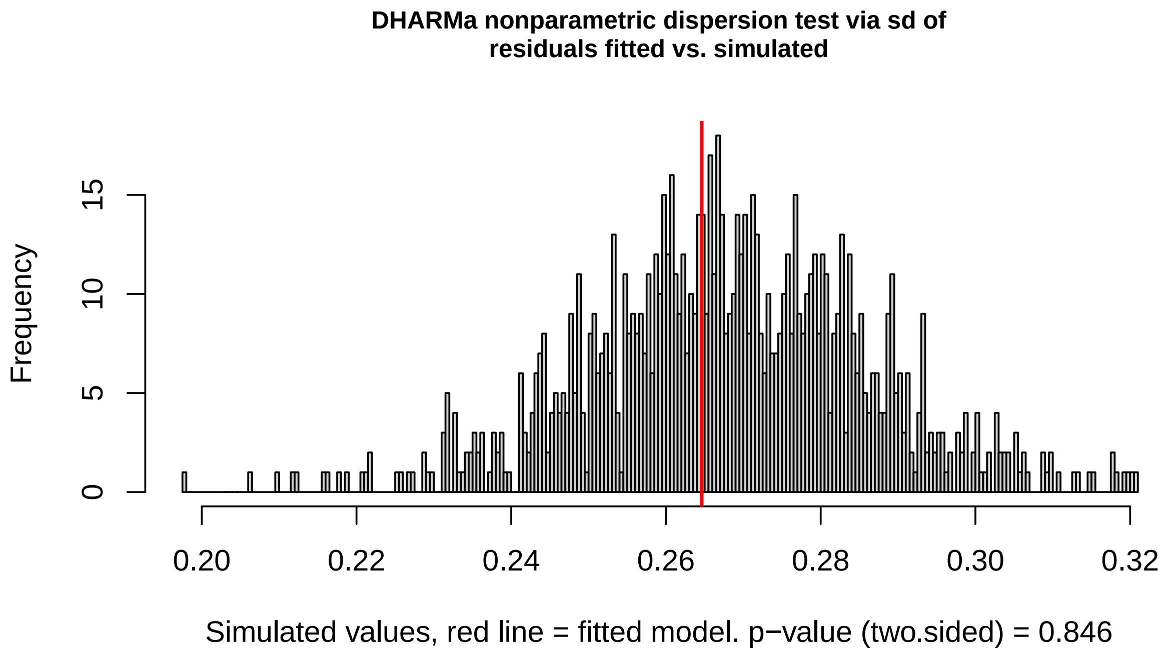

```
##
## DHARMa nonparametric dispersion test via sd of residuals fitted vs.
    simulated
##
## data: simulationOutput
## ratioObsSim = 0.98746, p-value = 0.846
## alternative hypothesis: two.sided
```

The pattern we observe in our own data is also found across sites by other groups, as demonstrated by our analysis of estimates extracted from the literature and given in Lit_behavior.csv, which contains: **river**: the river of origin of the fish
**course**: the course of the river of origin - upper or lower
**value**: the proportion of time the fish spent shoaling during a behavioral trial
**year**: the year in which the data were collected
**source**: the source of the data - either the paper from which they were extracted, or 'Lab', denoting that we collected them ourselves on fieldwork trip to Trinidad in March 2020.

```
db <- read.csv("Lit_behavior.csv")

# Model
shoaling_model <- glmmTMB(value ~ course + (1 | river) + (1 |
    year), db, family = beta_family(link = "logit"))
# summary(shoaling_model)
Anova(shoaling_model, type = "II", test.statistic = "Chisq")

## Analysis of Deviance Table (Type II Wald chisquare tests)
##
## Response: value
##          Chisq Df Pr(>Chisq)
```

```
## course 11.17  1  0.0008314 ***
## ---
## Signif. codes:  0 '***' 0.001 '**' 0.01 '*' 0.05 '.' 0.1 ' ' 1

# Effect size
parameters::model_parameters(shoaling_model)

## # Fixed Effects
##
## Parameter       | Coefficient |   SE |         95% CI |     z |      p
## ----------------------------------------------------------------------
## (Intercept)     |        0.39 | 0.21 | [-0.01,  0.79] |  1.89 | 0.058
## course [upper]  |       -1.18 | 0.35 | [-1.87, -0.49] | -3.34 | < .001
##
## # Random Effects
##
## Parameter                  | Coefficient
## ----------------------------------------
## SD (Intercept: river)      |    3.79e-06
## SD (Intercept: year)       |    6.96e-06
## SD (Residual)              |        2.43

z_to_r(3.34, dim(db)[1])

## r     |        95% CI
## ----------------------
## 0.58  | [0.28, 0.75]

# Diagnostic plots
sim_residuals_shoaling <- simulateResiduals(shoaling_model, 1000)
plot(sim_residuals_shoaling)
```

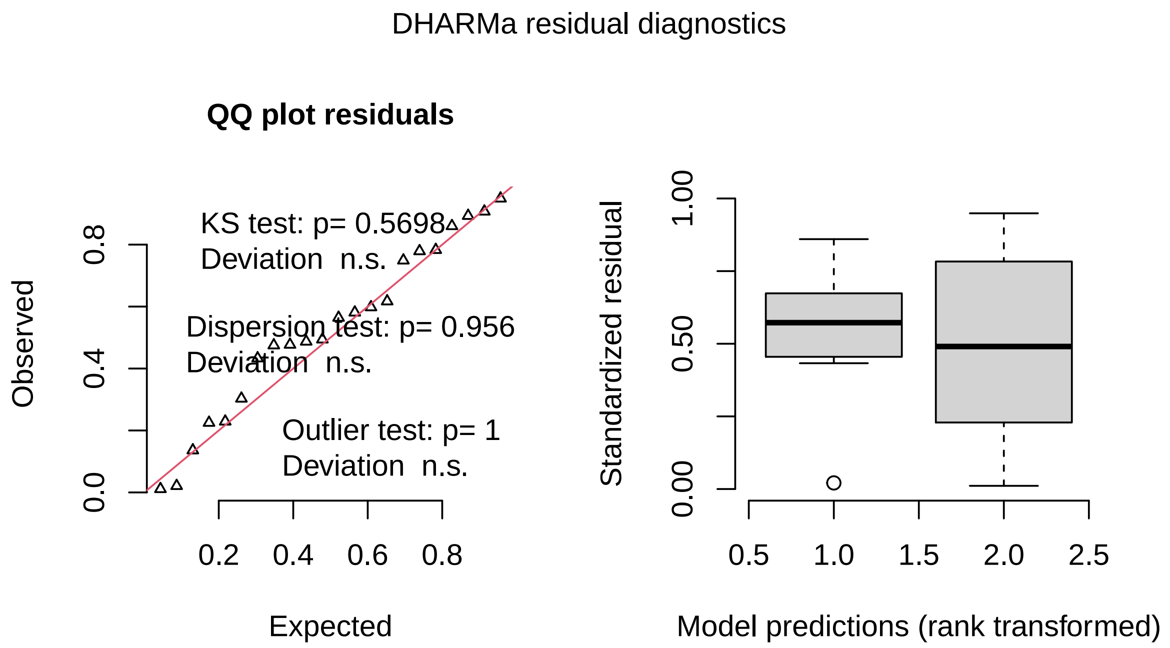

```
testDispersion(sim_residuals_shoaling)
```

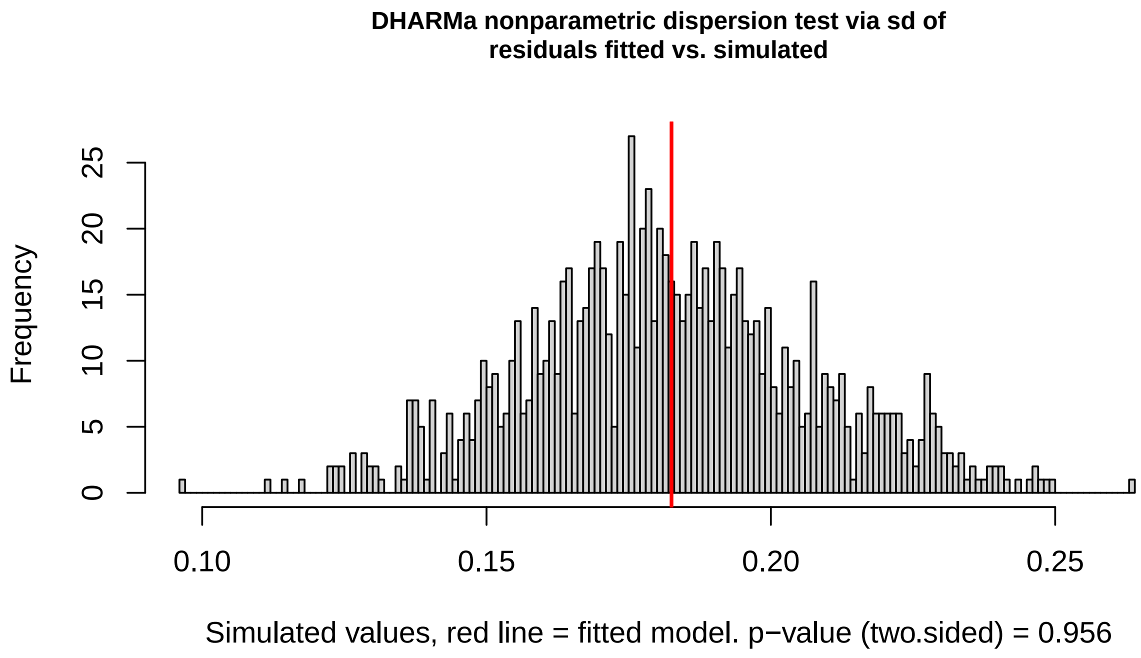

```
##
## DHARMa nonparametric dispersion test via sd of residuals fitted vs.
   simulated
##
## data:  simulationOutput
## ratioObsSim = 1.0006, p-value = 0.956
## alternative hypothesis: two.sided
```

#### 2.2 *Gyrodactylus* spp. prevalence differs across river courses - Fig. 5G

Here we provide the code used to test for difference in prevalence between guppy populations in the upper and lower courses of 11 Trinidadian rivers. These data come both from published estimates, and our own lab measurements, and are provided in the file 'Lit_prevalence.csv', which contains the following variables:

**river**: the river of origin of the fish
**course**: the course of the river of origin - upper or lower
**prev.yr**: the year in which the data were collected
**prev.n**: the number of fish sampled for that prevalence estimate
**prev**: the proportion of hosts that were found infected with *Gyrodactylus* spp.
**source**: the source of the data - either the paper from which they were extracted, or 'Lab', denoting that we collected them ourselves on fieldwork trip to Trinidad in March 2020.

```
dp <- read.csv("Lit_prevalence.csv")

# Model
prev_model <- glmmTMB(prev ~ course + (1 | river) + (1 | prev.yr),
    data = dp, family = beta_family(link = "logit"))
```

```
# Effect size parameters::model_parameters(prev_model)
# z_to_r(2.24,dim(dp)[1])

# summary(Mixed_model)
Anova(prev_model, type = "II", test.statistic = "Chisq")

## Analysis of Deviance Table (Type II Wald chisquare tests)
##
## Response: prev
##         Chisq Df Pr(>Chisq)
## course 5.0196  1    0.02506 *
## ---
## Signif. codes:  0 '***' 0.001 '**' 0.01 '*' 0.05 '.' 0.1 ' ' 1

# Diagnostic plots
sim_residuals_prev <- simulateResiduals(prev_model, 1000)
plot(sim_residuals_prev)
```

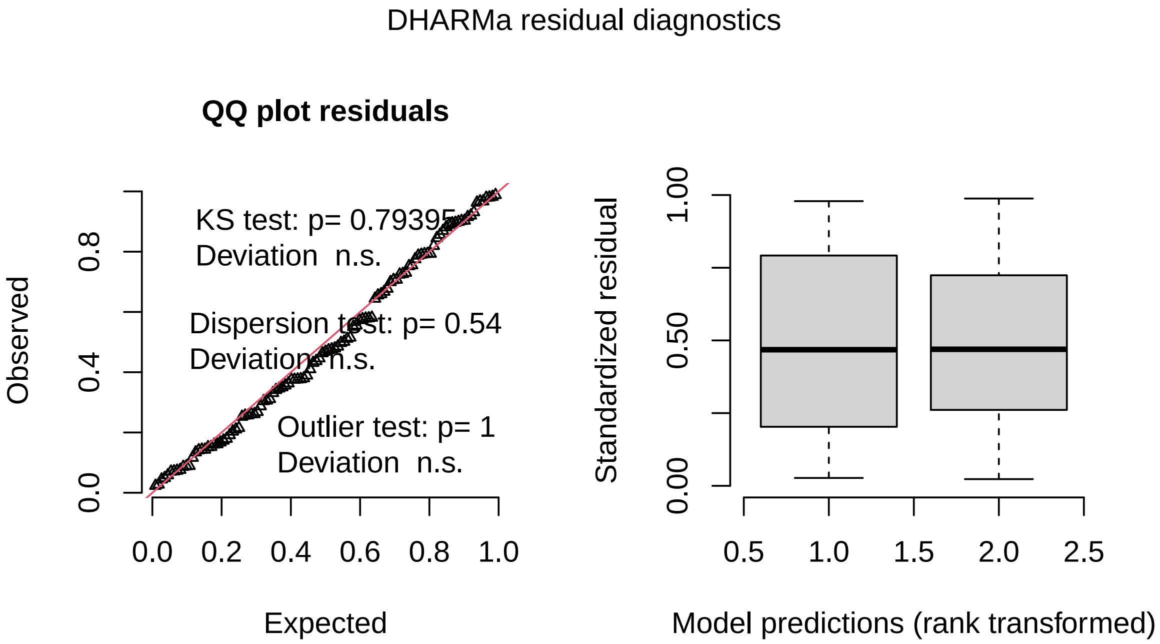

```
testDispersion(sim_residuals_prev)
```

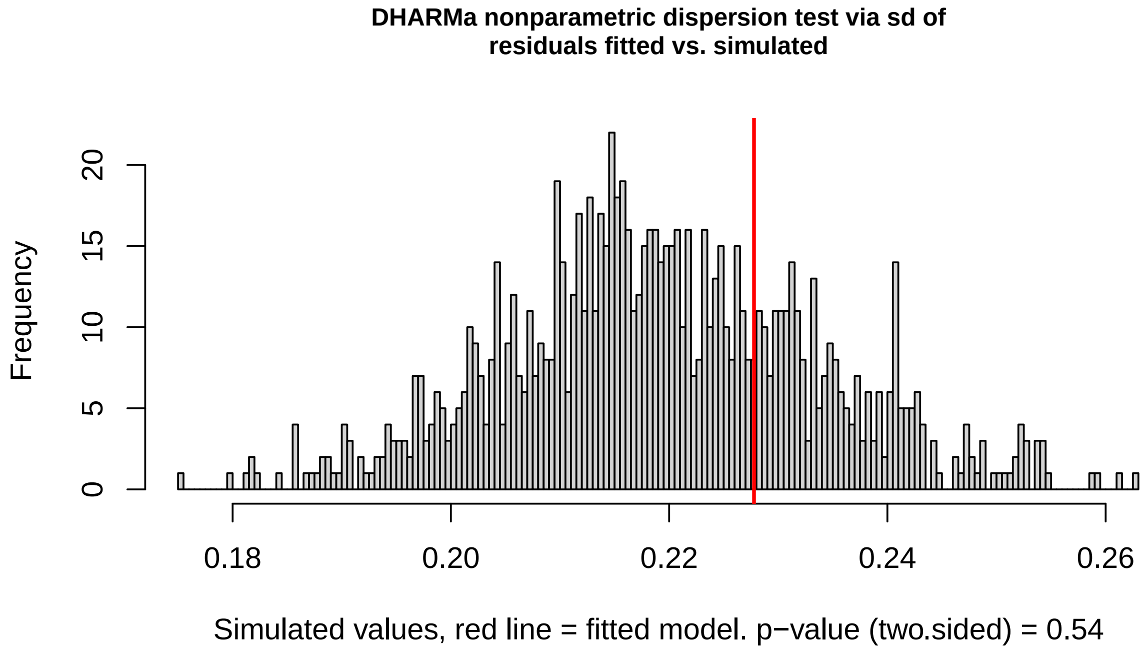

```
##
## DHARMa nonparametric dispersion test via sd of residuals fitted vs.
   simulated
##
## data:  simulationOutput
## ratioObsSim = 1.0393, p-value = 0.54
## alternative hypothesis: two.sided
```

##### 2.3 Guppy density differs non-significantly between courses - Fig. 5H

Here we provide the code used to test for difference in /% time spent shoaling between guppies from the upper and lower courses of five Trinidadian rivers. These data come both from published estimates, and our own lab measurements, and are provided in the file 'Lit_density.csv', which contains the following variables:

**river**: the river of origin of the fish
**course**: the course of the river of origin - upper or lower
**year.collected**: the year in which the data were collected
**value**: the estimated number of fish per square meter
**paper.source**: the source of the data

```
dd <- read.csv("Lit_density.csv")

# Model
density_model <- glmmTMB(value ~ course + (1 | river) + (1 |
    year.collected), data = dd)

# Effect size
parameters::model_parameters(density_model)
```

```
## # Fixed Effects
##
## Parameter       | Coefficient |   SE |          95% CI |    z |     p
## --------------------------------------------------------------------
## (Intercept)     |        4.04 | 3.01 | [-1.85, 9.93] | 1.34 | 0.179
## course [upper] |        4.18 | 2.83 | [-1.37, 9.73] | 1.48 | 0.140
##
## # Random Effects
##
## Parameter                          | Coefficient
## ------------------------------------------------
## SD (Intercept: river)              |        2.21
## SD (Intercept: year.collected) |        2.04
## SD (Residual)                      |        2.52
```

```
z_to_r(1.48, dim(dd)[1])
```

```
## r     |          95% CI
## ----------------------
## 0.29 | [-0.10, 0.58]
```

```
# summary(density_model)
Anova(density_model, type = "II")
```

```
## Analysis of Deviance Table (Type II Wald chisquare tests)
##
## Response: value
##          Chisq Df Pr(>Chisq)
## course 2.1757  1     0.1402
```

```
# Diagnostic plots
sim_residuals_density <- simulateResiduals(density_model, 1000)
plot(sim_residuals_density)
```

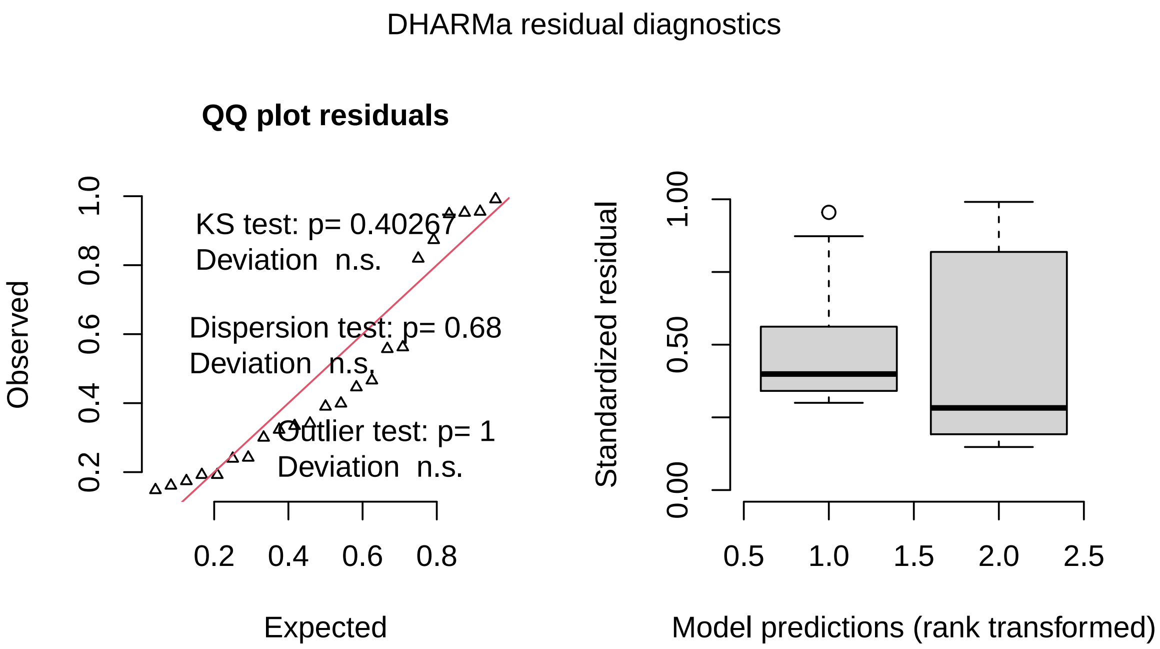

```
testDispersion(sim_residuals_density)
```

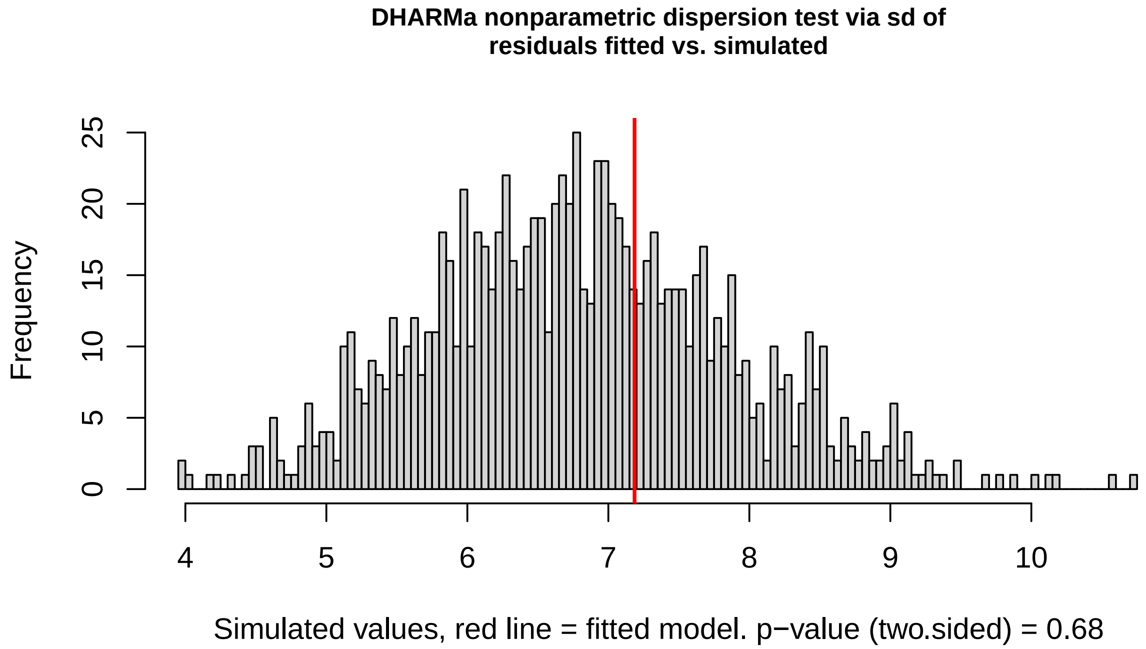

```
##
##  DHARMa nonparametric dispersion test via sd of residuals fitted vs.
    simulated
##
## data:  simulationOutput
## ratioObsSim = 1.0573, p-value = 0.68
```

```
## alternative hypothesis: two.sided
```

### 3 Testing model predictions of parasite intensity and virulence

#### 3.1 Testing for parasite intensity differences among lines from our four focal populations - Fig. 5I

This analysis uses the data in 'SI_parasitetraits.csv', which gives data per individual fish per row, and includes the following variables:

**line**: the identity of the isogenic line
**species**: the species identity of the line
**date**: the calendar date on which the observation was made in YYYYMMDD
**linf**: the total number of days the worm line was maintained in the lab
**lineday**: the number of days since the line was established in the lab
**lnldy**: lineday nested in line: gives each line/lineday combination a unique label to control for psuedoreplication in our analysis
**gyro**: the number of gyrodactylus found on the fish at that date
**type**: categorical variable describing whether the fish was dead, uninfected or infected
**course**: whether the isogenic line was initiated from a worm wild-caught in the upper or lower courses of a Trinidadian river, or from a commercially available lab stock fish ('Lab')
**river**: the river (or lab) from which the worm line originated
**lagint**: the mean intensity (total number of parasites/total number of infected fish) at the previous observation
**daysls**: the number of days since the previous observation
**lagdeathinf**: the number of dead fish/the number of infected fish at the previous observation [note this can be larger than 1 because new uninfected fish added at the previous timepoint would not be included in the denominator here, but may become infected and die by the subsequent timepoint]
**nfish**: the number of fish in the tank at the observation (excluding newly added uninfected fish)

##### 3.1.1 Data preprocessing

```
# Read data
df <- read.csv("SI_parasitetraits.csv")

# First, subset the data so there is 1 row per lnldy - i.e.
# per observation event, not per fish
drops <- c("gyro", "fish", "type", "gyroD")
df2 <- df[, !(names(df) %in% drops)]

ldd <- df2[!duplicated(df2, by = c("lnldy")), ]

# Now create timelagged variables, grouped by line
ldd <- ldd[order(ldd$line, ldd$lineday), ]
ldd <- ddply(ldd, .(line), .fun = mutate, laglineday = lag(lineday,
    n = 1, default = NA))

ldd <- ddply(ldd, .(line), .fun = mutate, inflastscreen = lag(ninftoday,
    n = 1, default = NA))

ldd <- ddply(ldd, .(line), .fun = mutate, Nlastscreen = lag(nfishn,
```

```
    n = 1, default = NA))

ldd <- ddply(ldd, .(line), .fun = mutate, Nglastscreen = lag(totgD,
    n = 1, default = NA))

ldd$daysls <- ldd$lineday - ldd$laglineday

ldd$lagint <- ldd$Nglastscreen/ldd$inflastscreen

# Define death proportion as the proportion of of infected
# hosts who died
ldd$lagdeathinf <- (ldd$ndeadtoday/ldd$inflastscreen)

# And death rate as death proportion divided by the days over
# which it occurred
ldd$lagdeathinfr <- ldd$lagdeathinf/ldd$daysls

# Update here % of checks within 1 day of 3
length(which(ldd$daysls >= 2 & ldd$daysls <= 4))/781

## [1] 0.7912932

# checking these variables - lagdeathinf
summary(ldd$lagdeathinf)

##     Min. 1st Qu.  Median    Mean 3rd Qu.    Max.    NA's
##  0.0000  0.0000  0.0000  0.1506  0.2000  2.0000      61

# 47 NAs are the starting days of each line, 19 extra NAs
# here reflect days where there were no infected fish
ldd$lagdeathinf[is.nan(ldd$lagdeathinf)] <- 0  # no deaths = 0

# Remove NAs
lddtv <- ldd[which(is.finite(ldd$lagdeathinf)), ]
lddtv <- lddtv[which(is.finite(lddtv$lagint)), ]

# Only consider up to day 65 for each line. Lines can begin
# to perform poorly after longer maintenance in the lab than
# this. We subset the data to day 65 because at this point
# many lines crashed, likely due to COVID-related changes in
# our management practices.
thresh_day <- 65
lddtv2 <- subset(lddtv, lineday < thresh_day)
lddtv_tradeoff <- subset(lddtv2, linf > 30)

# Remove 0s from the fish level dataframe
dfl <- df[which(df$gyro > 0), ]
# subset to focus on aripo and guanapo up to day
# thresh_day=65
thresh_day <- 65

dfagl <- subset(dfl, river == "Aripo" | river == "Guanapo")
dfl2 <- subset(dfl, lineday < thresh_day)
dfagl65 <- subset(dfagl, lineday < thresh_day)
```

```r
# subset the observation-level dataset to just the focal
# rivers
lddagALL <- subset(lddtv, river == "Aripo" | river == "Guanapo")
lddag <- subset(lddtv2, river == "Aripo" | river == "Guanapo")

# Can see that lines began performing poorly after day 65,
# likely due to COVID-induced changes in lab.
plot(lddagALL$totgD ~ lddagALL$lineday)
abline(v = thresh_day, col = "red")
```

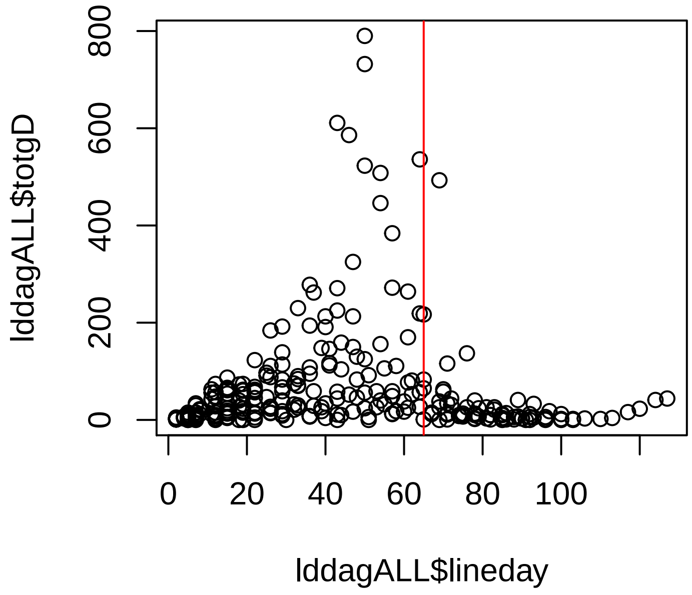

```r
# Test of all lines differing in intensity
dfl2 <- subset(dfl, lineday < thresh_day)
dfl3 <- subset(dfl2, linf > 30)
dfl4 <- subset(dfl3, gyro > 0)
ii <- Anova(lm(log(gyro) ~ line, data = dfl4), type = "II")
eta_squared(ii)

## # Effect Size for ANOVA
##
## Parameter | Eta2 |       90% CI
## ---------------------------------
## line      | 0.13 | [0.09, 0.15]

# Can make diagnostic plots plot(ii)
```

###### 3.1.2 Checking for correlations among variables

```r
# This function is from 'Mixed effects models and extensions
# in ecology with R' (2009).Zuur, AF, Ieno, EN, Walker, N,
# Saveliev, AA, and Smith, GM. Springer.
panel.cor <- function(x, y, digits = 2, prefix = "", cex.cor,
    ...) {
    usr <- par("usr")
    on.exit(par(usr))
    par(usr = c(0, 1, 0, 1))
    r <- abs(cor(x, y))
    txt <- format(c(r, 0.123456789), digits = digits)[1]
    txt <- paste(prefix, txt, sep = "")
    if (missing(cex.cor))
        cex.cor <- 0.8/strwidth(txt)
    text(0.5, 0.5, txt, cex = 1)
}

pairs(~nfish + lineday, data = dfagl65, lower.panel = panel.smooth,
    upper.panel = panel.cor, na.action = na.omit)
```

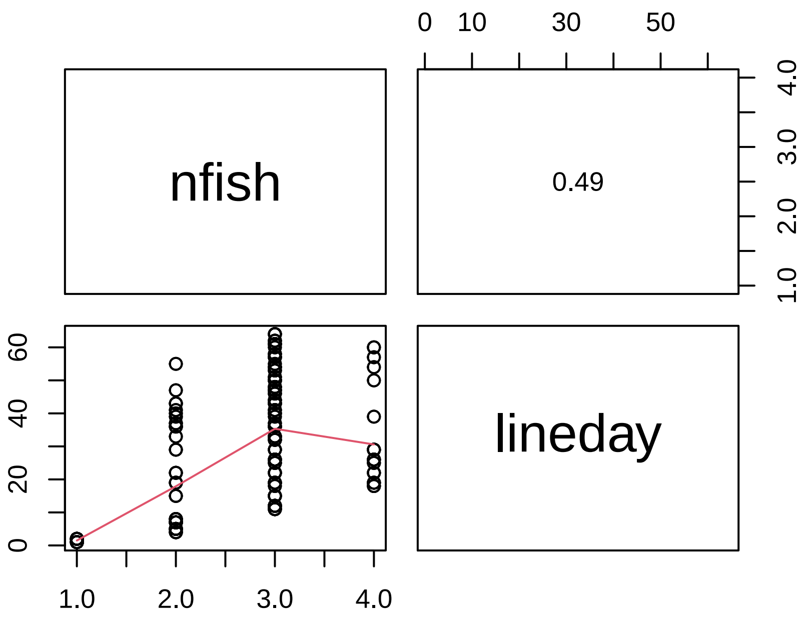

##### 3.1.3 GLMM for parasite intensity

This is the result presented in Fig. 5I.

```r
# To analyze the effect of course on intensity of all lines
# in the focal rivers
m_int <- glmmTMB(log(gyro) ~ course + river + nfish + (1 | line) +
    (1 | lnldy), data = dfagl65, na.action = na.omit)

# Effect size parameters::model_parameters(m_int)
```

```
# z_to_r(2.20,dim(dfagl65)[1])

# TAKE PARTIALS HERE AND AT DEATH summary(m_int)
Anova(m_int, type = 2)

## Analysis of Deviance Table (Type II Wald chisquare tests)
##
## Response: log(gyro)
##          Chisq Df Pr(>Chisq)
## course  4.8453  1    0.02772 *
## river   4.7907  1    0.02861 *
## nfish  56.1172  1  6.828e-14 ***
## ---
## Signif. codes:  0 '***' 0.001 '**' 0.01 '*' 0.05 '.' 0.1 ' ' 1

# Diagnostic plots
sim_residuals_int <- simulateResiduals(m_int, 1000)
plot(sim_residuals_int)
```

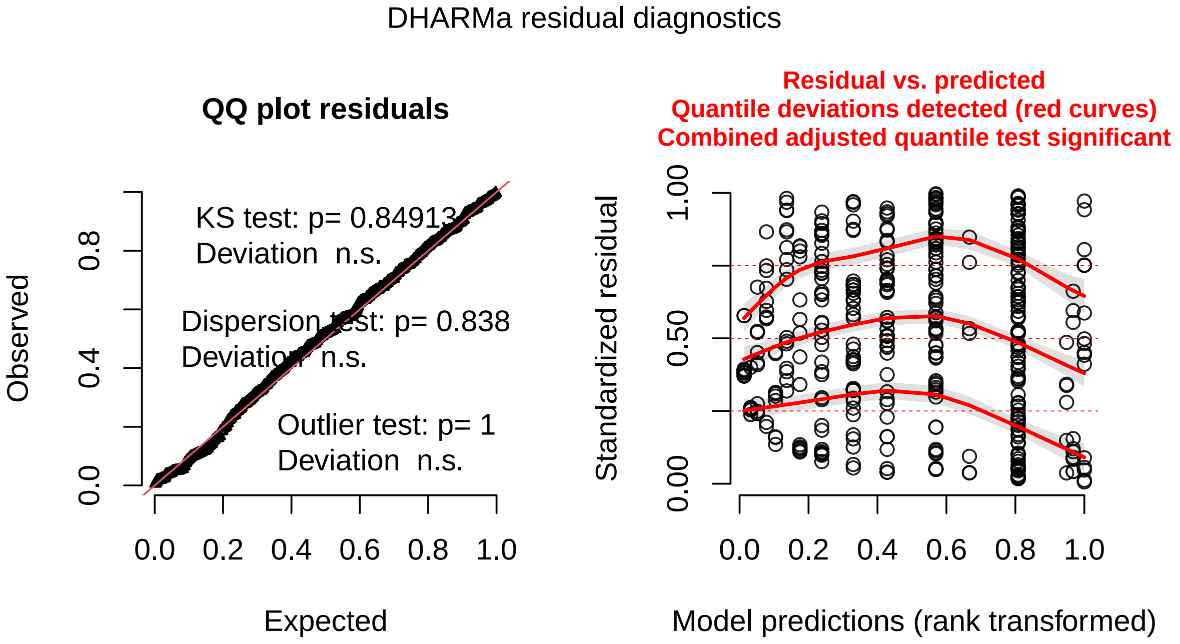

```
testDispersion(sim_residuals_int)
```

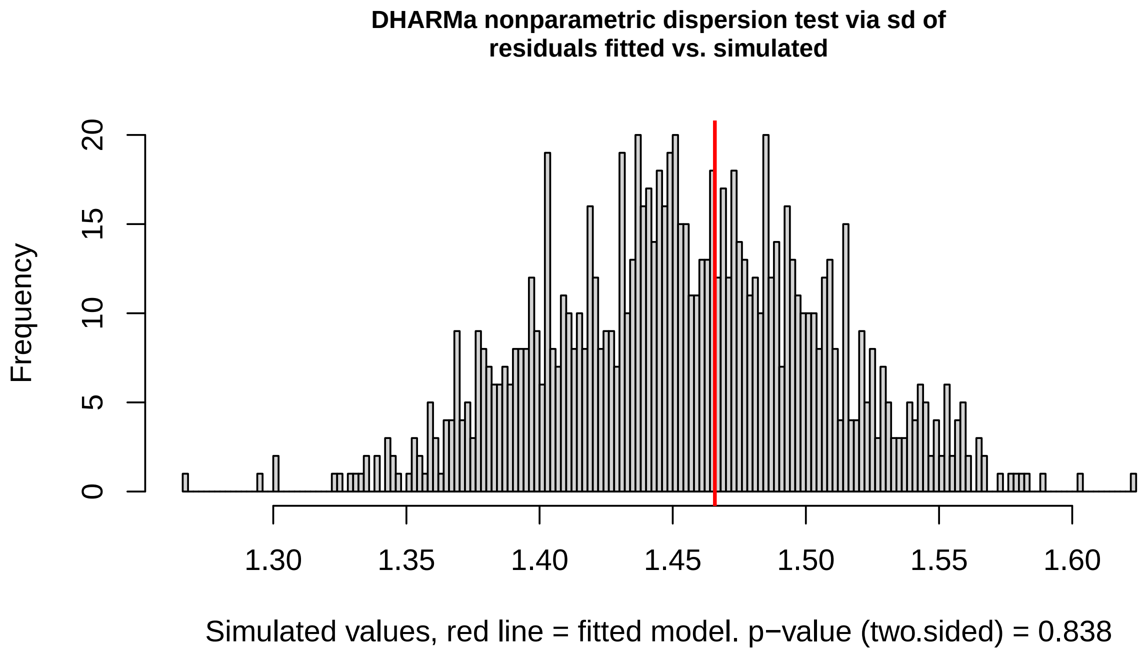

```
##
## DHARMa nonparametric dispersion test via sd of residuals fitted vs.
   simulated
##
## data:  simulationOutput
## ratioObsSim = 1.0078, p-value = 0.838
## alternative hypothesis: two.sided
```

###### 3.1.4 Using just the lines of known species identity, *G. turnbulli* and *G. bullatarudis* do not differ in their intensity

We identified some of our lab maintained lines to species level, and here test whether they differ in their intensities.

```
# subset to look only at lower courses as Gb lines only came
# from the lower course.

dfl_lo <- subset(dfl2, course == "Lower")

m_everyone <- glmmTMB(log(gyro) ~ species + river + nfish + (1 |
    line) + (1 | lineday), data = dfl_lo, na.action = na.omit)
# summary(m_everyone)
Anova(m_everyone, type = 2)

## Analysis of Deviance Table (Type II Wald chisquare tests)
##
## Response: log(gyro)
##            Chisq Df Pr(>Chisq)
## species   2.7295  1    0.09851 .
## river     6.6059  2    0.03677 *
## nfish    18.3203  1  1.867e-05 ***
```

```
## ---
## Signif. codes:  0 '***' 0.001 '**' 0.01 '*' 0.05 '.' 0.1 ' ' 1

# Effect size parameters::model_parameters(m_everyone)
# z_to_r(1.65,dim(dfl_lo)[1])

# Diagnostic plots
sim_residuals_everyone <- simulateResiduals(m_everyone, 1000)
plot(sim_residuals_everyone)
```

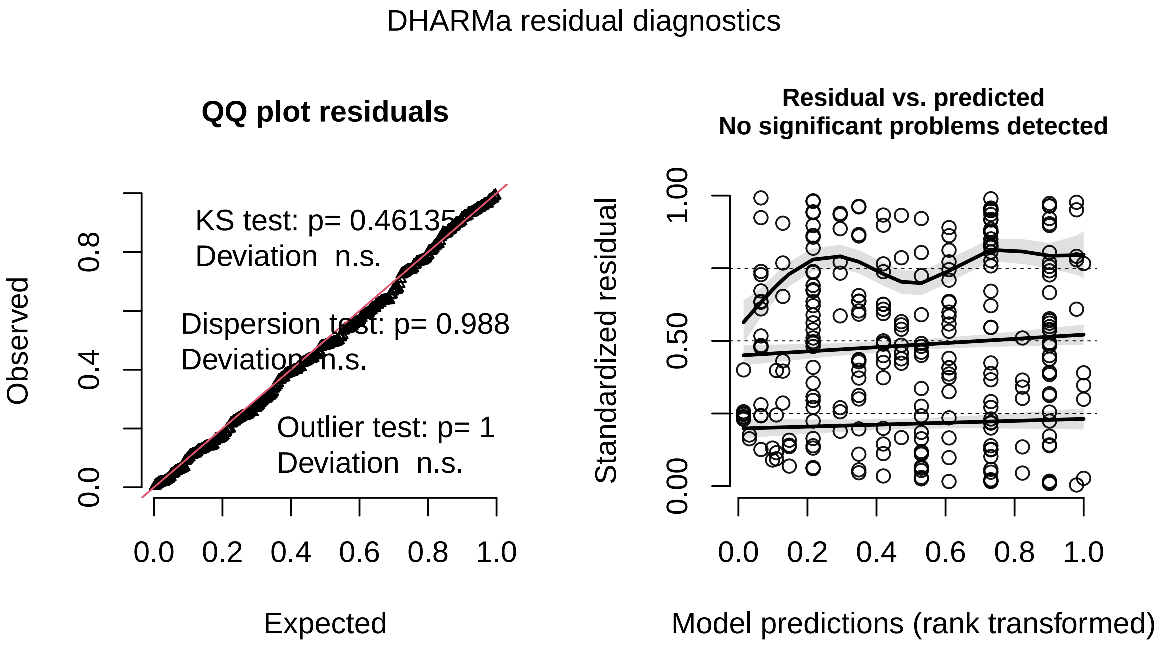

```
testDispersion(sim_residuals_everyone)
```

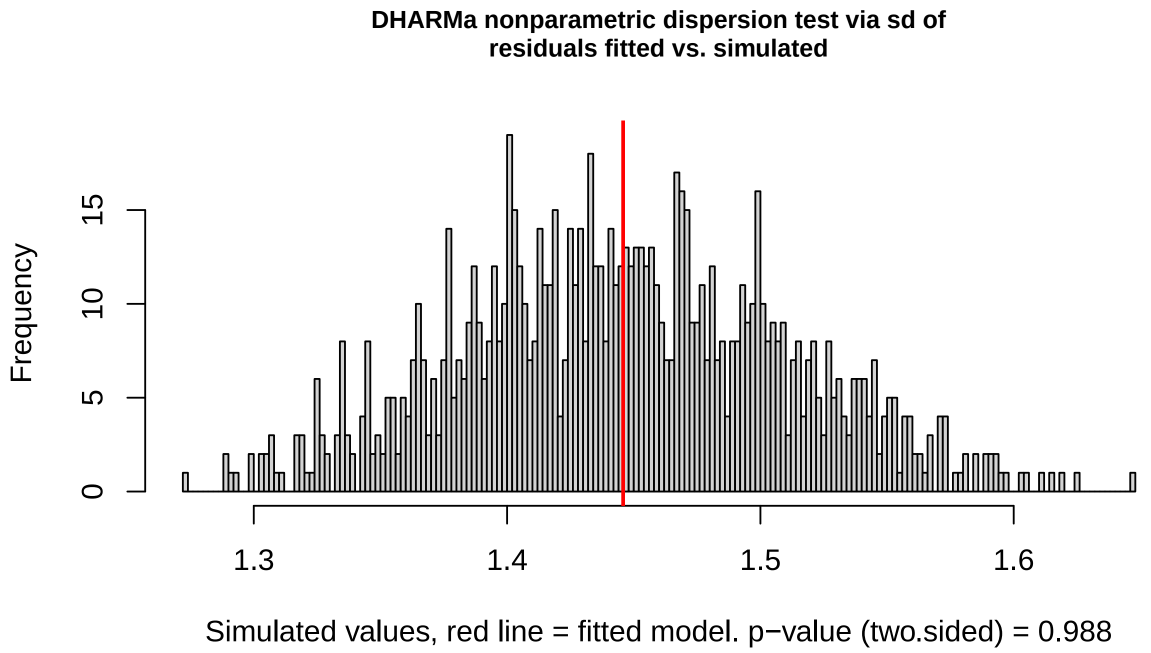

```
##
## DHARMa nonparametric dispersion test via sd of residuals fitted vs.
   simulated
##
## data:  simulationOutput
## ratioObsSim = 0.99985, p-value = 0.988
## alternative hypothesis: two.sided
```

###### 3.1.5 Lower course *Gyrodactylus turnbulli* tend to attain higher intensities than their upper course counterparts

For *G. turnbulli* only (because we only had 3 confirmed *G. bullatarudis* lines, all from the lower course), we test whether worms from the upper or lower course attain higher intensities.

```
dfagl65_Gt <- subset(dfagl65, species == "t")

# Model for Gt only
m_Gt <- glmmTMB(log(gyro) ~ course + river + nfish + (1 | line) +
    (1 | lnldy), data = dfagl65_Gt, na.action = na.omit)
# summary(m_Gt)
Anova(m_Gt, type = 2)

## Analysis of Deviance Table (Type II Wald chisquare tests)
##
## Response: log(gyro)
##           Chisq Df Pr(>Chisq)
## course   3.1474  1    0.07605 .
## river    4.5287  1    0.03333 *
## nfish   24.1918  1   8.72e-07 ***
## ---
## Signif. codes:  0 '***' 0.001 '**' 0.01 '*' 0.05 '.' 0.1 ' ' 1
```

```
# Effect size parameters::model_parameters(m_Gt)
# z_to_r(1.77,dim(dfagl65_Gt)[1])

sim_residuals_Gt_int <- simulateResiduals(m_Gt, 1000)
plot(sim_residuals_Gt_int)
```

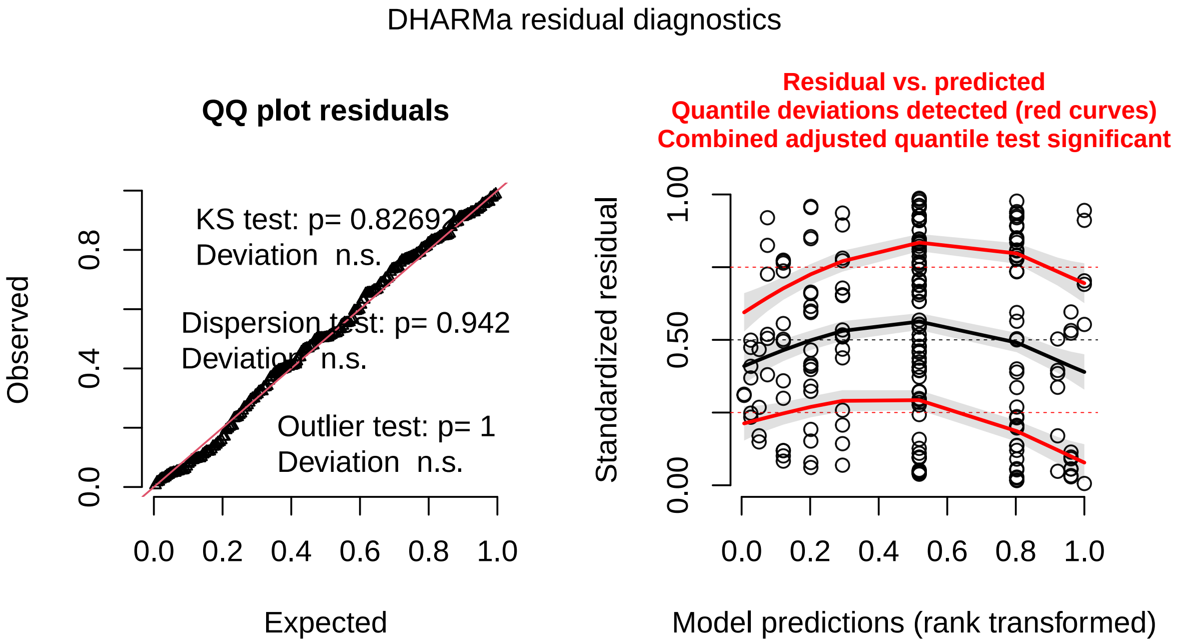

```
testDispersion(sim_residuals_Gt_int)
```

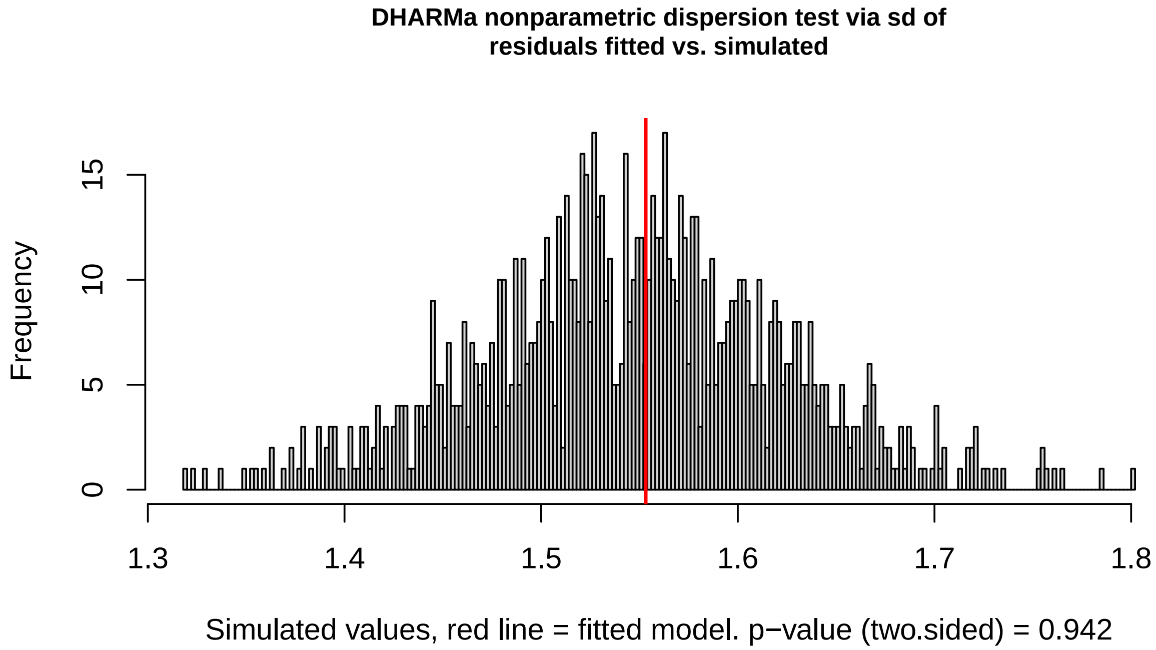

```
##
## DHARMa nonparametric dispersion test via sd of residuals fitted vs.
   simulated
##
## data:  simulationOutput
## ratioObsSim = 1.004, p-value = 0.942
## alternative hypothesis: two.sided
```

#### 3.2 Testing for host death rate differences among lines from our four focal populations - Fig. 5J

##### 3.2.1 Checking for correlations among variables

```
pairs(~nfish + lineday + linf + lagint, data = lddag, lower.panel = panel.
   smooth,
    upper.panel = panel.cor, na.action = na.omit)
```

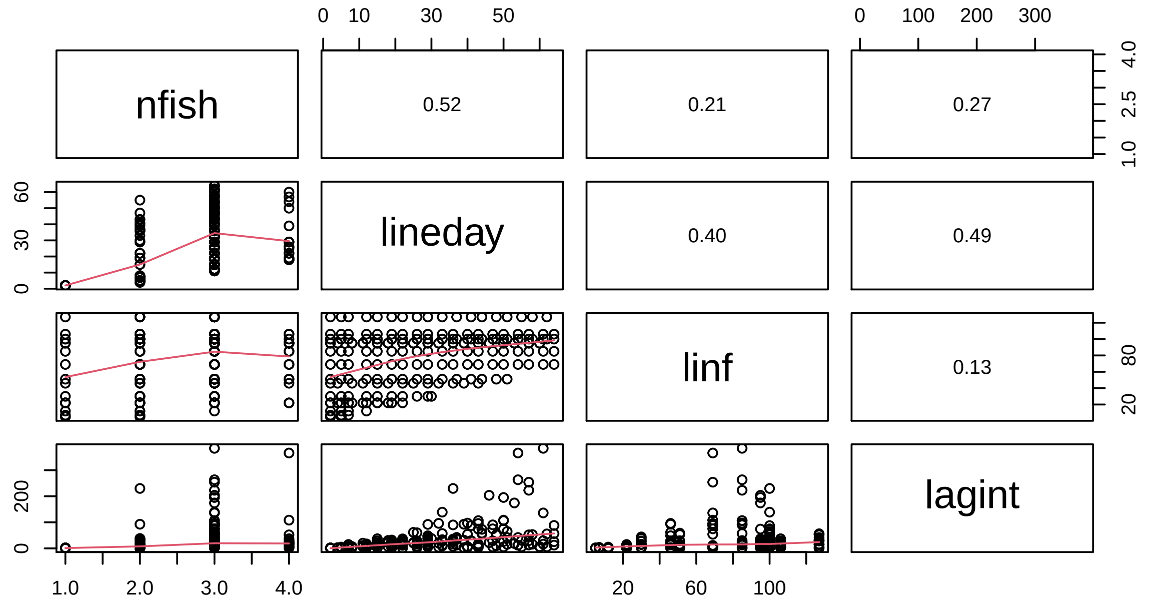

##### 3.2.2 GLMM for host death rate

```
# Test infected host death rate across courses Main result
# presented in Figure 5J
mdeath <- glmmTMB(lagdeathinfr ~ course + river + (1 | line) +
     (1 | lnldy), family = tweedie(link = "log"), data = lddag)

# Effect size parameters::model_parameters(mdeath)
# z_to_r(2.18,dim(lddag)[1])

# summary(m4)
Anova(mdeath, type = 2)
```

```
## Analysis of Deviance Table (Type II Wald chisquare tests)
##
## Response: lagdeathinfr
##          Chisq Df Pr(>Chisq)
## course 4.7452  1    0.02938 *
## river  0.0013  1    0.97162
## ---
## Signif. codes:  0 '***' 0.001 '**' 0.01 '*' 0.05 '.' 0.1 ' ' 1
```

```
# Diagnostic plots
death_residuals <- simulateResiduals(mdeath, n = 1000)
plot(death_residuals)
```

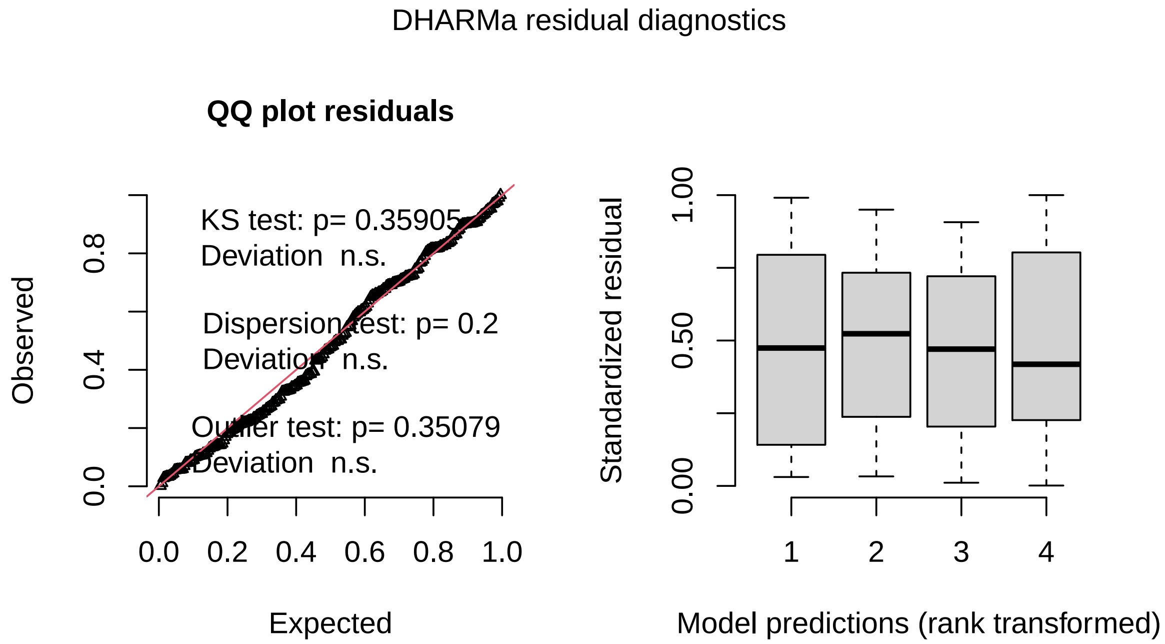

```
testDispersion(death_residuals)
```

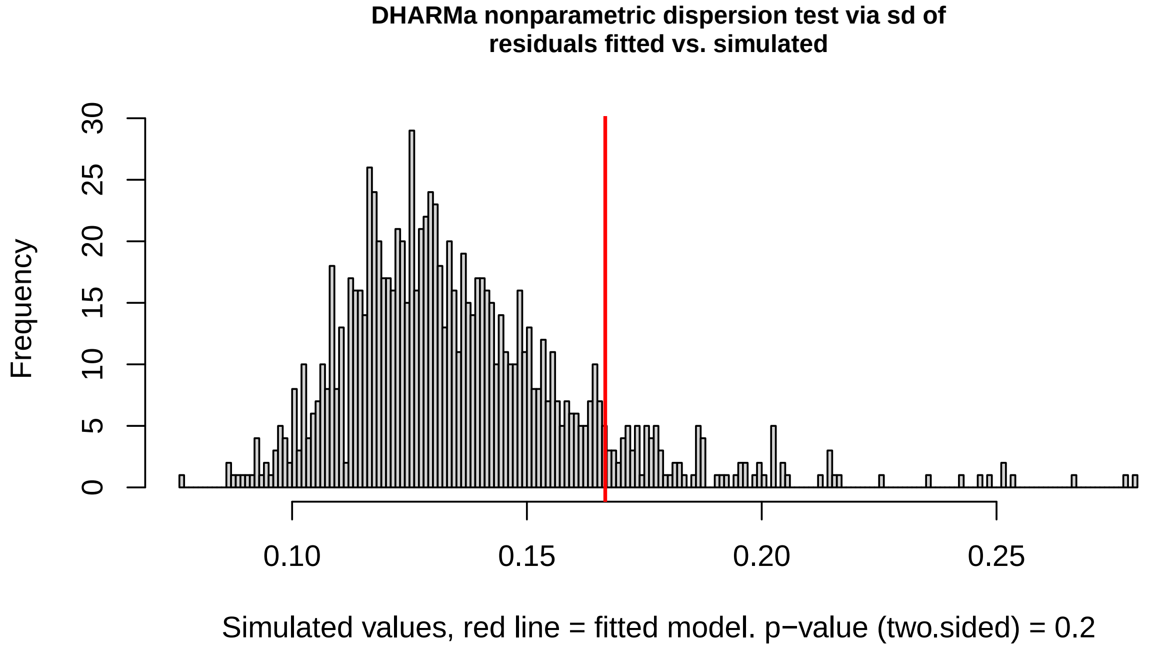

```
##
## DHARMa nonparametric dispersion test via sd of residuals fitted vs.
   simulated
##
## data:  simulationOutput
## ratioObsSim = 1.2264, p-value = 0.2
## alternative hypothesis: two.sided
```

###### 3.2.3 GLMMs for host death rate among lines of known species identity

For a subset of the lines, we were able to identify the worms to species level. Here we test whether the species differ in the rate at which they kill hosts, and,

###### 3.2.4 The species do not differ in the death rates of their hosts

```
# Use only lower course lines
lddag_lo <- subset(lddtv2, course == "Lower")

# Test effect of species on death rate
m_everyone <- glmmTMB(lagdeathinfr ~ species + river + (1 | line) +
    (1 | lnldy), family = tweedie(link = "log"), data = lddag_lo)

# Effect size parameters::model_parameters(m_everyone)
# z_to_r(0.94,dim(lddag_lo)[1])

# summary(m_everyone)
Anova(m_everyone, type = 2)

## Analysis of Deviance Table (Type II Wald chisquare tests)
```

```
##
## Response: lagdeathinfr
##           Chisq Df Pr(>Chisq)
## species 0.8907  1    0.34528
## river   5.4850  2    0.06441 .
## ---
## Signif. codes:  0 '***' 0.001 '**' 0.01 '*' 0.05 '.' 0.1 ' ' 1

# Diagnostic plots
sim_residuals_everyone_death <- simulateResiduals(m_everyone,
    1000)
plot(sim_residuals_everyone_death)
```

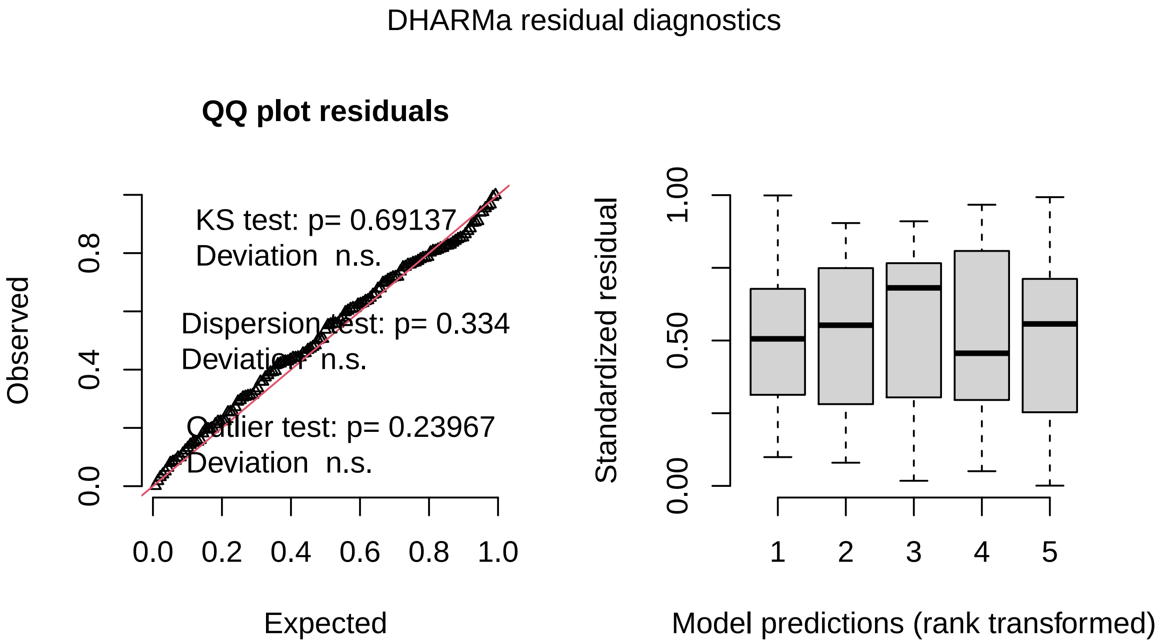

```
testDispersion(sim_residuals_everyone_death)
```

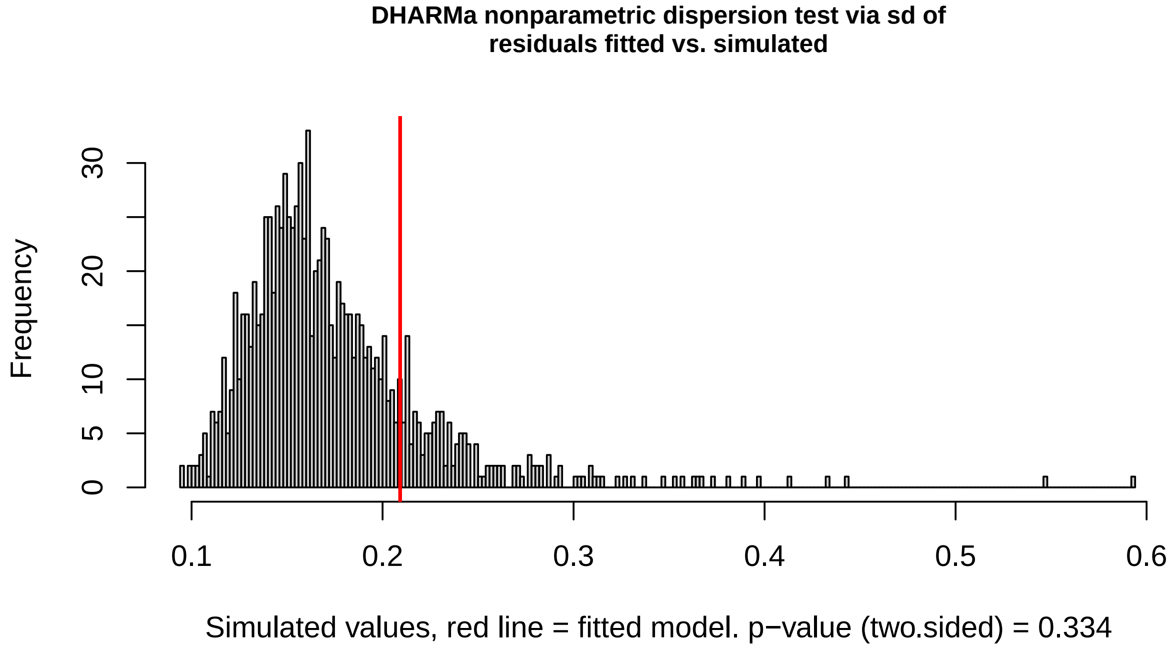

```
##
## DHARMa nonparametric dispersion test via sd of residuals fitted vs.
   simulated
##
## data: simulationOutput
## ratioObsSim = 1.2013, p-value = 0.334
## alternative hypothesis: two.sided
```

###### 3.2.5 Lower course *Gyrodactylus turnbulli* do cause higher host death rates than their upper course counterparts

For *G. turnbulli* only (because we only had 3 confirmed *G. bullatarudis* lines, all from the lower course, we test whether worms from the upper or lower course kill fish at a faster rate.

```
# Subset to death rate imposed by known Gts
lddag_Gt <- subset(lddag, species == "t")

m_Gt <- glmmTMB(lagdeathinfr ~ course + river + (1 | line) +
    (1 | lnldy), family = tweedie(link = "log"), data = lddag_Gt)

# Effect size parameters::model_parameters(m_Gt)
# z_to_r(2.62,dim(lddag_Gt)[1])

# summary(m_Gt)
Anova(m_Gt, type = 2)

## Analysis of Deviance Table (Type II Wald chisquare tests)
##
## Response: lagdeathinfr
##         Chisq Df Pr(>Chisq)
## course 6.862  1   0.008805 **
```

```
## river   2.785  1   0.095152 .
## ---
## Signif. codes:  0 '***' 0.001 '**' 0.01 '*' 0.05 '.' 0.1 ' ' 1

# Diagnostic plots
sim_residuals_Gt_death <- simulateResiduals(m_Gt, 1000)
plot(sim_residuals_Gt_death)
```

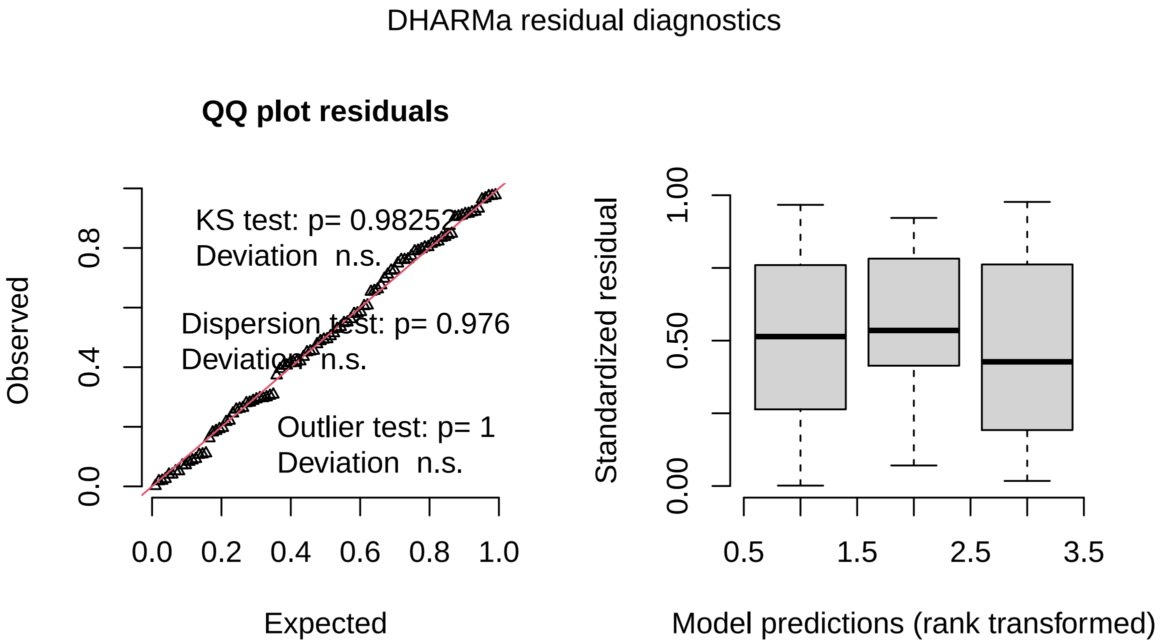

```
testDispersion(sim_residuals_Gt_death)
```

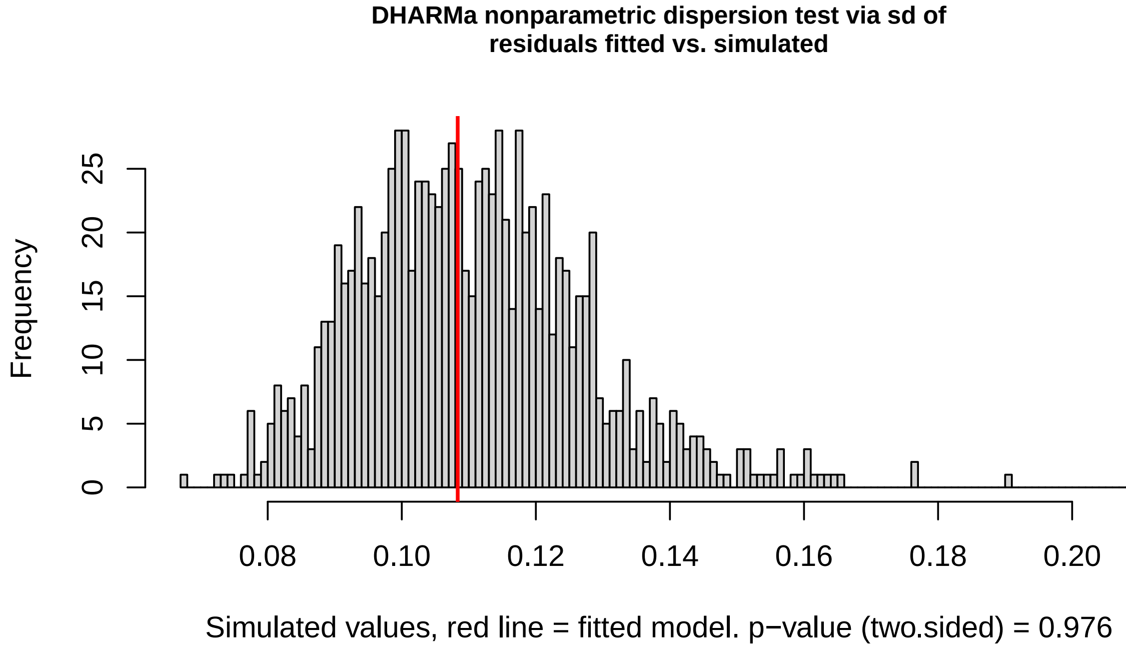

```
##
## DHARMa nonparametric dispersion test via sd of residuals fitted vs.
   simulated
##
## data:  simulationOutput
## ratioObsSim = 0.9799, p-value = 0.976
## alternative hypothesis: two.sided
```

### 4 Coevolutionary model - Fig. 4

This section contains a model and evolutionary analysis of that model for a logistically growing host and a parasite capable of superinfection (similar to coinfection) that evolves along a virulence-transmission tradeoff. Similarly, hosts can recover from infection. See the main text for further details.

#### 4.1 Model setup

```
# The equilibrium of S and a single parasite is feasible as
# long as (d+v+y)/B < (a-d)/q, derived from invasion of I
# into S alone

# Note that here d is all non-parasite induced mortality,
# which will be split later into background and predation B
# is transmission rate and = c*T

# Calculated equilibrium densities symbolically Susceptible
# host density
S_eq = function(a, q, d, B, v, y, w) {
    (d + v + y)/B
```

```
}

# Infected host density
I_eq = function(a, q, d, B, v, y, w) {
    -((d + w) * (2 * q * y^3 + 4 * d * q * y^2 + 2 * d^2 * q *
        y + 2 * q * v * y^2 + 2 * q * w * y^2 - B * y * ((B *
        d^4 + 4 * d^3 * q * v + B * a^2 * d^2 + B * a^2 * w^2 +
        B * a^2 * y^2 + B * d^2 * v^2 + B * d^2 * w^2 + B * d^2 *
        y^2 + B * v^2 * w^2 + 4 * d^2 * q * v^2 + 4 * q * v^2 *
        w^2 - 2 * B * a * d^3 + 2 * B * d^3 * v + 2 * B * d^3 *
        w + 2 * B * d^3 * y - 2 * B * a * d^2 * v - 2 * B * a *
        d * w^2 - 4 * B * a * d^2 * w + 2 * B * a^2 * d * w -
        2 * B * a * d * y^2 - 4 * B * a * d^2 * y + 2 * B * a^2 *
        d * y - 2 * B * a * v * w^2 + 2 * B * d * v * w^2 + 2 *
        B * d * v^2 * w + 2 * B * a^2 * w * y + 4 * B * d^2 *
        v * w + 2 * B * d^2 * v * y + 2 * B * d^2 * w * y + 4 *
        d * q * v * w^2 + 8 * d * q * v^2 * w + 8 * d^2 * q *
        v * w + 4 * d * q * v * y^2 + 4 * d * q * v^2 * y + 8 *
        d^2 * q * v * y + 4 * q * v * w * y^2 + 4 * q * v * w^2 *
        y + 4 * q * v^2 * w * y - 4 * B * a * d * v * w - 2 *
        B * a * d * v * y - 4 * B * a * d * w * y - 2 * B * a *
        v * w * y + 2 * B * d * v * w * y + 12 * d * q * v *
        w * y)/B)^(1/2) - B * a * y^2 + B * d * y^2 + B * d^2 *
        y + B * v * w * y + 2 * d * q * v * y + 2 * d * q * w *
        y + 2 * q * v * w * y - B * a * d * y - B * a * w * y +
        B * d * v * y + B * d * w * y))/(2 * y * (B * q * d^2 +
        2 * B * q * d * w + 2 * B * q * d * y + B * q * w^2 +
        2 * B * q * w * y + B * q * y^2))
}

# Recovered host density
R_eq = function(a, q, d, B, v, y, w) {
    y * I_eq(a, q, d, B, v, y, w)/(d + w)
}

# Total host density
H_eq = function(a, q, d, B, v, y, w) {
    S_eq(a, q, d, B, v, y, w) + I_eq(a, q, d, B, v, y, w) + R_eq(a,
        q, d, B, v, y, w)
}

# Proportion of hosts that are infected
prev_eq = function(a, q, d, B, v, y, w) {
    I_eq(a, q, d, B, v, y, w)/(S_eq(a, q, d, B, v, y, w) + I_eq(a,
        q, d, B, v, y, w) + R_eq(a, q, d, B, v, y, w))
}

# Use the simplest possible function for host shoaling rate
# and transmissibility combine to determine transmission
# rate. B=c*T
Transmission_func = function(c, Ti) {
    c * Ti
}
```

```
# Define a function to capture all, non-virulent host
# mortality What was previously d is now split into d+pred/c.
# The c1 constant is ommitted since it is chosen as 1 for
# simplicity
host_death = function(back_d, pred, c) {
    back_d + pred/c
}

# Define virulence as a function of transmission rate
virulence_func = function(parm1, parm2, Ti) {
    parm1 * Ti^parm2
}

# Transmission between two host genotypes differing in
# shoaling rate is the geometric mean, as mass action would
# suggest
trans_func = function(c1, c2, Ti) {
    sqrt(c1 * c2) * Ti
}
```

#### 4.2 Calculating evolution

```
# Define virulence as a function of transmission rate
virulence_func = function(parm1, parm2, Ti) {
    parm1 * Ti^parm2
}

# Calculate transmissibility, T, given virulence, v, and
# tradeoff parameters parm1 and parm2
inv_virulence_func = function(parm1, parm2, v) {
    (v/parm1)^(1/parm2)
}

# Define the superinfection function. Here is where one could
# rewrite to implement a superinfection rate dependent on
# Transmissibility, if desirable
superinfection_func = function(Ti, Tj, sigma) {
    sigma
}

# Define components of parasite fitness. I broke this up into
# three components for convenience. Note that from here on
# out we use d just to refer to the non-predation part of
# host background death rate.
inv_fitA = function(a, q, d, pred, c, parm1, parm2, Tm, Tr, y,
    w, sigma) {
    Transmission_func(c, Tm) * S_eq(a, q, host_death(d, pred,
        c), Transmission_func(c, Tr), virulence_func(parm1, parm2,
        Tr), y, w)
}

# Death and recovery of hosts infected with the invader
inv_fitB = function(a, q, d, pred, c, parm1, parm2, Tm, Tr, y,
```

```
    w, sigma) {
    -host_death(d, pred, c) - virulence_func(parm1, parm2, Tm) -
        y
}

# Gains and losses from superinfection
inv_fitC = function(a, q, d, pred, c, parm1, parm2, Tm, Tr, y,
    w, sigma) {
    I_eq(a, q, host_death(d, pred, c), Transmission_func(c, Tr),
        virulence_func(parm1, parm2, Tr), y, w) * (superinfection_func(Tm,
        Tr, sigma) * Transmission_func(c, Tm) - superinfection_func(Tr,
        Tm, sigma) * Transmission_func(c, Tr))
}

# Put the three components together into one function
inv_fit = function(a, q, d, pred, c, parm1, parm2, Tm, Tr, y,
    w, sigma) {
    inv_fitA(a, q, d, pred, c, parm1, parm2, Tm, Tr, y, w, sigma) +
        inv_fitB(a, q, d, pred, c, parm1, parm2, Tm, Tr, y, w,
            sigma) + inv_fitC(a, q, d, pred, c, parm1, parm2,
        Tm, Tr, y, w, sigma)
}

# Take derivatives of inv_fit
Fit_diffTm = Deriv(inv_fit, "Tm")
Fit_diffTm2 = Deriv(Fit_diffTm, "Tm")
Fit_diffTr = Deriv(inv_fit, "Tr")
Fit_diffTr2 = Deriv(Fit_diffTr, "Tr")

# Evaluate derivatives when Tm=Tr
Diff1Tm = function(a, q, d, pred, c, parm1, parm2, Tm, y, w,
    sigma) {
    Fit_diffTm(a, q, d, pred, c, parm1, parm2, Tm, Tm, y, w,
        sigma)
}

Diff2Tm = function(a, q, d, pred, c, parm1, parm2, Tm, y, w,
    sigma) {
    Fit_diffTm2(a, q, d, pred, c, parm1, parm2, Tm, Tm, y, w,
        sigma)
}

Diff1Tr = function(a, q, d, pred, c, parm1, parm2, Tr, y, w,
    sigma) {
    Fit_diffTr(a, q, d, pred, c, parm1, parm2, Tr, Tr, y, w,
        sigma)
}

Diff2Tr = function(a, q, d, pred, c, parm1, parm2, Tr, y, w,
    sigma) {
    Fit_diffTr2(a, q, d, pred, c, parm1, parm2, Tr, Tr, y, w,
        sigma)
}
```

```
# Transmission between two host genotypes differing in
# shoaling rate is the geometric mean, as mass action would
# suggest
trans_func = function(c1, c2, Ti) {
    sqrt(c1 * c2) * Ti
}

# Similarly, an invader's death from predation depends on the
# geometric mean of it and the resident's shoaling rates
# Subtracting 1 from this NGM fitness proxy so that the sign
# reflects invasibility
Host_fit_proxy = function(a, q, cm, cr, back_d, pred, parm1,
    parm2, Ti, y, w) {
    ((a - H_eq(a, q, host_death(back_d, pred, cr), trans_func(cr,
        cr, Ti), virulence_func(parm1, parm2, Ti), y, w) * q) *
        (host_death(back_d, pred, sqrt(cm * cr)) * virulence_func(parm1,
            parm2, Ti) + host_death(back_d, pred, sqrt(cm * cr)) *
            w + host_death(back_d, pred, sqrt(cm * cr)) * y +
            virulence_func(parm1, parm2, Ti) * w + w * y + host_death(back_d,
            pred, sqrt(cm * cr))^2 + trans_func(cm, cr, Ti) *
            I_eq(a, q, host_death(back_d, pred, cr), trans_func(cr,
                cr, Ti), virulence_func(parm1, parm2, Ti), y,
                w) * host_death(back_d, pred, sqrt(cm * cr)) +
            trans_func(cm, cr, Ti) * I_eq(a, q, host_death(back_d,
                pred, cr), trans_func(cr, cr, Ti), virulence_func(parm1,
                parm2, Ti), y, w) * w + trans_func(cm, cr, Ti) *
            I_eq(a, q, host_death(back_d, pred, cr), trans_func(cr,
                cr, Ti), virulence_func(parm1, parm2, Ti), y,
                w) * y))/(host_death(back_d, pred, sqrt(cm *
        cr))^2 * virulence_func(parm1, parm2, Ti) + host_death(back_d,
        pred, sqrt(cm * cr))^2 * w + host_death(back_d, pred,
        sqrt(cm * cr))^2 * y + host_death(back_d, pred, sqrt(cm *
        cr))^3 + trans_func(cm, cr, Ti) * I_eq(a, q, host_death(back_d,
        pred, cr), trans_func(cr, cr, Ti), virulence_func(parm1,
        parm2, Ti), y, w) * host_death(back_d, pred, sqrt(cm *
        cr))^2 + host_death(back_d, pred, sqrt(cm * cr)) * virulence_func(
           parm1,
        parm2, Ti) * w + host_death(back_d, pred, sqrt(cm * cr)) *
        w * y + trans_func(cm, cr, Ti) * I_eq(a, q, host_death(back_d,
        pred, cr), trans_func(cr, cr, Ti), virulence_func(parm1,
        parm2, Ti), y, w) * host_death(back_d, pred, sqrt(cm *
        cr)) * virulence_func(parm1, parm2, Ti) + trans_func(cm,
        cr, Ti) * I_eq(a, q, host_death(back_d, pred, cr), trans_func(cr,
        cr, Ti), virulence_func(parm1, parm2, Ti), y, w) * host_death(back_d,
        pred, sqrt(cm * cr)) * w + trans_func(cm, cr, Ti) * I_eq(a,
        q, host_death(back_d, pred, cr), trans_func(cr, cr, Ti),
        virulence_func(parm1, parm2, Ti), y, w) * host_death(back_d,
        pred, sqrt(cm * cr)) * y + trans_func(cm, cr, Ti) * I_eq(a,
        q, host_death(back_d, pred, cr), trans_func(cr, cr, Ti),
        virulence_func(parm1, parm2, Ti), y, w) * virulence_func(parm1,
        parm2, Ti) * w) - 1
}
```

```
# Take derivatives of the host fitness proxy wrt invader or
# resident trait
Fit_diffcm = Deriv(Host_fit_proxy, "cm")
Fit_diffcm2 = Deriv(Fit_diffcm, "cm")
Fit_diffcr = Deriv(Host_fit_proxy, "cr")
Fit_diffcr2 = Deriv(Fit_diffcr, "cr")

# Evaluate derivatives when Tm=Tr
Diff1cm = function(a, q, cm, back_d, pred, parm1, parm2, Ti,
    y, w) {
    Fit_diffcm(a, q, cm, cm, back_d, pred, parm1, parm2, Ti,
        y, w)
}

Diff2cm = function(a, q, cm, back_d, pred, parm1, parm2, Ti,
    y, w) {
    Fit_diffcm2(a, q, cm, cm, back_d, pred, parm1, parm2, Ti,
        y, w)
}

Diff1cr = function(a, q, cr, back_d, pred, parm1, parm2, Ti,
    y, w) {
    Fit_diffcr(a, q, cr, cr, back_d, pred, parm1, parm2, Ti,
        y, w)
}

Diff2cr = function(a, q, cr, back_d, pred, parm1, parm2, Ti,
    y, w) {
    Fit_diffcr2(a, q, cr, cr, back_d, pred, parm1, parm2, Ti,
        y, w)
}

# Make sure a resulting c CSS is feasible. Return NA if not.
feasible_c = function(a, q, d, c, parm1, parm2, Ti, y, w) {
    ifelse(length(c) > 0 && (d + virulence_func(parm1, parm2,
        Ti) + y)/(c * Ti) < (a - d)/q, as.numeric(c), NA)
}
```

##### 4.3 Write functions to find a range over which to calculate host and parasite evolution

```
# Define how much resolution with which to calculate host and
# parasite evolution.
grad_length = 50

# Calculate the T which maximizes R0
T_max_function <- function(d, pred, c, y, parm1, parm2) {
    ((host_death(d, pred, c) + y)/((parm2 - 1) * parm1))^(1/parm2)
}

# Calculate R0-1
R0_m1 <- function(a, q, d, pred, c, T_func, y, parm1, parm2) {
    c * T_func * (a - host_death(d, pred, c))/q/(host_death(d,
```

```
        pred, c) + virulence_func(parm1, parm2, T_func) + y) -
        1
}

# Find the maximum R0 attainable by parasites
R0_max_m1 <- function(a, q, d, pred, c, y, parm1, parm2) {
    R0_m1(a, q, d, pred, c, T_max_function(d, pred, c, y, parm1,
        parm2), y, parm1, parm2)
}

# Find the minimum c that supports parasites, given
# predation, the tradeoff, etc
c_min_function <- function(a_func, q_func, d_func, pred_func,
    y_func, parm1_func, parm2_func) {
    return(uniroot(R0_max_m1, interval = c(1e-07, 1000), a = a_func,
        q = q_func, d = d_func, pred = pred_func, y = y_func,
        parm1 = parm1_func, parm2 = parm2_func)$root)
}

# Calculate the lowest and highest T values that can survive
# given predation, c, etc.
T_extremes_function <- function(a_func, q_func, d_func, pred_func,
    c_func, y_func, parm1_func, parm2_func) {
    uniroot.all(R0_m1, c(1e-08, 100), n = 1e+06, a = a_func,
        q = q_func, d = d_func, pred = pred_func, c = c_func,
        y = y_func, parm1 = parm1_func, parm2 = parm2_func)
}
```

##### 4.4 Find intersections of CSS curves and determine if they meet the strong convergence stability criterion

```
# For the sake of coevolutionary convergence stability, will
# need to calculate the impact of host trait on parasite
# selection gradient and vice versa.
c_on_grad_T <- Deriv(Diff1Tm, "c")
T_on_grad_c <- Deriv(Diff1cm, "Ti")

# Check the strong convergence stability criterion
coCSS_check <- function(a, q, d, pred, c, parm1, parm2, Ti, y,
    w, sigma) {
    D2_cond <- c_on_grad_T(a, q, d, pred, c, parm1, parm2, Ti,
        y, w, sigma) * T_on_grad_c(a, q, c, d, pred, parm1, parm2,
        Ti, y, w)
    temp_out <- 0.5 * (Diff2cm(a, q, c, d, pred, parm1, parm2,
        Ti, y, w) - 0.5 * Diff2cr(a, q, c, d, pred, parm1, parm2,
        Ti, y, w)) * 0.5 * (Diff2Tm(a, q, d, pred, c, parm1,
        parm2, Ti, y, w, sigma) - Diff2Tr(a, q, d, pred, c, parm1,
        parm2, Ti, y, w, sigma)) - c_on_grad_T(a, q, d, pred,
        c, parm1, parm2, Ti, y, w, sigma) * T_on_grad_c(a, q,
        c, d, pred, parm1, parm2, Ti, y, w) - D2_cond

    # Return 1 if the strong convergence stability criterion for
    # being a coCSS is met. Otherwise, return 0.
```

```
    return(ifelse((D2_cond < 0 || temp_out > 0), 1, 0))
}

# Find the intersection of two curves and make sure it is a
# coCSS thanks to the strong convergence stability criterion
# Input parasite evolution first and keep virulence as the
# x-values and as the y-values

# Find the slope and intercept of a curve given two points
slope_int_finder <- function(pointx1, pointy1, pointx2, pointy2) {
    slope = (pointy2 - pointy1)/(pointx2 - pointx1)
    int = pointy2 - pointx2 * slope
    return(c(slope, int))
}

# Find the intersection of two lines given their slopes and
# intercepts
simple_intersection_finder <- function(parasite_slope, parasite_int,
    host_slope, host_int) {
    x_out = (host_int - parasite_int)/(parasite_slope - host_slope)
    y_out = x_out * host_slope + host_int
    return(c(x_out, y_out))
}

# Find the values closest to the intersection of a parasite
# evolution curve: x1(y1) and host evolution curve: y2(x2)
# Input other parameter values to enable checking of the
# strong convergence stability criterion
Intersection_closest = function(x1, y1, x2, y2, a, q, d, pred,
    parm1, parm2, y, w, sigma) {
    # Assumes both x vectors are same length and both y vectors
    # are same length

    # Calculate the slope and intercept from each two consecutive
    # points on the parasite curve
    parasite_slope_int = array(NA, dim = c((length(x1) - 1),
        2))

    # Same for host
    host_slope_int = array(NA, dim = c((length(x1) - 1), 2))
    for (i in 1:(length(x1) - 1)) {
        parasite_slope_int[i, ] = slope_int_finder(x1[i], y1[i],
            x1[i + 1], y1[i + 1])
        host_slope_int[i, ] = slope_int_finder(x2[i], y2[i],
            x2[i + 1], y2[i + 1])
    }

    # Calculate intersections of all lines defined by consecutive
    # points on host and parsite curves. First layer is x value
    # of intersection. Second is y value. Third is whether or not
    # these points fall within the range of their parent points.
    # Rows loop through parasite points. Columns loop through
    # host points.
    intersections <- array(NA, dim = c((length(x1) - 1), (length(x1) -
```

```
        1), 3))
    for (i in 1:(length(x1) - 1)) {
        for (j in 1:(length(x1) - 1)) {
            intersections[i, j, 1:2] <- simple_intersection_finder(parasite_
                slope_int[i,
                 1], parasite_slope_int[i, 2], host_slope_int[j,
                 1], host_slope_int[j, 2])

            # This will be 1 when points fall within the range of their
            # parent points and are thus an approximation of the
            # intersection of the two curves
            intersections[i, j, 3] <- as.numeric(!is.na(intersections[i,
                j, 1] * intersections[i, j, 2]) && intersections[i,
                j, 1] >= max(c(x1[i], x2[j])) && intersections[i,
                j, 1] <= min(c(x1[i + 1], x2[j + 1])))
        }
    }

    # Indices of the intersection values.
    parasite_cand <- which(intersections[, , 3] == 1, arr.ind = T)[1]
    host_cand <- which(intersections[, , 3] == 1, arr.ind = T)[2]

    # Check strong convergence
    coCSS_status <- coCSS_check(a, q, d, pred, intersections[parasite_cand,
        host_cand, 2], parm1, parm2, inv_virulence_func(parm1,
        parm2, intersections[parasite_cand, host_cand, 1]), y,
        w, sigma)

    # If strong convergence is satisfied...
    if (coCSS_status == 1) {
        # ...returns v_min, v_max, c_min, c_max for new, refined
        # search
        return(c(min(c(x1[parasite_cand], x2[host_cand])), max(c(x1[parasite_
            cand +
             1], x2[host_cand + 1])), c(y1[parasite_cand], y2[host_cand])[which
                .min(c(x1[parasite_cand],
             x2[host_cand]))], c(y1[parasite_cand + 1], y2[host_cand +
             1])[which.max(c(x1[parasite_cand + 1], x2[host_cand +
             1]))]))
    } else {
        print("Not␣exactly␣one,␣definite␣coCSS␣intersection")
    }
}

# Same as above but actually estimate the intersection
Intersection_finder = function(x1, y1, x2, y2, a, q, d, pred,
    parm1, parm2, y, w, sigma) {

    # Assumes both x vectors are same length and both y vectors
    # are same length
    parasite_slope_int = array(NA, dim = c((length(x1) - 1),
        2))
    host_slope_int = array(NA, dim = c((length(x1) - 1), 2))
    for (i in 1:(length(x1) - 1)) {
```

```
        parasite_slope_int[i, ] = slope_int_finder(x1[i], y1[i],
            x1[i + 1], y1[i + 1])
        host_slope_int[i, ] = slope_int_finder(x2[i], y2[i],
            x2[i + 1], y2[i + 1])
    }

    # First layer is x value of intersection. Second is y value.
    # Third is whether or not these points fall within their
    # parent points. Rows loop through parasite points. Columns
    # loop through host points.
    intersections <- array(NA, dim = c((length(x1) - 1), (length(x1) -
        1), 3))
    for (i in 1:(length(x1) - 1)) {
        for (j in 1:(length(x1) - 1)) {
            intersections[i, j, 1:2] <- simple_intersection_finder(parasite_
                slope_int[i,
                 1], parasite_slope_int[i, 2], host_slope_int[j,
                 1], host_slope_int[j, 2])
            intersections[i, j, 3] <- as.numeric(!is.na(intersections[i,
                j, 1] * intersections[i, j, 2]) && intersections[i,
                j, 1] >= max(c(x1[i], x2[j])) && intersections[i,
                j, 1] <= min(c(x1[i + 1], x2[j + 1])))
        }
    }
    # Indices of the intersection values
    parasite_cand <- which(intersections[, , 3] == 1, arr.ind = T)[1]
    host_cand <- which(intersections[, , 3] == 1, arr.ind = T)[2]

    coCSS_status <- coCSS_check(a, q, d, pred, intersections[parasite_cand,
        host_cand, 2], parm1, parm2, inv_virulence_func(parm1,
        parm2, intersections[parasite_cand, host_cand, 1]), y,
        w, sigma)

    # If the strong convergence stability criterion is
    # satisfied...
    if (coCSS_status == 1) {
        # return the best approximation of the intersection
        return(c(intersections[parasite_cand, host_cand, 1],
            intersections[parasite_cand, host_cand, 2]))
    } else {
        print("Not exactly one, definite coCSS intersection")
    }
}
```

##### 4.5 Define a function that calculates coevolution given input parameters

This function will take parameters as inputs and give the key outputs of coevolution at a lower value of predation, a higher one, and their relationship for some variables

```
key_coevo_outputs <- function(parms) {

    # Separate the single input vector into various, named, input
    # parameters
```

```
a_ex = parms[1]
q_ex = parms[2]
d_ex = parms[3]
pred1 = parms[4]
pred2 = parms[5]
parm1_ex = parms[6]
parm2_ex = parms[7]
y_ex = parms[8]
w_ex = parms[9]
sigma_ex = parms[10]

# Set a minimum value of c based on the smallest c that can
# support parasites with T,v that optimizes R0
c_min1 <- c_min_function(a_ex, q_ex, d_ex, pred1, y_ex, parm1_ex,
    parm2_ex)
# Set the maximum value of c arbitrarily 50x greater than
# that.
c_max1 <- c_min1 * 50

# Do this independently for the second predation value
c_min2 <- c_min_function(a_ex, q_ex, d_ex, pred2, y_ex, parm1_ex,
    parm2_ex)
# Set the maximum value of c arbitrarily 50x greater than
# that.
c_max2 <- c_min2 * 50

# Set the extreme values of transmissibility based on the
# smallest and largest values of T that can survive the ideal
# conditions of maximum shoaling rate
T_extremes1 <- T_extremes_function(a_ex, q_ex, d_ex, pred1,
    c_max1, y_ex, parm1_ex, parm2_ex)
T_min1 <- T_extremes1[1]
T_max1 <- T_extremes1[2]

T_extremes2 <- T_extremes_function(a_ex, q_ex, d_ex, pred2,
    c_max2, y_ex, parm1_ex, parm2_ex)
T_min2 <- T_extremes2[1]
T_max2 <- T_extremes2[2]

# Start with a broad range of v and c The v range is
# logarithmically spaced in T
v_range_initial1 <- virulence_func(parm1_ex, parm2_ex, 10^linspace(log10(T
   _min1),
    log10(T_max1), grad_length))
c_range_initial1 <- 10^linspace(log10(c_min1 * 1.01), log10(c_max1),
    grad_length)

v_range_initial2 <- virulence_func(parm1_ex, parm2_ex, 10^linspace(log10(T
   _min2),
    log10(T_max2), grad_length))
c_range_initial2 <- 10^linspace(log10(c_min2 * 1.01), log10(c_max2),
    grad_length)

# Define the CSS functions to search within the chosen ranges
```

```
# Write a function to find the c value hosts evolve given
# paramters. Very similar to evo_checkedroots
host_evo_checkedroots1 = function(a_func, q_func, back_d_func,
    pred_func, parm1_func, parm2_func, Ti_func, y_func, w_func) {
    # Find root of the fitness gradient
    roots_storage = uniroot.all(Diff1cm, c(c_min1, c_max1),
        tol = 1e-06, n = 500, a = a_func, q = q_func, back_d = back_d_func
            ,
        pred = pred_func, parm1 = parm1_func, parm2 = parm2_func,
        Ti = Ti_func, y = y_func, w = w_func)

    # Evaluate stability of those roots
    cchecks = array(NA, dim = c(length(roots_storage), 2))
    if (length(roots_storage) > 0) {
        for (j in 1:length(roots_storage)) {
            # Evolutionary stability
            cchecks[j, 1] = Diff2cm(a_func, q_func, roots_storage[j],
              back_d_func, pred_func, parm1_func, parm2_func,
              Ti_func, y_func, w_func)
            # Convergence stability
            cchecks[j, 2] = Diff2cr(a_func, q_func, roots_storage[j],
              back_d_func, pred_func, parm1_func, parm2_func,
              Ti_func, y_func, w_func) - Diff2cm(a_func,
              q_func, roots_storage[j], back_d_func, pred_func,
              parm1_func, parm2_func, Ti_func, y_func, w_func)
        }
    }
    # Only return CSSes
    temp_out = roots_storage[which(cchecks[, 1] < 0 & cchecks[,
        2] > 0)]
    out = feasible_c(a_func, q_func, host_death(back_d_func,
        pred_func, temp_out), temp_out, parm1_func, parm2_func,
        Ti_func, y_func, w_func)

    # If no CSS was found but the fitness gradient is always
    # pointing up or down, return the maximum or minimum value of
    # c
    if (length(which(!is.na(out))) < 1 && length(which(mapply(Diff1cm,
        a = a_func, q = q_func, back_d = back_d_func, pred = pred_func,
        parm1 = parm1_func, parm2 = parm2_func, Ti = Ti_func,
        y = y_func, w = w_func, cm = linspace(c_min1, c_max1,
            100)) > 0)) == 100) {
        out = c_max1
    }
    if (length(which(!is.na(out))) < 1 && length(which(mapply(Diff1cm,
        a = a_func, q = q_func, back_d = back_d_func, pred = pred_func,
        parm1 = parm1_func, parm2 = parm2_func, Ti = Ti_func,
        y = y_func, w = w_func, cm = linspace(c_min1, c_max1,
            100)) < 0)) == 100) {
        out = c_min1
    }
    return(out)
}
```

```
# Same but for the second predation value
host_evo_checkedroots2 = function(a_func, q_func, back_d_func,
    pred_func, parm1_func, parm2_func, Ti_func, y_func, w_func) {
    roots_storage = uniroot.all(Diff1cm, c(c_min2, c_max2),
        tol = 1e-06, n = 500, a = a_func, q = q_func, back_d = back_d_func
          ,
        pred = pred_func, parm1 = parm1_func, parm2 = parm2_func,
        Ti = Ti_func, y = y_func, w = w_func)
    cchecks = array(NA, dim = c(length(roots_storage), 2))
    if (length(roots_storage) > 0) {
        for (j in 1:length(roots_storage)) {
            cchecks[j, 1] = Diff2cm(a_func, q_func, roots_storage[j],
              back_d_func, pred_func, parm1_func, parm2_func,
              Ti_func, y_func, w_func)
            cchecks[j, 2] = Diff2cr(a_func, q_func, roots_storage[j],
              back_d_func, pred_func, parm1_func, parm2_func,
              Ti_func, y_func, w_func) - Diff2cm(a_func,
              q_func, roots_storage[j], back_d_func, pred_func,
              parm1_func, parm2_func, Ti_func, y_func, w_func)
        }
    }
    temp_out = roots_storage[which(cchecks[, 1] < 0 & cchecks[,
        2] > 0)]
    out = feasible_c(a_func, q_func, host_death(back_d_func,
        pred_func, temp_out), temp_out, parm1_func, parm2_func,
        Ti_func, y_func, w_func)
    if (length(which(!is.na(out))) < 1 && length(which(mapply(Diff1cm,
        a = a_func, q = q_func, back_d = back_d_func, pred = pred_func,
        parm1 = parm1_func, parm2 = parm2_func, Ti = Ti_func,
        y = y_func, w = w_func, cm = linspace(c_min2, c_max2,
            100)) > 0)) == 100) {
        out = c_max2
    }
    if (length(which(!is.na(out))) < 1 && length(which(mapply(Diff1cm,
        a = a_func, q = q_func, back_d = back_d_func, pred = pred_func,
        parm1 = parm1_func, parm2 = parm2_func, Ti = Ti_func,
        y = y_func, w = w_func, cm = linspace(c_min2, c_max2,
            100)) < 0)) == 100) {
        out = c_min2
    }
    return(out)
}

# Write a function that finds CSS value(s) given parameters.
# In simpler cases, there is only one CSS for a given set of
# parameter values
evo_checkedroots = function(a_func, q_func, d_func, pred_func,
    c_func, parm1_func, parm2_func, y_func, w_func, sigma_func) {
    # Find the talues of Tm that make the Diff1Tm=0
    roots_storage = uniroot.all(Diff1Tm, c(1e-07, 100), tol = 1e-06,
        n = 500, a = a_func, q = q_func, d = d_func, pred = pred_func,
        c = c_func, parm1 = parm1_func, parm2 = parm2_func,
        y = y_func, w = w_func, sigma = sigma_func)
    Tchecks = array(NA, dim = c(length(roots_storage), 2))
```

```
    for (j in 1:length(roots_storage)) {
        # Evolutionary stability criterion
        Tchecks[j, 1] = Diff2Tm(a_func, q_func, d_func, pred_func,
            c_func, parm1_func, parm2_func, roots_storage[j],
            y_func, w_func, sigma_func)
        # Convergence stability criterion
        Tchecks[j, 2] = Diff2Tr(a_func, q_func, d_func, pred_func,
            c_func, parm1_func, parm2_func, roots_storage[j],
            y_func, w_func, sigma_func) - Diff2Tm(a_func,
            q_func, d_func, pred_func, c_func, parm1_func,
            parm2_func, roots_storage[j], y_func, w_func,
            sigma_func)
    }
    # Return the value(s) of Tm that satisfy the derivative
    # conditions
    temp_out = roots_storage[which(Tchecks[, 1] < 0 & Tchecks[,
        2] > 0)]
    return(temp_out)
}

# Find out shoaling rates hosts will evolve, need at least a
# little predators or else hosts just evolve no
host_evo_v1_initial = mapply(host_evo_checkedroots1, a_func = a_ex,
    q_func = q_ex, back_d_func = d_ex, pred_func = pred1,
    parm1 = parm1_ex, parm2 = parm2_ex, Ti_func = inv_virulence_func(parm1
       _ex,
        parm2_ex, v_range_initial1), y_func = y_ex, w_func = w_ex)
# Calculate parasite evolution over range of shoaling rates
parasite_evo_c1_initial = mapply(evo_checkedroots, a_func = a_ex,
    q_func = q_ex, d_func = d_ex, pred = pred1, c_func = c_range_initial1,
    parm1 = parm1_ex, parm2 = parm2_ex, y_func = y_ex, w_func = w_ex,
    sigma_func = sigma_ex)

# Same but for more predation
host_evo_v2_initial = mapply(host_evo_checkedroots2, a_func = a_ex,
    q_func = q_ex, back_d_func = d_ex, pred_func = pred2,
    parm1 = parm1_ex, parm2 = parm2_ex, Ti_func = inv_virulence_func(parm1
       _ex,
        parm2_ex, v_range_initial2), y_func = y_ex, w_func = w_ex)
parasite_evo_c2_initial = mapply(evo_checkedroots, a_func = a_ex,
    q_func = q_ex, d_func = d_ex, pred = pred2, c_func = c_range_initial2,
    parm1 = parm1_ex, parm2 = parm2_ex, y_func = y_ex, w_func = w_ex,
    sigma_func = sigma_ex)

# Find the points closest to the intersection in order to
# define a newer, parameter range that focuses more on the
# area around the intersections.
low_pred_temp <- Intersection_closest(v_range_initial1, host_evo_v1_
    initial,
    virulence_func(parm1_ex, parm2_ex, parasite_evo_c1_initial),
    c_range_initial1, a_ex, q_ex, d_ex, pred1, parm1_ex,
    parm2_ex, y_ex, w_ex, sigma_ex)
high_pred_temp <- Intersection_closest(v_range_initial2,
    host_evo_v2_initial, virulence_func(parm1_ex, parm2_ex,
```

```
        parasite_evo_c2_initial), c_range_initial2, a_ex,
    q_ex, d_ex, pred2, parm1_ex, parm2_ex, y_ex, w_ex, sigma_ex)

c_min_temp1 <- min(low_pred_temp[3:4])
c_min_temp2 <- min(high_pred_temp[3:4])
c_max_temp1 <- max(low_pred_temp[3:4])
c_max_temp2 <- max(high_pred_temp[3:4])
v_min_temp1 <- min(low_pred_temp[1:2])
v_max_temp1 <- max(low_pred_temp[1:2])
v_min_temp2 <- min(high_pred_temp[1:2])
v_max_temp2 <- max(high_pred_temp[1:2])

# Then make the range 2 times broader but centered the same
# and make sure it can't go too low, e.g., negative, and no
# need to go all the way up to the max
c_min_new1 <- max(c((c_min_temp1 + c_max_temp1)/2 - (c_max_temp1 -
    c_min_temp1), c_min1 * 1.01))
c_max_new1 <- min(c((c_min_temp1 + c_max_temp1)/2 + (c_max_temp1 -
    c_min_temp1), c_max1 * 0.99))
c_min_new2 <- max(c((c_min_temp2 + c_max_temp2)/2 - (c_max_temp2 -
    c_min_temp2), c_min2 * 1.01))
c_max_new2 <- min(c((c_min_temp2 + c_max_temp2)/2 + (c_max_temp2 -
    c_min_temp2), c_max2 * 0.99))
v_min_new1 <- max(c((v_min_temp1 + v_max_temp1)/2 - (v_max_temp1 -
    v_min_temp1), virulence_func(parm1_ex, parm2_ex, T_min1 *
    1.01)))
v_max_new1 <- min(c((v_min_temp1 + v_max_temp1)/2 + (v_max_temp1 -
    v_min_temp1), virulence_func(parm1_ex, parm2_ex, T_max1) *
    0.99))
v_min_new2 <- max(c((v_min_temp2 + v_max_temp2)/2 - (v_max_temp2 -
    v_min_temp2), virulence_func(parm1_ex, parm2_ex, T_min2 *
    1.01)))
v_max_new2 <- min(c((v_min_temp2 + v_max_temp2)/2 + (v_max_temp2 -
    v_min_temp2), virulence_func(parm1_ex, parm2_ex, T_max2) *
    0.99))

# Make the final c_range and v_range
c_range1 <- linspace(c_min_new1, c_max_new1, grad_length)
v_range1 <- linspace(v_min_new1, v_max_new1, grad_length)
c_range2 <- linspace(c_min_new2, c_max_new2, grad_length)
v_range2 <- linspace(v_min_new2, v_max_new2, grad_length)

# Find out shoaling rates hosts will evolve, need at least a
# little predators or else hosts just evolve no shoaling
host_evo_v1 = mapply(host_evo_checkedroots1, a_func = a_ex,
    q_func = q_ex, back_d_func = d_ex, pred_func = pred1,
    parm1 = parm1_ex, parm2 = parm2_ex, Ti_func = inv_virulence_func(parm1
        _ex,
        parm2_ex, v_range1), y_func = y_ex, w_func = w_ex)
# Calculate parasite evolution over range of shoaling rates
parasite_evo_c1 = mapply(evo_checkedroots, a_func = a_ex,
    q_func = q_ex, d_func = d_ex, pred = pred1, c_func = c_range1,
    parm1 = parm1_ex, parm2 = parm2_ex, y_func = y_ex, w_func = w_ex,
    sigma_func = sigma_ex)
```

```
# Same but for more predation
host_evo_v2 = mapply(host_evo_checkedroots2, a_func = a_ex,
    q_func = q_ex, back_d_func = d_ex, pred_func = pred2,
    parm1 = parm1_ex, parm2 = parm2_ex, Ti_func = inv_virulence_func(parm1
       _ex,
        parm2_ex, v_range2), y_func = y_ex, w_func = w_ex)
parasite_evo_c2 = mapply(evo_checkedroots, a_func = a_ex,
    q_func = q_ex, d_func = d_ex, pred = pred2, c_func = c_range2,
    parm1 = parm1_ex, parm2 = parm2_ex, y_func = y_ex, w_func = w_ex,
    sigma_func = sigma_ex)

# Find the intersections. The way intersection_finder works
# guarantees this will be a coCSS. These are the
# coevolutionary values of virulence and shoaling rate at low
# and high predation respectively
v_int1 = Intersection_finder(v_range1, host_evo_v1, virulence_func(parm1_
    ex,
    parm2_ex, parasite_evo_c1), c_range1, a_ex, q_ex, d_ex,
    pred1, parm1_ex, parm2_ex, y_ex, w_ex, sigma_ex)[1]
c_int1 = Intersection_finder(v_range1, host_evo_v1, virulence_func(parm1_
    ex,
    parm2_ex, parasite_evo_c1), c_range1, a_ex, q_ex, d_ex,
    pred1, parm1_ex, parm2_ex, y_ex, w_ex, sigma_ex)[2]

v_int2 = Intersection_finder(v_range2, host_evo_v2, virulence_func(parm1_
    ex,
    parm2_ex, parasite_evo_c2), c_range2, a_ex, q_ex, d_ex,
    pred2, parm1_ex, parm2_ex, y_ex, w_ex, sigma_ex)[1]
c_int2 = Intersection_finder(v_range2, host_evo_v2, virulence_func(parm1_
    ex,
    parm2_ex, parasite_evo_c2), c_range2, a_ex, q_ex, d_ex,
    pred2, parm1_ex, parm2_ex, y_ex, w_ex, sigma_ex)[2]

v_ratio <- v_int2/v_int1
c_ratio <- c_int2/c_int1

# Calculate important outputs based on these coevolutionary
# points. Will often calcualte values at low predation, high
# predation, and their ratio.

# Prevalence
prev1 <- prev_eq(a_ex, q_ex, host_death(d_ex, pred1, c_int1),
    inv_virulence_func(parm1_ex, parm2_ex, v_int1) * c_int1,
    v_int1, y_ex, w_ex)
prev2 <- prev_eq(a_ex, q_ex, host_death(d_ex, pred2, c_int2),
    inv_virulence_func(parm1_ex, parm2_ex, v_int2) * c_int2,
    v_int2, y_ex, w_ex)
prev_ratio <- prev2/prev1

# Host density
H1 <- H_eq(a_ex, q_ex, host_death(d_ex, pred1, c_int1), inv_virulence_func
    (parm1_ex,
    parm2_ex, v_int1) * c_int1, v_int1, y_ex, w_ex)
```

```
    H2 <- H_eq(a_ex, q_ex, host_death(d_ex, pred2, c_int2), inv_virulence_func
        (parm1_ex,
        parm2_ex, v_int2) * c_int2, v_int2, y_ex, w_ex)
    H_ratio <- H2/H1

    # Overall host mortality rate from all sources
    mort1 <- host_death(d_ex, pred1, c_int1) + v_int1 * prev1
    mort2 <- host_death(d_ex, pred2, c_int2) + v_int2 * prev2
    mort_ratio <- mort2/mort1

    # How much of the change in mortality is due to parasites?
    mort_change_par = (v_int2 * prev2 - v_int1 * prev1)/(mort2 -
        mort1)

    # Transmission rate
    trans1 <- c_int1 * inv_virulence_func(parm1_ex, parm2_ex,
        v_int1)
    trans2 <- c_int2 * inv_virulence_func(parm1_ex, parm2_ex,
        v_int2)
    trans_ratio <- trans2/trans1

    # What proportion of the increase in virulence is due to
    # sociality? Calculate as 1-proportion due to predators
    # Calculate proportion due to predators as the mean of the
    # difference between the high predation curve at low shoaling
    # and the v_int1 and the difference between v_int2 and the
    # low predation curve at high shoaling

    c_range_wide <- linspace(c_int1 - 0.2 * c_int1, c_int2 +
        0.2 * c_int1, grad_length)
    parasite_wide_1 <- mapply(evo_checkedroots, a_func = a_ex,
        q_func = q_ex, d_func = d_ex, pred = pred1, c_func = c_range_wide,
        parm1 = parm1_ex, parm2 = parm2_ex, y_func = y_ex, w_func = w_ex,
        sigma_func = sigma_ex)
    parasite_wide_2 <- mapply(evo_checkedroots, a_func = a_ex,
        q_func = q_ex, d_func = d_ex, pred = pred2, c_func = c_range_wide,
        parm1 = parm1_ex, parm2 = parm2_ex, y_func = y_ex, w_func = w_ex,
        sigma_func = sigma_ex)

    vir_change_soc = 1 - mean(c(virulence_func(parm1_ex, parm2_ex,
        parasite_wide_2[which.min(abs(c_range_wide - c_int1))]) -
        v_int1, v_int2 - virulence_func(parm1_ex, parm2_ex, parasite_wide_1[
            which.min(abs(c_range_wide -
        c_int2))])), na.rm = T)/(v_int2 - v_int1)

    # Output these results
    return(c(prev1, prev2, prev_ratio, H1, H2, H_ratio, v_int1,
        v_int2, v_ratio, c_int1, c_int2, c_ratio, mort1, mort2,
        mort_ratio, mort_change_par, trans1, trans2, trans_ratio,
        vir_change_soc))
}

# As an example, this calculation is performed for some
# example parameter values
```

```
parms_focal <- c(0.1055767, 0.0169234, 0.001297164, 0.00739921,
    0.03299583, 1377273, 3.607885, 0.02027253, 0.03272097, 1.207408)
key_coevo_outputs(parms_focal)

# Calculate the strength of pathways
Pardiff_mort_c = Deriv(host_death, "c")
Pardiff_mort_pred = Deriv(host_death, "pred")
I_eq_long = function(a, q, d, pred, c, Ti, v, y, w) {
    I_eq(a, q, host_death(d, pred, c), c * Ti, v, y, w)
}
Pardiff_Ir_c = Deriv(I_eq_long, "c")
Pardiff_Ir_pred = Deriv(I_eq_long, "pred")

outputs_and_paths <- function(parms) {
    output1 <- key_coevo_outputs(parms)
    parms2 <- parms * c(1, 1, 1, 1.01, 0.99, 1, 1, 1, 1, 1)
    output2 <- key_coevo_outputs(parms2)

    dcdPred_low = (output2[10] - output1[10])/(parms2[4] - parms[4])
    dcdPred_hi = (output2[11] - output1[11])/(parms2[5] - parms[5])

    parmort_c_low = -parms[4]/output1[10]^2
    parmort_c_hi = -parms[5]/output1[11]^2

    parmort_pred_low = 1/output1[10]
    parmort_pred_hi = 1/output1[11]

    parsup_c_low = parms[10] * I_eq(parms[1], parms[2], host_death(parms[3],
        parms[4], output1[10]), output1[10] * inv_virulence_func(parms[6],
        parms[7], output1[7]), output1[7], parms[8], parms[9])
    parsup_c_hi = parms[10] * I_eq(parms[1], parms[2], host_death(parms[3],
        parms[5], output1[11]), output1[11] * inv_virulence_func(parms[6],
        parms[7], output1[8]), output1[8], parms[8], parms[9])

    parsup_Ir_low = output1[10] * parms[10]
    parsup_Ir_hi = output1[11] * parms[10]

    parIr_c_low = Pardiff_Ir_c(parms[1], parms[2], parms[3],
        parms[4], output1[10], inv_virulence_func(parms[6], parms[7],
            output1[7]), output1[7], parms[8], parms[9])
    parIr_c_hi = Pardiff_Ir_c(parms[1], parms[2], parms[3], parms[5],
        output1[11], inv_virulence_func(parms[6], parms[7], output1[8]),
        output1[8], parms[8], parms[9])

    parIr_pred_low = Pardiff_Ir_pred(parms[1], parms[2], parms[3],
        parms[4], output1[10], inv_virulence_func(parms[6], parms[7],
            output1[7]), output1[7], parms[8], parms[9])
    parIr_pred_hi = Pardiff_Ir_pred(parms[1], parms[2], parms[3],
        parms[5], output1[11], inv_virulence_func(parms[6], parms[7],
            output1[8]), output1[8], parms[8], parms[9])

    pathway1 = mean(c(virulence_deriv(parms[6], parms[7], inv_virulence_func(
       parms[6],
        parms[7], output1[7])) * -1/Diff2Tm(parms[1], parms[2],
```

```
      parms[3], parms[4], output1[10], parms[6], parms[7],
      inv_virulence_func(parms[6], parms[7], output1[7]), parms[8],
      parms[9], parms[10]) * 1/inv_virulence_func(parms[6],
      parms[7], output1[7]) * parmort_pred_low, virulence_deriv(parms[6],
      parms[7], inv_virulence_func(parms[6], parms[7], output1[8])) *
      -1/Diff2Tm(parms[1], parms[2], parms[3], parms[5], output1[11],
      parms[6], parms[7], inv_virulence_func(parms[6], parms[7],
          output1[8]), parms[8], parms[9], parms[10]) * 1/inv_virulence_func
             (parms[6],
      parms[7], output1[8]) * parmort_pred_hi))

  pathway2 = mean(c(virulence_deriv(parms[6], parms[7], inv_virulence_func(
     parms[6],
      parms[7], output1[7])) * -1/Diff2Tm(parms[1], parms[2],
      parms[3], parms[4], output1[10], parms[6], parms[7],
      inv_virulence_func(parms[6], parms[7], output1[7]), parms[8],
      parms[9], parms[10]) * parsup_Ir_low * parIr_pred_low,
      virulence_deriv(parms[6], parms[7], inv_virulence_func(parms[6],
          parms[7], output1[8])) * -1/Diff2Tm(parms[1], parms[2],
          parms[3], parms[5], output1[11], parms[6], parms[7],
          inv_virulence_func(parms[6], parms[7], output1[8]),
          parms[8], parms[9], parms[10]) * parsup_Ir_hi * parIr_pred_hi))

  pathway3 = mean(c(virulence_deriv(parms[6], parms[7], inv_virulence_func(
     parms[6],
      parms[7], output1[7])) * -1/Diff2Tm(parms[1], parms[2],
      parms[3], parms[4], output1[10], parms[6], parms[7],
      inv_virulence_func(parms[6], parms[7], output1[7]), parms[8],
      parms[9], parms[10]) * 1/inv_virulence_func(parms[6],
      parms[7], output1[7]) * parmort_c_low * dcdPred_low,
      virulence_deriv(parms[6], parms[7], inv_virulence_func(parms[6],
          parms[7], output1[8])) * -1/Diff2Tm(parms[1], parms[2],
          parms[3], parms[5], output1[11], parms[6], parms[7],
          inv_virulence_func(parms[6], parms[7], output1[8]),
          parms[8], parms[9], parms[10]) * 1/inv_virulence_func(parms[6],
          parms[7], output1[8]) * parmort_c_hi * dcdPred_hi))

  pathway4 = mean(c(virulence_deriv(parms[6], parms[7], inv_virulence_func(
     parms[6],
      parms[7], output1[7])) * -1/Diff2Tm(parms[1], parms[2],
      parms[3], parms[4], output1[10], parms[6], parms[7],
      inv_virulence_func(parms[6], parms[7], output1[7]), parms[8],
      parms[9], parms[10]) * (parsup_c_low + parsup_Ir_low *
      parIr_c_low) * dcdPred_low, virulence_deriv(parms[6],
      parms[7], inv_virulence_func(parms[6], parms[7], output1[8])) *
      -1/Diff2Tm(parms[1], parms[2], parms[3], parms[5], output1[11],
      parms[6], parms[7], inv_virulence_func(parms[6], parms[7],
          output1[8]), parms[8], parms[9], parms[10]) * (parsup_c_hi +
      parsup_Ir_hi * parIr_c_hi) * dcdPred_hi))

  return(c(output1, pathway1, pathway2, pathway3, pathway4))
}
```

#### 5 Evolutionary algorithm to find parameter values for the theoretical model

```
# Create an array of standards for outputs
output_standards = array(NA, dim = 20)
output_standards[1] = 0.3105011
output_standards[2] = 0.3942108
# prev_ratio
output_standards[3] = 1.269596
# H1
output_standards[4] = 8.188143
# H2
output_standards[5] = 4.675556
# H_ratio
output_standards[6] = 0.5710154
# c_ratio point estimate = 1.9275381.
output_standards[12] = 1.9275381
# Field mortality digitized from Reznick
output_standards[13] = 0.01065757
output_standards[14] = 0.02637546
# Mort_ratio
output_standards[15] = 2.47481
# trans1 should be 0.014447744 based on Upper Aripo estimates
# of transmission rate from 'Are host-parasite interactions
# influenced by adaptation to predators? A test with guppies
# and Gyrodactylus in experimental stream channels' see
# SI_cT_estimation.R
output_standards[17] = 0.014447744

tradeoff_calculator <- function(d_func) {
    partial_virulence <- curve_data[, 1] - d_func
    partial_virulence[which(partial_virulence < 0)] <- 0
    temp_glm <- glm(partial_virulence ~ log(curve_data[, 2]),
        family = Gamma(link = log))
    return(c(exp(as.numeric(temp_glm$coefficients)[1]), as.numeric(temp_glm$
       coefficients)[2]))
}
tradeoff_calculator2 <- function(d_func) {
    partial_virulence <- curve_data2[, 1] - d_func
    partial_virulence[which(partial_virulence < 0)] <- 0
    temp_glm <- glm(partial_virulence ~ log(curve_data2[, 2]),
        family = Gamma(link = log))
    return(c(exp(as.numeric(temp_glm$coefficients)[1]), as.numeric(temp_glm$
       coefficients)[2]))
}

# Number of 'genotypes' per generation of the evolutionary
# algorithm and maximum number of generations for which the
# algorithm is run
num_genotypes <- 300
num_generations <- 20

input_standards <- array(NA, dim = 10)
```

```r
# Training values for a
input_standards[1] = 0.1062086
# none for q d
input_standards[3] = 0.001409638
# None for pred1, pred2, parm1, parm2

# recovery rate
input_standards[8] = 0.03279148
# waning immunity
input_standards[9] = log(0.5)/-21

# Size of mutations of parameter sets in the algorithm
mutation_size = 0.05

# Starting values, can use tradeoff_calculator2 instead
parm_start <- c(input_standards[1], 0.01, input_standards[3],
    0.01, 0.03, tradeoff_calculator(input_standards[3])[1], tradeoff_
        calculator(input_standards[3])[2],
    input_standards[8], input_standards[9], 0.5)

# parm_start<-c(input_standards[1],1e-2,
# input_standards[3],1e-2, #3e-2,tradeoff_calculator2
# (input_standards[3])[1],
# #tradeoff_calculator2(input_standards[3])[2],
# input_standards[8], input_standards[9],0.5)

# Create an array to store the outputs
key_outputs <- array(NA, dim = c(num_genotypes, 20, num_generations))

# Create an array to hold parameters over the 'generations'
# of the algorithm
parms_thisgen = array(NA, dim = c(num_genotypes, 10, (num_generations +
    1)))

# Initialize with starting values
parms_thisgen[1, , 1] <- parm_start

# Track the best scores across generations
best_scores <- array(NA, dim = num_generations)

for (i in 1:num_generations) {
    # Mutate the parameters based on last round's winner
    for (j in 2:num_genotypes) {
        parms_thisgen[j, c(1, 2, 3, 4, 5, 8, 9, 10), i] <- parms_thisgen[1,
            c(1, 2, 3, 4, 5, 8, 9, 10), i] * 10^rnorm(8, 0, mutation_size)
        parms_thisgen[j, c(6, 7), i] <- tradeoff_calculator(parms_thisgen[j,
            3, i])
    }

    # Calculate results from each strategy The errors I've
    # investigated so far stem from hosts not evolving to a CSS
    # but to max c instead if pred is too high.
    for (j in 1:num_genotypes) {
```

```
        parm_temp <- NA
        tryCatch({
            parm_temp <- c(parms_thisgen[j, , i])
            key_outputs[j, , i] <- key_coevo_outputs(parm_temp)
        }, error = function(e) {
            cat("ERROR:", conditionMessage(e), "\n")
        })
        print(i)
        print(j)
    }

    # Score each genotype
    genotype_out_ind_scores <- array(NA, dim = c(num_genotypes,
        20))
    genotype_in_ind_scores <- array(NA, dim = c(num_genotypes,
        10))
    genotype_sum_scores <- array(NA, dim = c(num_genotypes))
    for (j in 1:num_genotypes) {
        for (m in which(!is.na(output_standards))) {
            if (is.na(key_outputs[j, m, i])) {
                genotype_out_ind_scores[j, m] <- 1e+05
            } else {
                genotype_out_ind_scores[j, m] <- abs(key_outputs[j,
                  m, i] - output_standards[m])/output_standards[m]
            }
        }
        # Score inputs
        for (n in c(1, 3, 8, 9, 10)) {
            genotype_in_ind_scores[j, n] <- abs(parms_thisgen[j,
                n, i] - input_standards[n])/input_standards[n]
        }
        genotype_sum_scores[j] <- sum(genotype_out_ind_scores[j,
            ], na.rm = T) + sum(genotype_in_ind_scores[j, ],
            na.rm = T)
    }
    best_index <- NA
    best_index <- which.min(genotype_sum_scores)
    parms_thisgen[1, , i + 1] <- parms_thisgen[best_index, ,
        i]
    current_best_score <- min(genotype_sum_scores[best_index],
        best_scores, na.rm = T)
    best_scores[i] <- current_best_score
}

# The best fit parameter values
parms_focal <- parms_thisgen[best_index, , num_generations]
key_outputs[best_index, , num_generations]

# Average accuracy of fitting:
# genotype_sum_scores[best_index]/15
```

#### 6 Sensitivity analysis for the theoretical model

```
h <- 5000 #number of parameter sets
# random seed for replicability
set.seed(1)
# latin hypercube of size h, number of parameters
lhs <- maximinLHS(h, 10)

# parameters: a, q, d, pred1, pred2, parm1, parm2, y, w,
# sigma
a_default <- parms_focal[1] #host birth rate
q_default <- parms_focal[2] #host crowding coefficient
d_default <- parms_focal[3] #host background death rate
pred1_default <- parms_focal[4] #lower predation
pred2_default <- parms_focal[5] #upper predation
parm1_default <- parms_focal[6] #multiplier of virulence-transmission trade-
    off
parm2_default <- parms_focal[7] #exponentially relates virulence-transmission
     trade-off
y_default <- parms_focal[8] #recovery rate
w_default <- parms_focal[9] #immunity waning rate
sigma_default <- parms_focal[10] #superinfection rate

min_mult <- 0.5
max_mult <- 1.5
# minimum is half that of the default value
a_min <- a_default * min_mult
q_min <- q_default * min_mult
d_min <- d_default * min_mult
pred1_min <- pred1_default * min_mult
pred2_min <- pred2_default * min_mult
parm1_min <- parm1_default * min_mult
parm2_min <- parm2_default * min_mult
y_min <- y_default * min_mult
w_min <- w_default * min_mult
sigma_min <- sigma_default * min_mult

# maximum is one and a half times that of the default value
a_max <- a_default * max_mult
q_max <- q_default * max_mult
d_max <- d_default * max_mult
pred1_max <- pred1_default * max_mult
pred2_max <- pred2_default * max_mult
parm1_max <- parm1_default * max_mult
parm2_max <- parm2_default * max_mult
y_max <- y_default * max_mult
w_max <- w_default * max_mult
sigma_max <- sigma_default * max_mult

# Rescaling Latin Hypercube Sample
params.set <- cbind(a = lhs[, 1] * (a_max - a_min) + a_min, q = lhs[,
     2] * (q_max - q_min) + q_min, d = lhs[, 3] * (d_max - d_min) +
     d_min, pred1 = lhs[, 4] * (pred1_max - pred1_min) + pred1_min,
     pred2 = lhs[, 5] * (pred2_max - pred2_min) + pred2_min, parm1 = lhs[,
```

```
    6] * (parm1_max - parm1_min) + parm1_min, parm2 = lhs[,
    7] * (parm2_max - parm2_min) + parm2_min, y = lhs[, 8] *
    (y_max - y_min) + y_min, w = lhs[, 9] * (w_max - w_min) +
    w_min, sigma = lhs[, 10] * (sigma_max - sigma_min) +
    sigma_min)

labels = c("a", "q", "d", "pred1", "pred2", "parm1", "parm2",
  "y", "w", "sigma")
```

#### 6.1 Calculate model outputs across parameter values

```
# counter
j <- 1

# how big the data frame to hold the parameter values and
# results should be number of parameters (10) + number of
# results/dependent variables examined (in this case, 20 so
# 10 + 20 = 30)

results_length <- 33

# create dataframe to hold results where each column is
# either an input or output of the model and each row is a
# different run of the model based on the lhs parameter sets
data <- data.frame(matrix(rep(NA, h * results_length), nrow = h))

# for each parameter set...
for (i in 1:h) {

  # grab the model inputs
  data[i, 1:10] <- params <- as.list(c(params.set[i, ]))

  # create variable to catch errors
  skip_to_next <- F

  # if source code cannot find coCSS/throws and error,
  # skip_to_next is set to TRUE and loop goes on to the next
  # parameter set
  result = tryCatch(outputs_and_paths(c(params.set[i, ])),
    error = function(e) {
      skip_to_next <<- TRUE
    })

  if (skip_to_next) {
    # update counter when skipping to next loop because of error
    j <- j + 1
    next
  }

  # if able to find coCSS, record model outputs
  data[j, 11] <- result[1]  #prev1 - prevalence of parasite under the lower
    predation
```

```
    data[j, 12] <- result[2]  #prev2 - prevalence of parasite under the higher
        predation
    data[j, 13] <- result[3]  #prev_ratio - ratio of prevalence under low to
       high predation
    data[j, 14] <- result[4]  #H1 - host density under lower predation
    data[j, 15] <- result[5]  #H2 - host density under higher predation
    data[j, 16] <- result[6]  #H_ratio - ratio of host densities
    data[j, 17] <- result[7]  #v_int1 - virulence at coCSS under lower
       predation
    data[j, 18] <- result[8]  #v_int2 - virulence at coCSS under high
       predation
    data[j, 19] <- result[9]  #v_ratio - ratio of virulences
    data[j, 20] <- result[10]  #c_int1 - shoaling at lower predation
    data[j, 21] <- result[11]  #c_int2 - shoaling at higher predation
    data[j, 22] <- result[12]  #c_ratio - ratio of shoaling
    data[j, 23] <- result[13]  #mort1 - overall host mortality under lower
       predation
    data[j, 24] <- result[14]  #mort2 - overall host mortality under high
       perdation
    data[j, 25] <- result[15]  #mort_ratio - ratio of overall mortality
    data[j, 26] <- result[16]  #mort_change_par - change in mortality due to
       parasites
    data[j, 27] <- result[17]  #trans1 - transmission under lower predation
    data[j, 28] <- result[18]  #trans2 - transmission under upper predation
    data[j, 29] <- result[19]  #trans_ratio - ratio of transmission
    data[j, 30] <- result[20]  #pathway 1
    data[j, 31] <- result[21]  #pathway 2
    data[j, 32] <- result[22]  #pathway 3
    data[j, 33] <- result[23]  #pathway 4
    print(j)
    j <- j + 1
}

# in case there are runs that threw errors, remove them
data <- na.omit(data)
```

##### 6.2 Compute correlation coefficients and intervals

```
# response of mortality change due to parasites to all
# parameters calculated in the sixteenth output -
# mort_change_par
bonferroni.alpha <- 0.05/10

out_list <- list()
for (i in 11:33) {
    out_list[[i]] <- pcc(data[, 1:10], data[, i], nboot = 1000,
        rank = TRUE, conf = 1 - bonferroni.alpha)
    print(i)
}

summary_list <- list()
for (i in 11:33) {
```

```
    summary_list[[i]] <- print(out_list[[i]])
}

# Results for a given output are in each row Results for a
# given parameter are in each column First layer is min ci,
# second is value, third is max ci
parameter_effects <- array(NA, dim = c(23, 10, 3))
for (i in 11:33) {
    parameter_effects[i - 10, , 1] <- summary_list[[i]]$`min. c.i.`
    parameter_effects[i - 10, , 2] <- summary_list[[i]]$original
    parameter_effects[i - 10, , 3] <- summary_list[[i]]$`max. c.i.`
}

positive_effects <- list()
negative_effects <- list()
for (i in 11:33) {
    positive_effects[[i]] <- which(parameter_effects[i - 10,
        , 1] > 0)
    negative_effects[[i]] <- which(parameter_effects[i - 10,
        , 3] < 0)
}

# Can see which parameters had a positive or negative effect
# on each outcome with commands like the following:
# labels[positive_effects[[11]]]

# Need to sum things a bit for the proportion of virulence
# increase via shoaling (i.e., non-consumptive effects of
# predators) the change in morality due to parasites and the
# change in virulence due to sociality can be greater than 1
# if other parameters decrease mortality or virulence to make
# displaying them more intuitive we convert any number above
# 1 to 1. This is equivalent to saying 'of the increase in x,
# how much is factor y responsible for?'
mcdp <- data[, 26]
mcdp[which(mcdp > 1)] <- 1

# Classify possible outcomes for vcds, the percent of
# virulence increase that is via increased shoaling rate
for (i in 1:dim(data)[1]) {
    temp_num = data[i, 33] + data[i, 32]
    temp_denom = data[i, 30] + data[i, 31] + data[i, 32] + data[i,
        33]
    if (temp_num < 0) {
        vcds[i] <- 0
    }
    if (temp_num > 0 & temp_denom > 0) {
        vcds[i] <- min(c(temp_num/temp_denom, 1))
    }
    if (temp_num > 0 & temp_denom < 0) {
        vcds[i] <- 1
    }
}
```
